# Centralized brain networks underlie body part coordination during grooming

**DOI:** 10.1101/2024.12.17.628844

**Authors:** Pembe Gizem Özdil, Jonathan Arreguit, Clara Scherrer, Auke Ijspeert, Pavan Ramdya

## Abstract

Animals must coordinate multiple body parts to perform important tasks such as grooming, or locomotion. How this movement synchronization is achieved by the nervous system remains largely unknown. Here, we uncover the neural basis of body part coordination during goal-directed antennal grooming in the fly, *Drosophila melanogaster*. We find that unilateral or bilateral grooming of one or both antenna, respectively, arises from synchronized movements of the head, antennae, and forelegs. Simulated replay of these body part kinematics in a biomechanical model shows that this coordination makes grooming more efficient by permitting unobstructed, forceful collisions between the foreleg tibiae and antennae. Movements of one body part do not require proprioceptive sensory feedback from the others: neither amputation of the forelegs or antennae, nor immobilization of the head prevented movements of the other unperturbed body parts. By constructing a comprehensive antennal grooming network from the fly brain connectome, we find that centralized interneurons and shared premotor neurons interconnect and thus likely synchronize neck, antennal, and foreleg motor networks. A simulated activation screen of neurons in this network reveals cell classes required for the coordination of antennal movements during unilateral grooming. These cells form two coupled circuit motifs that enable robust body part synchronization: a recurrent excitatory subnetwork that promotes contralateral antennal pitch and broadcast inhibition that suppresses ipsilateral antennal pitch. Similarly centralized controllers may enable the flexible co-recruitment of multiple body parts to subserve a variety of behaviors.

## Introduction

Complex animal behaviors rely upon the adept coordination of multiple body parts. For example, walking requires synchronized movements of each limb to efficiently move the body through space^1,2^. This coordination requires the co-activation of multiple, distinct motor networks (those for each moving leg) as well as the suppression of other networks (those for stabilizing the other legs in stance). Thus, body part coordination depends critically upon effective communication between neuronal populations controlling each appendage^3^.

The organization of interlimb and intersegmental networks has been most extensively studied the context of vertebrate locomotion^4–9^. In rodents, inhibitory *V*_0_ commissural interneurons in the spinal cord regulate left-right alternation, while excitatory *V*_0_ neurons mediate left-right synchrony in a speed-dependent manner^10,11^. Such commissural interneurons have been identified in swimming and walking across species ^7,9,12–14^, implying that these coordination mechanisms are evolutionarily conserved. Similarly, intersegmental interneurons have been described for insect locomotor coordination^1,15^. These advances highlight that our understanding of body part coordination remains largely limited to the identification of key cell types rather than the elucidation of systems-level network architectures and circuit mechanisms.

The adult fly, *Drosophila melanogaster*, is an ideal experimental model for gaining both a more comprehensive and deep understanding of motor control. Flies generate numerous behaviors that require movement synchronization^16–18^. In addition, the fly’s brain and motor system—the ventral nerve cord (VNC)—have been fully mapped^19–25^. This enables the detailed analysis of circuit connectivity. Finally, extensive libraries of transgenic driver lines make it possible to genetically target and manipulate specific neuronal subtypes^26,27^.

Here, we investigated goal-directed antennal grooming in the fly to obtain a multi-level mechanistic understanding of body part coordination. Grooming is an ethologically important, evolutionarily conserved behavior comprised of precisely targeted limb movements to remove debris or parasites from the body^28^ and is performed by both mammals and insects^29–33^. Adult flies groom many different body parts—their antennae, eyes, proboscis, legs, wings, and abdomen—following a prioritization sequence that is governed by a suppression hierarchy^18,34–36^. Optogenetic neural activation experiments in *Drosophila* have identified key neurons responsible for grooming including peripheral sensory neurons^37–39^, brain interneurons^40^, descending neurons projecting from the brain to downstream VNC motor networks^40–42^, and interneurons within the VNC^43,44^ which may contribute to central pattern generation for limb control^45^. Nevertheless, the organizational logic of grooming kinematics and underlying motor networks remains largely unknown.

Numerous tools and resources now allow us to overcome this gap. First, pose estimation software enables high-throughput 3D measurements of body kinematics^46,47^. Second, these kinematic data can be replayed in a biomechanical model of the fly to infer contact forces^48–50^. Third, the brain and VNC connectomes can be used to simulate network dynamics^49,51,52^. Here, we combine these tools and resources to uncover kinematic and neural mechanisms for body part coordination during antennal grooming. Flies principally perform two subtypes of grooming, unilateral or bilateral, for cleaning one or both antennae, respectively. These are distinguished by their differential synchronization of head, antennae, and foreleg movements. Simulated replay of these kinematics in a biomechanical model shows that coordination increases grooming efficiency by preventing obstructions and enabling forceful foreleg-antennal collisions. Fixing the head in place or removing the antennae or forelegs, does not disrupt synchronization, revealing that proprioceptive sensory feedback is not required. Indeed, the fly brain connectome reveals that centralized and shared premotor interneurons bind motor modules for these body parts. Finally, simulated activation and silencing of neurons in the antennal grooming network identifies coupled recurrent excitatory and broadcast inhibition circuit motifs that enable robust body part coordination.

## Results

### Antennal grooming arises from coordinated movements of the head, antennae, and forelegs

To precisely quantify antennal grooming, we developed an experimental system that allows us to measure head, antennal, and foreleg kinematics in tethered flies **(Fig. 1A)**. We reliably elicited antennal grooming through bilateral optogenetic stimulation of antennal Johnston’s Organ F (‘JO-F’) neurons^40^ (*aJO-GAL4-1> CsChrimson*; **Extended Data Fig. 1A**), or by presenting both antennae with a brief puff of air.

**Fig 1.**
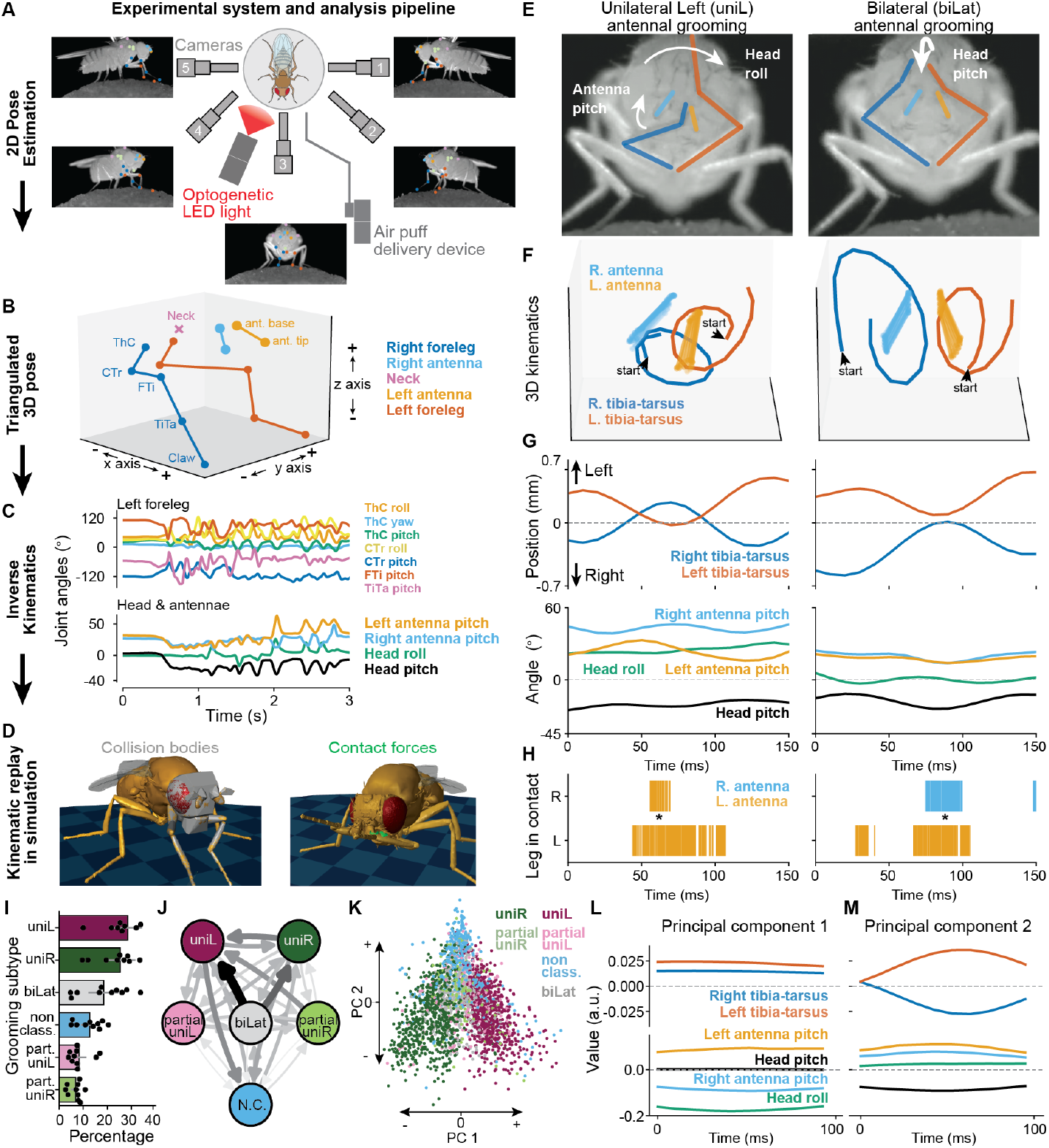
Kinematic analysis reveals two major subtypes of *Drosophila* antennal grooming. **(A)** Schematic of the experimental system used to optogenetically-elicit and record antennal grooming (not to scale). The behavior of a tethered fly on a spherical treadmill is captured by five cameras with different view angles. Video recordings are then used to measure 2D poses. A 617 nm LED light is used to activate CsChrimson expressed in antennal Johnston’s Organs. A separate device can deliver air puffs to the fly’s antennae. **(B)** Tracked 2D keypoints from each camera are then triangulated to reconstruct 3D poses. Shown is a schematic of the 3D pose coordinate system used to track kinematics, where the x-axis is anteroposterior, the y-axis is mediolateral, and the z-axis is dorsoventral. Arrows specify the positive and negative directions along each axis. Body parts are color-coded. **(C)** From the 3D poses, we use inverse kinematics to calculate joint angles for the neck, antennae, and forelegs. **(D)** Joint angles are used to control joint actuators in NeuroMechFly, a physics-based simulation of the adult fly. Collision bodies **(left)** can be used to quantify the contact forces **(right)** between the antennae and forelegs. **(E)** Front camera images overlaid with color-coded ‘bones’ of the legs (blue/right, orange/left) and antennae (light blue/right, light orange/left). Illustrated are two antennal grooming subtypes: unilateral left (‘uniL’) and bilateral (‘biLat’) across panels E-H. Head and antennal movements are schematized (white arrows). **(F)** Visualization of sample 3D kinematic trajectories of the base and tip of the antennae as well as the tibia-tarsus joints of the forelegs during antennal grooming. Joints are color-coded as in panel E. **(G, top)** Mediolateral positions of the tibia-tarsus leg joints. Positive values represent the left side in fly-centric coordinates. Joints are color-coded as in panel E. **(G, bottom)** Head and antennal degree of freedom angles. For head roll, positive values are leftward. For head pitch, negative values are downward. For antennal pitch, positive values are upward. **(H)** Contact diagrams inferred from collisions between the foreleg segments and the antennae. This was derived using kinematic replay of joint angles in NeuroMechFly. Asterisks mark the occurrence of each corresponding antennal grooming subtype. **(I)** Percentage of time spent performing each class of optogenetically-elicited antennal grooming. Each circle represents a biological replicate (n=10, N=33). Error bars show mean and 95% confidence intervals. **(J)** Antennal grooming classes visualized in a graph network where each arrow represents transitions from one class to another. Darker and thicker arrows represent a higher frequency of state transitions. Color coding as in panel I. **(K)** Reduced dimensionality representation of antennal grooming kinematics. Each dot represents a 100 ms epoch of 3D positions of antennal key points and foreleg tibia-tarsus joints (only along the y and z axes); head roll&pitch, antennal pitch, and some leg joint angles (i.e., ThC roll, pitch; CTr roll, pitch). Epochs are color-coded by antennal grooming class as in panels I-K. **(L-M)** Representations of joint kinematics along the **(L)** first and **(M)** second principal components which describe 27.5% and 13.9% of the variance, respectively. Values are in arbitrary units. Color code is the same as in panel G.

We recorded animal behavior simultaneously from five camera viewpoints^46^ and then used these videos to track 2D positions of keypoints on the antennae, neck, and forelegs^53^. These positions were then triangulated in 3D^47^ **(Fig. 1B)** and then, via sequential inverse kinematics^54^, used to compute joint angles **(Fig. 1C; Supplementary Video 1)**. In addition to providing quantitative measurements of grooming movements, these joint angles could be replayed in NeuroMechFly^48–50^, a biomechanical model of the fly **(Fig. 1D; Supplementary Video 2)**, to infer contacts and forces between body parts that are otherwise challenging to measure experimentally.

Visual inspection of our behavioral videos revealed that optogenetically-elicited antennal grooming tends to fall into two subtypes with distinct body part kinematics: (i) unilateral grooming of either the right (‘uniR’) or left (‘uniL’) antenna by both forelegs, or (ii) bilateral (‘biLat’) grooming in which each foreleg simultaneously grooms its ipsilateral antenna **(Fig. 1E; Supplementary Video 3)**. Importantly, both subtypes were also observed in response to air-puffs **(Supplementary Video 4)**, with quantitatively similar head, antenna, and foreleg kinematics **(Extended Data Fig. 2)**.

Unilateral grooming in response to optogenetic **(Fig. 1F; Extended Data Fig. 1B)** or air-puff **(Extended Data Fig. 1C)** stimulation is characterized by several kinematic features. First, the forelegs move laterally toward the targeted antenna and produce cyclical, synchronized leg sweeps **(Fig. 1G, top)**. Second, the non-targeted antenna is pitched upwards around the mediolateral axis, possibly to avoid collisions with the legs **(Fig. 1G, bottom)**. Third, the head is pitched down and rolled to the side, bringing the targeted antenna into the task space of the forelegs **(Fig. 1G, bottom)**. By contrast, during bilateral grooming, the antennae do not appear to move. As well, the head does not rotate but is instead pitched downwards, lowering both antennae to the work space of the forelegs. Indeed, simulated replay of these kinematics in our biomechanical model confirmed that, during unilateral grooming, collisions occur between both forelegs and the targeted antenna whereas, during bilateral grooming, each foreleg principally collides with its ipsilateral antenna **(Fig. 1H)**. Thus, the kinematics of the head, antennae, and forelegs are differentially correlated during unilateral versus bilateral antennal grooming **(Extended Data Fig. 1D)**.

We quantified the frequency of grooming subtypes by manually classifying behaviors across multiple flies (n=10 animals) during optogenetic stimulation **(Fig. 1I)**. Unilateral and bilateral grooming are the most frequent, occurring in more than 70% of behavioral events. The remaining (*<*30%) behaviors could not clearly be defined as antennal grooming (e.g., leg lifting) and thus were labeled unclassified (‘non-class’). In other more rare instances, behaviors only partially matched unilateral coordination (‘partial uni’). We often observed that flies transitioned smoothly between different grooming subtypes. The most frequent transitions occurred from bilateral to unilateral grooming **(Fig. 1J; Extended Data Fig. 1E)**. Importantly, our classification of antennal grooming into unilateral and bilateral subtypes was also observed using an unbiased, dimensionality reduction approach. Principal component analysis (PCA) performed on the same head and foreleg kinematics data revealed marked subdivisions along the first two principal components which explained over 40% of the variance (**Extended Data Fig. 1F**). Specifically, unilateral grooming subtypes reside on either side of a central space filled by bilateral and non-classified subtypes **(Fig. 1K)**. Strikingly, features observed during each grooming subtype were evident in time-series data from these first two principal components. The first principal component resembles unilateral grooming: the left and right tibia-tarsus joints are positioned laterally on one side of the midline, the left and right antennae are pitched in opposite directions, and there is a low degree of head pitch when head roll angles are larger (**Fig. 1L**). Consistent with this, uniR and uniL are found on opposite (negative versus positive) sides of this first principal component (**Fig. 1K**). Kinematics in the second principal component were reminiscent of bilateral grooming: the tibia-tarsus joints are symmetrically on opposite sides of the midline, antennal pitch angles are similar, and although head roll is nearly zero, head pitch is large (**Fig. 1M**). Indeed, bilateral and non-classified grooming are distributed along this second principal component axis (**Fig. 1K**).

The synchronization of body part movements during antennal grooming can also be quantified as a systematic correlation of their kinematics over time **(Extended Data Fig. 1D)**. To rule out the possibility that these correlations trivially arise from the displacement of the head and antennae by forceful contact with the forelegs, we optogenetically elicited antennal grooming in animals with their forelegs amputated. There we observed similar head/antennal kinematics reflected in overlapping spatial occupancies **(Extended Data Fig. 3)**. Thus, the head, antennae, and forelegs appear to be actively coordinated.

### Body part coordination improves grooming efficiency

Having observed stereotypically synchronized head, antennal, and foreleg movements during antennal grooming, we next asked to what extent this coordination increases grooming efficiency by facilitating contacts between the forelegs and antennae. For example, we hypothesized that during unilateral grooming of the left antenna: (i) leftward roll of the head might bring the left antenna closer to the forelegs, (ii) upward pitch of the non-targeted right antenna might prevent contact with forelegs, and (iii) leftward shift of the forelegs might facilitate contact with the targeted left antenna.

Testing these hypotheses experimentally would require measuring foreleg-antennal contacts while perturbing single degrees of freedom (e.g., by eliminating head pitch without affecting head roll). Such experiments are currently not technically feasible—we lack both a means of measuring body part contacts as well as the ability to genetically perturb motor neurons driving individual antennal and neck degrees of freedom. Therefore, we performed perturbation experiments in NeuroMechFly^48–50^. Specifically, we replayed real, recorded body part kinematics in our simulation while measuring collisions and forces between the antennae and forelegs. We repeated this experiment while systematically modulating the amplitude of individual degrees of freedom—forward head pitch, sideways head roll, or upward pitch of the non-targeted antenna.

We first investigated the importance of downward head pitch during bilateral grooming. We replayed measured kinematics and quantified antenna-leg collisions from real data (‘*gain* = 1’, **Fig. 2A, top**), or while virtually fixing the head in its rest position (‘*gain* = 0’, **Fig. 2A, bottom**). Compared with our real data (**Fig. 2B, top**), when the head was fixed in place we observed that the leg segments in contact with the antennae shifted from the tibiae to the more distal tarsi (**Fig. 2B, bottom; Supplementary Video 5, left)**. By systematically performing this experiment using kinematic data from multiple animals with substantial head pitch (median greater than 14 degrees) but minimal head roll (**Extended Data Fig. 4A**) we confirmed that larger head pitch results in increased tibia-antenna contact (**Fig. 2C, left**) and decreased tarsus-antenna contact (**Fig. 2C, right**). Thus, downward head pitch during bilateral antennal grooming may serve to maximize contact between the fly’s foreleg tibia and antennae. Why might flies prioritize tibial contact with the antennae? One possibility is that the tibiae may exert more force on the antenna compared with the more compliant tarsi—a thinner multi-segmented structure with numerous passive joints. Consistent with this, even though our simulated tarsi are less compliant than real tarsi, they nevertheless exert less force on the antennae, on average, than the tibiae do (**Extended Data Fig. 4B,C**).

**Fig 2.**
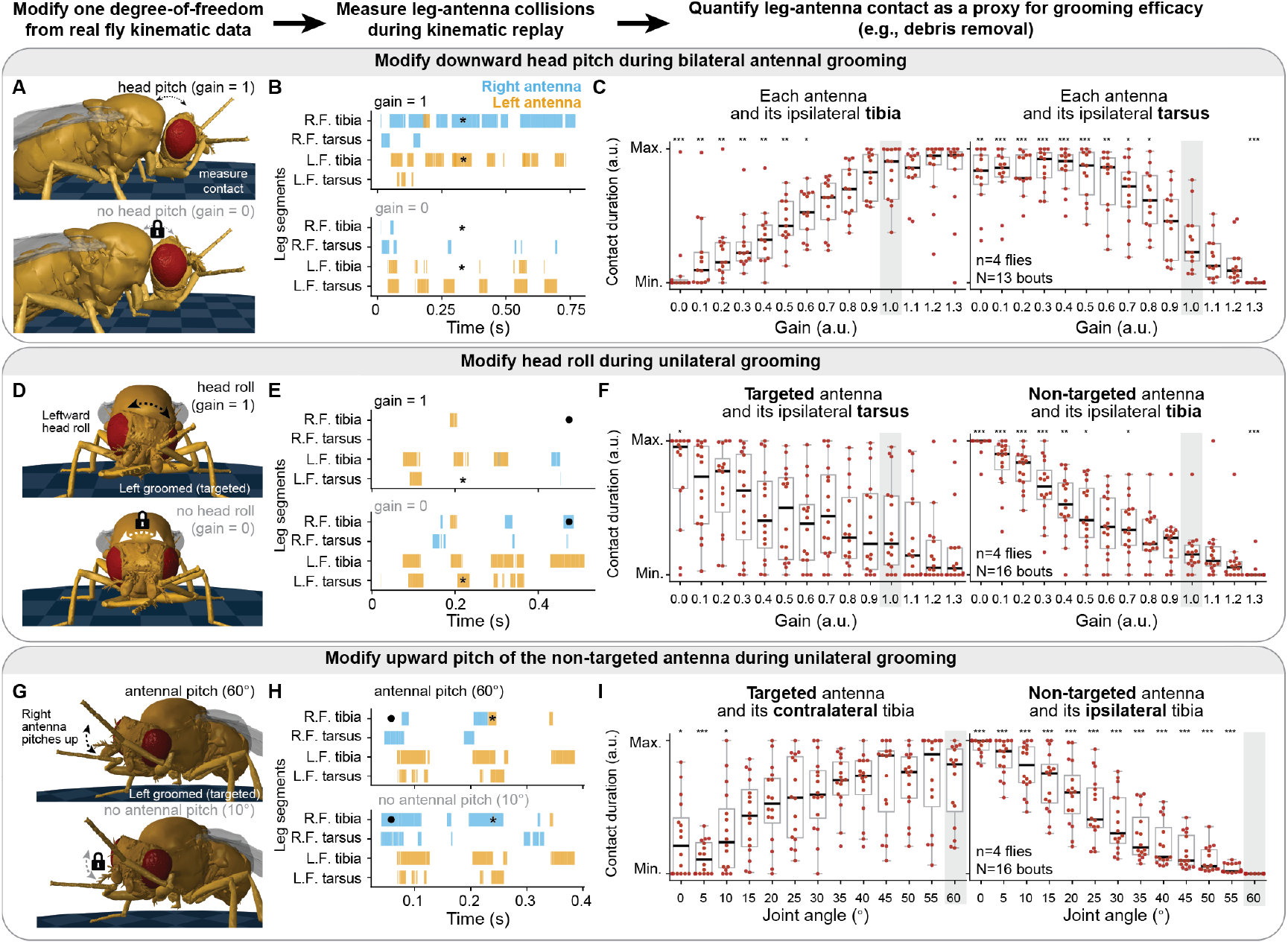
Kinematic replay in a biomechanical model reveals the contribution of head and antennal movements to foreleg-antennal interactions. We generated a kinematics dataset to be replayed in simulation, allowing us to gradually perturb individual joint degrees of freedom while measuring contacts between the forelegs and antennae. **(A, D, G)** Snapshots from kinematic replay simulations with either an intact (top), or perturbed (bottom) **(A)** head pitch, **(D)** head roll, or **(G)** antennal pitch. **(B, E, H)** Collision diagrams between tibia and tarsus foreleg segments and both antennae (right/blue, left/orange) either in intact (top), or perturbed (bottom) **(B)** head pitch, **(E)** head roll, or **(H)** antennal pitch. In panel B, asterisks indicate where tibial collisions disappear as head pitch is decreased. In panel E, asterisks indicate that when head roll decreases, there are increased collisions between the targeted antenna (left/orange) and the ipsilateral, left tarsus. As well, circles indicate increased collisions between the non-targeted antenna (right/blue) and the ipsilateral, right tibia. In panel H, as antennal pitch decreases asterisks indicate reduced collisions between the targeted antenna (left/orange) and its contralateral, right tibia. As well, circles show increased collision between the non-targeted antenna (right/blue) and its ipsilateral, right tibia. **(C, F, I)** Contact duration between specific antennal and foreleg segments as a function of the movement magnitude of a joint degree of freedom. Contact is normalized between minimum and maximum values across all gains/magnitudes for each trial. Data are presented for: **(C)** n=4 flies, N=13 bilateral grooming bouts; **(F, I)** n=4 flies, N=16 unilateral grooming (uniL and uniR combined) bouts. Intact, unmodified kinematics are highlighted in light gray boxes. Box plots show the median and quartiles. Box plot whiskers extend to 1.5 times the interquartile range (IQR). Shown are statistical results for a two-sided Mann–Whitney U test comparing the intact distribution with other gains/magnitudes: ***: *P <* 0.001, **: *P <* 0.01, *: *P <* 0.05 and not significant (NS): *P* ≥ 0.05. P values were corrected using the Simes–Hochberg procedure.

During unilateral grooming, flies roll their heads to the side, lowering the targeted antenna. Similar to head pitch during bilateral grooming, we hypothesized that this head roll might bring the targeted antenna into the task space of the legs, while positioning the non-targeted antenna further away. To test this, we replayed unilateral grooming in our simulation while modulating the amplitude of head roll. Indeed, collision diagrams show that during, for example, unilateral left antennal grooming, compared with intact head roll (gain=1, orange epochs **Fig. 2D-E, top**) when head roll is suppressed, there is increased contact between the right leg and the non-targeted, right antenna (gain=0, blue periods **Fig. 2D-E, bottom; Supplementary Video 5, middle**). Using kinematic data from multiple flies during unilateral grooming with appreciable head roll (median more than 8 degrees) (**Extended Data Fig. 4D**), we confirmed that suppressing head roll results in (i) a shift from contact with the ipsilateral tibia to the more distal tarsus (**Fig. 2F, left**) as well as (ii) an increase in collisions between the non-targeted antenna and its ipsilateral tibia (**Fig. 2F, right**). Thus, head roll appears to bring the targeted antenna toward and the non-targeted antenna away from the task space of the foreleg tibiae.

Finally, we asked whether upward pitch of the non-targeted antenna facilitates unilateral grooming by allowing the fly to avoid undesired leg collisions. Because in our real experiments antennal poses were often obstructed during leg-antenna interactions, direct replay of real antennal joint angles was not possible. Therefore, we instead set the antennal pitch degree of freedom to a constant value ranging from 0^°^ to 60^°^ in increments of 5^°^—a range of angles that resembles those measured from real flies (**Extended Data Fig. 4D**). We found that when the non-targeted antenna was pitched upward (angle 60^°^) both tibiae principally contact the targeted antenna (orange, **Fig. 2G-H, top**). However, when the non-targeted antenna remains in its resting position (angle 10^°^) it obstructs the ipsilateral tibia, reducing contact with the targeted antenna (**Fig. 2G-H, bottom; Supplementary Video 5, right**). This was consistent across multiple animals and grooming epochs: suppressing upward pitch of the non-targeted antenna reduces grooming of the targeted antenna by the contralateral tibia (**Fig. 2I, left**) due to increased collisions with the non-targeted antenna (**Fig. 2I, right**).

Thus, head and antennal movements during grooming appear to optimize tibial contact with the targeted antenna(e) by (i) bringing the targeted antenna into the foreleg task space via downward head pitch or sideways head roll and (ii) preventing collisions between the legs and non-targeted antenna via sideways head roll and upward pitch of the non-targeted antenna. Next, we sought to decipher the neural mechanisms underlying this tripartite coordination of body parts during unilateral antennal grooming.

### Multi-body part synchronization does not rely on proprioceptive feed-back

The synchronous activation of motor networks for the head, antenna, and forelegs during unilateral grooming can arise from several potential control frameworks. First, in a ‘sensory feedback’ framework, movements of one body part (e.g., the head) may generate proprioceptive signals that initiate and/or maintain motor programs for the other two body parts (e.g., the antenna and forelegs) (**Fig. 3A**). Within this framework, we can envisage three means of yielding tripartite coordination of the head, antennae, and legs: (i) proprioceptive feedback from moving one body part could drive movements of a second whose proprioceptive feedback would in turn drive a third (‘cascading coordination’), (ii) proprioceptive feedback from two moving body parts may both be needed to drive movements of a third (‘additive coordination’), or (iii) proprioceptive feedback from one moving body part may drive movements of the other two (‘diverging coordination’) (**Extended Data Fig. 5**). In an alternative framework, proprioceptive feedback-independent or ‘open-loop’ mechanisms might underlie synchronous movements of the head, antennae, and legs (**Fig. 3B**). Open-loop models can be classified based on the origin of movement synchronization in the brain’s sensorimotor pathway: it may arise at the sensory layer of JO neurons (‘input shared’), via an ensemble of central neurons (‘central hub’), or as a consequence of intercommunicating motor modules (‘output shared’).

**Fig 3.**
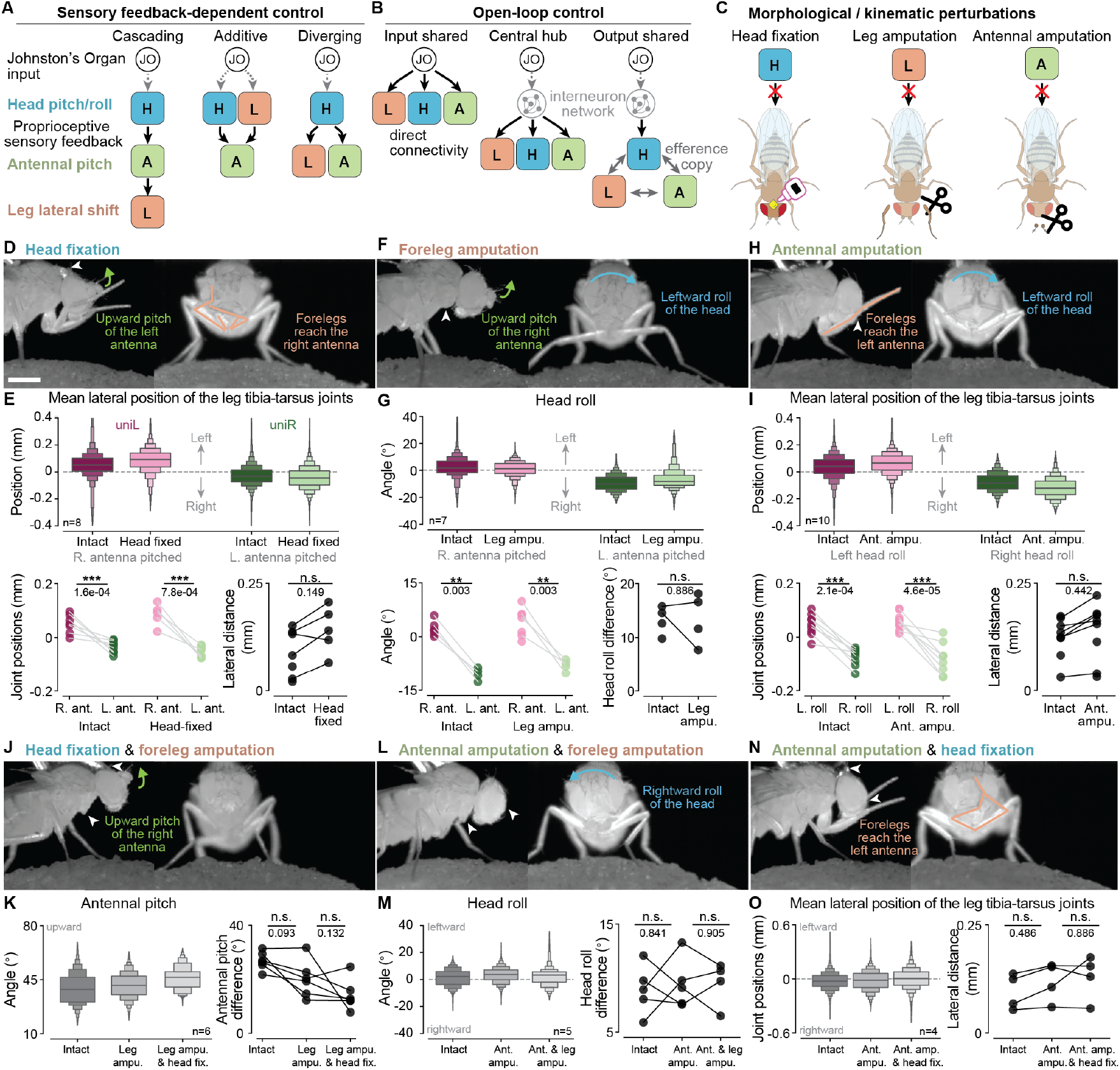
Experimental perturbations show that sensory feedback is not essential for body part coordination. **(A)** Proposed control models that depend upon proprioceptive sensory feedback. Each colored block represents a motor module consisting of motor neurons and their premotor partners controlling a particular body part. For each model only one of several possible configurations is shown. In *cascading coordination*, proprioceptive sensory feedback from the first moving body part drives movements of the following body parts. In *additive coordination*, feedback from the first two moving body parts jointly drive movements of the third. In *diverging coordination*, feedback from one body part drives the movements of the other two. **(B)** Alternatively, ‘open-loop’ control models do not depend upon proprioceptive sensory feedback. Body part coordination can be driven at different levels along the sensorimotor pathway, beginning from immediately downstream of JO sensory input (‘Input shared’), to central interneurons (‘Central hub’), and finally using efference copy from the motor modules themselves (‘Output shared’). **(C)** Morphological and kinematic perturbations used to test the contribution of sensory feedback to antennal grooming. These include amputating the forelegs and/or antennae, as well as fixing the head in place with UV-curable glue. Each perturbation blocks one arrow in the sensory feedback-dependent control diagrams. **(D, F, H, J, L, N)** Front and side camera images overlaid with line drawings of the legs (orange), and arrows indicating movements of an antenna (green), and/or head rotations (blue). The locations of experimental perturbation(s) are indicated (white arrowheads). **(E, G, I) (top)** Distribution of **(E**,**I)** tibia-tarsus joints’ mean lateral positions and **(G)** head roll during unilateral left (magenta) or unilateral right (green) antennal grooming in either intact (darker color) or experimental (lighter color) animals. **(bottom-left)** Median values of each kinematic variable across trials for each fly and grooming subtype. **(bottom-right)** Differences between uniL and uniR kinematic variables for intact versus experimental conditions. For **(E)** n=8, **(G)** n=7, and **(I)** n=10 animals. **(K, M, O) (left)** Distribution of joint angles and positions for the remaining freely moving body part in experiments perturbing two body parts at once. **(right)** Differences between the 90^th^ and 10^th^ percentile of **(K)** the pitched antenna’s joint angles, and **(M, O)** median differences between body part movements to the left and right. For **(K)** n=6, **(M)** n=5, and **(O)** n=4 animals. In boxen plots, the median is represented by the largest middle line. Each successive level outward contains half of the remaining data. In scatter plots, each dot represents an individual fly, with lines connecting the same fly across behavioral subtypes or experimental conditions. One-sided Mann–Whitney U tests compare uniR versus uniL under the same conditions, while two-sided tests compare data across experimental conditions (e.g., intact versus head-fixed). Significance levels are indicated as follows: ***: *P <* 0.001, **: *P <* 0.01, *: *P <* 0.05 and not significant (NS): *P* ≥ 0.05.

We first aimed to distinguish between proprioceptive sensory feedback versus open-loop control frameworks. To do so, we measured antennal grooming in flies both before and after body part manipulations intended to eliminate proprioceptive sensory feedback: foreleg amputation, antennal amputation, and/or head fixation (**Fig. 3C**). Additionally, to test additive feedback models we simultaneously perturbed two body parts (e.g., amputating the forelegs and immobilizing the head). In total, we tested six perturbations: (i) fixation of the head (**Fig. 3D**), (ii) amputation of the forelegs (**Fig. 3F**), (iii) amputation of the antennae (**Fig. 3H**), (iv) head fixation and foreleg amputation (**Fig. 3J**), (v) antennal and foreleg amputation (**Fig. 3L**), and (vi) antennal amputation and head fixation (**Fig. 3N**). To quantify the impact of perturbing one body part, we investigated the kinematics of the remaining two intact body parts. For example, we examined which antenna the forelegs reach laterally towards while the fly pitches one antenna upward (**Fig. 3E, top**). We observed that flies preserve their leg and antenna coordination pattern following head immobilization (**Fig. 3E, bottom; Supplementary Video 6**). Although we measured minor changes in foreleg trajectories, particularly in the proximal leg joints, (**Extended Data Fig. 6**), this is likely because flies have more room to move when the head is fixed and not pitched downward. Similarly, amputation of the forelegs did not alter the relationship between head roll and antennal pitch during unilateral grooming (**Fig. 3G; Supplementary Video 7**). Finally, after antennal amputation, we observed that the lateral position of the forelegs still tracked the direction of head rotation (**Fig. 3I; Supplementary Video 8**).

Next, we perturbed two body parts simultaneously and measured the movement range of the remaining body part. In foreleg amputated and head-fixed flies without significant neck and leg proprioceptive sensory feedback, we found that flies still actively lift their antenna (**Fig. 3K; Supplementary Video 9**). We note that in intact flies the forelegs push the antenna closer to the head, reducing antennal pitch angles. As well, after both foreleg and antennal amputations, we observed that the head still rolls in both directions (**Fig. 3M; Supplementary Video 10**). Finally, amputating the antennae and fixing the head in place did not disrupt lateral movements of the forelegs (**Fig. 3O; Supplementary Video 11**).

Because no perturbation significantly altered the coordinated movements of intact body parts, we conclude that proprioceptive sensory feedback is not required for head, antennae, and leg movement synchronization during antennal grooming. Other, open-loop control mechanisms are thus more likely at play.

### A centralized brain network links multiple motor modules

To evaluate potential open-loop control models for body part synchronization, we extended our ‘input’, ‘central’, and ‘output’ models to include real neuronal subtypes including sensory inputs, interneurons, and motor modules (i.e., premotor neurons and their target motor neurons moving a specific body part). In this extended ‘open-loop’ framework we could envision at least four different neural network architectures that might enable the synchronization of head, antenna, and foreleg movements: via (i) shared antennal Johnston’s Organ sensory input (‘input shared’), (ii)common input from central interneurons controlling premotor-motor modules (‘central hub’), (iii)coupling between premotor circuits for each body part (‘premotor coupling’), or (iv) shared premotor circuits for multiple body parts (‘shared premotor’) (**Fig. 4A**).

**Fig 4.**
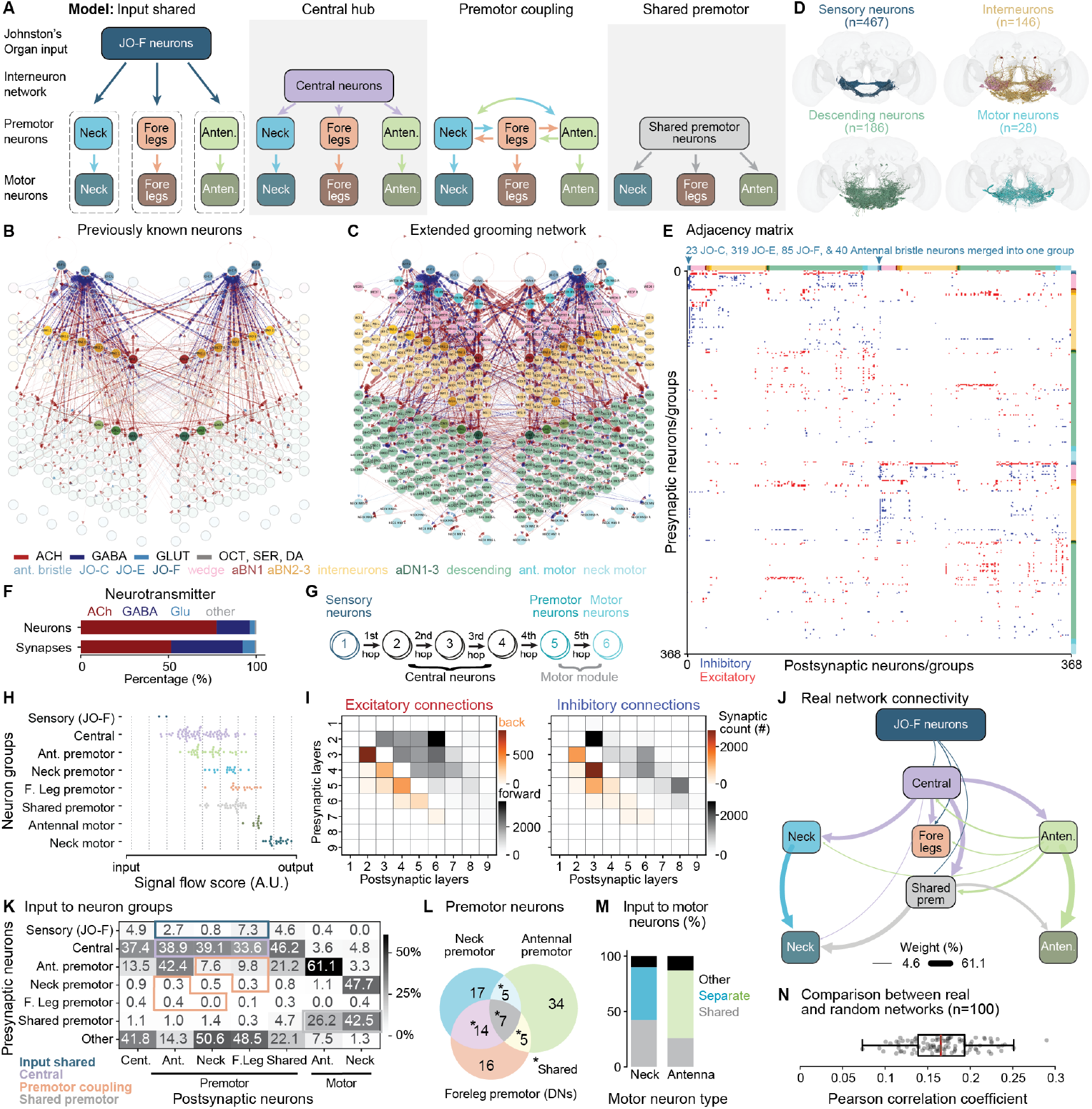
Body part motor modules are linked by central circuits in the fly brain connectome. **(A)** Schematized network models for open-loop motor coordination of the head, forelegs, and antennae. In the *input shared* model, JO-F sensory neurons directly project onto all three motor modules consisting of premotor and motor neurons. In the *central hub* model, a group of central neurons diverge onto all three motor modules. In the *premotor coupling* model, each motor modules communicates via their distinct yet interconnected premotor neurons. In the *shared premotor* model, all three sets of motor neurons are controlled by shared premotor neurons. **(B)** Graph visualization of the connectivity of previously identified^40^ antennal grooming neurons (highlighted) in the fly brain connectome^19–21^. Neuron types are color-coded the same across panels B-E. **(C)** Graph visualization of our more comprehensive antennal grooming network constructed using the fly brain connectome and seeded from the network in panel B. Arrows indicate pre- to postsynaptic connectivity. Line colors and widths indicate neurotransmitter identities and synaptic weights, respectively. **(D)** Renderings of all sensory, interneuron, descending, or motor neurons in the antennal grooming network. **(E)** Adjacency matrix of the constructed network, ordered by neuron type as in panel C. The connectivity matrix was binarized, making excitatory connections +1 and inhibitory connections -1. **(F)** The frequency of different neurotransmitters across neurons (top) and synaptic connections (bottom) in the network. **(G)** The flow of signals across five hops in the connectome-derived grooming network. Premotor neurons are defined as being directly upstream and projecting more than 5% of their outputs onto motor neurons. Central neurons are defined as situated between JO-F sensory neurons and motor modules (premotor and motor neurons) within a maximum of four hops. **(H)** Neuronal groups ordered by their signal flow scores, ranging from input-like (left) to output-like (right). Each dot represents one neuron, with JO-F neurons merged into one group for each side of the brain. The axis was divided into nine intervals, and neurons were assigned to their respective layers. **(I)** Heatmaps showing (left) excitatory and (right) inhibitory connectivity between layers. Indicated are the degree of feedback (orange) versus feedforward (black) connectivity. **(J)** Real connectivity diagram of the network (to be compared with those in panel A). Line widths are proportional to the percentage of connections between neuron groups (real values are given in panel K). Connections below 4.6% are not shown. The same color code is used across panels J-M. **(K)** Heatmap showing the contribution of inputs from one neuron group to another. Expected connections for each hypothetical model are outlined. **(L)** Venn diagram showing the number of neurons classified as being premotor to neck (blue), antennal (green), or leg (orange) motor neurons. Also indicated are shared premotor neurons that synapse upon more than one type of motor neuron (asterisks). **(M)** Relative contributions of inputs from premotor neuron types to motor neuron groups. “Separate” refers to premotor neurons that project onto only one motor neuron type, and “shared” refers to those projecting onto more than one motor neuron type. **(N)** Pearson correlation coefficients comparing connectivity diagrams (as in panel J) derived from the real adjacency matrix with those from randomly shuffled adjacency matrices. Each dot represents a shuffled network constructed using a random seed. Box plots show the median (red line) and quartiles. Whiskers extend to the full distribution, excluding outliers beyond 1.5 times the interquartile range (IQR).

To investigate the degree to which these network architectures might underlie open-loop coordination, we used the adult female whole-brain connectome^19–21,55^ to construct a comprehensive network of antennal grooming-related neurons. We began with neurons that had previously been described as involved in antennal grooming. These included sensory neurons like the antennal Johnston’s Organ (‘JO’ C-E-F)^37,40^ and mechanosensory bristles^39^, brain interneurons (aBN1,2,3), and descending neurons (aDN1,2,3)^40^ (**Fig. 4B**). To these we added antennal and neck motor neurons, enabling us to define the motor modules for these body parts. Then, we systematically incorporated neurons monosynaptically connected to any of these seed neurons (**Extended Data Fig. 7A**) with synaptic connections to the seed network exceeding a threshold defined by a parameter sweep (**Extended Data Fig. 7B**), and informed by previous work^20^. This threshold excluded extraneous neurons with little information flow to or from antennal grooming neurons while still retaining a broad range of neuron types (**Extended Data Fig. 7C**).

Our final antennal grooming network consists of 827 neurons with sparse connectivity (2195 connected neuron pairs or 0.3% sparsity) (**Fig. 4C-E**). Of these connections ∼ 31% are contralateral across brain hemispheres. Although ∼ 77% of neurons are excitatory, they contribute only ∼ 55% of synapses (**Fig. 4F**). Thus, on average, inhibitory neurons contribute proportionally more synapses to this network, consistent with previous findings^56^. Additionally, we observed high degree distributions among inhibitory interneurons (**Extended Data Fig. 7D**, circled in black), and the excitatory aBN1, suggesting that these neurons may influence network dynamics on a global scale.

To test the relative match to our different open-loop control models (**Fig. 4A**), we next categorized interneurons as being either central or premotor. Neurons were defined as ‘central’ if they lay on the path (on average with more than 5% of synaptic inputs) from ‘JO-F’ sensory inputs to motor neurons within five hops^56^ (**Fig. 4G**). From this group of ‘central neurons’ we then reclassified as ‘premotor neurons’ those with at least 5% of their outputs directly targeting motor neurons controlling the antennae or neck (**Fig. 4G**).

Ultimately, our approach classified neurons in our antennal grooming network into four major groups: sensory (JO-F), central, premotor (antennal, neck, foreleg, or shared), and motor (antennal or neck). We used a signal flow sorting algorithm^57^ to measure the extent to which information flows in a feedforward manner in this network. This algorithm scores each node in the graph based on its proximity to the input and output. Signal flow scores across nodes in our neuron groups (**Extended Data Fig. 7E**), exhibited a clear gradient in which, as expected, sensory JO-F neurons were situated closest to the input, followed by central, premotor, and finally motor neurons near the output (**Fig. 4H**). We next asked to what degree neurons form feedback connections to preceding layers. Specifically, we divided the signal flow axis into nine layers to get sufficiently many (∼ 40) neurons per layer. Then we examined the connectivity between neurons in each layer by summing the number of synapses made between each neuron pair. For both excitatory and inhibitory connections, we found that the grooming network is predominantly feedforward (gray), with some feedback (orange) connections enriched near sensory layers 2 and 3 (**Fig. 4I**). We speculate that this feedback might reflect presynaptic inhibition upon sensory inputs^58^.

Close examination of our network’s connectivity matrix appears to immediately exclude two open-loop models (**Fig. 4J-K**; **Extended Data Fig. 8**). First, JO-F neurons connect only minimally to premotor and motor neurons. Therefore, sensory input does not appear to directly drive synchrony across motor modules (**Fig. 4A**, ‘input shared’ model). Second, premotor modules do not appear to be connected strongly to one another (**Fig. 4A**, ‘premotor coupling’ model). By contrast, we observe strong connectivity between central and premotor neurons (**Fig. 4J-K**) whereby individual central neurons project onto shared premotor or multiple categories of premotor neurons (**Extended Data Fig. 7F**). This finding supports the ‘central hub’ model. Similarly, there exist common premotor neurons which target multiple groups of motor neurons (**Fig. 4J-K**), consistent with our ‘shared premotor’ model. However, shared premotor neurons contribute only ∼ 42% and ∼ 26% of synapses to neck and antennal motor neurons, respectively (**Fig. 4L-M**). As well, in the VNC some shared premotor neurons project to both neck and foreleg motor neurons (**Extended Data Fig. 7G**). Interestingly, within the VNC, the axons of descending neuron arising from the brain represent the largest fraction of shared (rather than foreleg-or neck-specific) premotor neurons (**Extended Data Fig. 7H**). This suggests that brain networks may also be principally responsible for directly coordinating leg and neck movements within the VNC.

In sum, our findings indicate that central interneurons, with a smaller contribution from shared premotor circuits, are best positioned to coordinate antennal, neck, and leg motor modules. Correlation analyses with randomized adjacency matrices confirm that the real network’s configuration is highly non-random (**Fig. 4N**). More specifically, comparing the real connectome network with randomized versions of this network (**Extended Data Fig. 9A-B**) shows that the proportion of connections in the ‘central’ and ‘shared premotor’ models is significantly greater than expected by chance (**Extended Data Fig. 9C**, purple and gray boxes). As well, connections associated with the ‘input-shared’ and ‘premotor coupling’ models are significantly lower or not statistically different than that expected by chance aside from antennal premotor to foreleg premotor connectivity (**Extended Data Fig. 9C**, dark blue and orange boxes).

### Simulating a connectome-derived antennal grooming network

Connectivity analysis revealed that central neurons likely coordinate the activity of head, antennae, and foreleg motor modules. However, static connectivity information alone is insufficient to understand the contributions of individual neurons and circuit motifs to behavioral dynamics. For example, instead of forming a continuous gradient of behavioral subtypes, behavioral responses tended to be either unilateral, or bilateral. This pattern suggests that the antennal grooming network may operate using winner-take-all action selection, a process whose study requires investigating the temporal evolution of neural activity. Therefore, to explore how our network might drive this selection process, we simulated its dynamics.

Specifically, we built a connectome-derived artificial neural network, in which each neuron is modeled as a leaky integrator^59^ (see Methods). We optimized network parameters to generate outputs that matched a training dataset consisting of head and antennal kinematics from flies (n=10 animals) whose JO-F neurons were stimulated with diverse optogenetic patterns including steps of varying duration, and pulses of varying frequency (**Fig. 5A, left**). To keep sensory input well-defined, these animals’ forelegs were amputated (**Fig. 5A, middle**). This allowed us to (i) prevent leg-antennal contact during grooming and thereby limit mechanosensory feedback from the head^39^ and forelegs^60^, as well as (ii) reduce the importance of ascending leg proprioceptive sensory feedback^61^. Importantly, our previous amputation experiments (**Fig. 3**) demonstrated that head and antennal coordination can occur even in the absence of the forelegs. From these video data we computed head and antennal kinematics (**Fig. 5A, right; Extended Data Fig. 10A**) as target outputs for our network to replicate during simulations.

**Fig 5.**
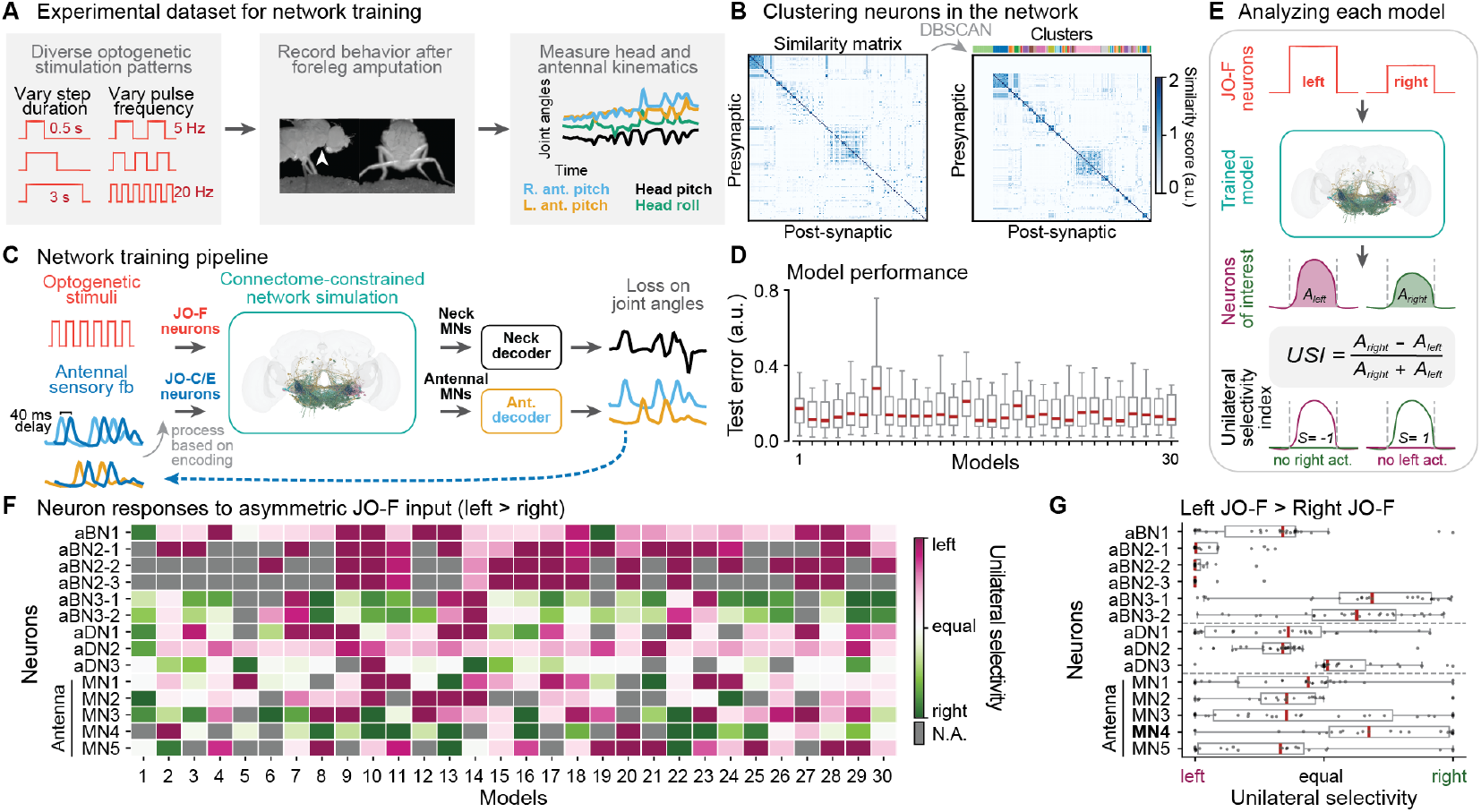
Training and evaluating connectome-derived artificial neural networks for antennal grooming. **(A)** The training dataset includes head and antennal kinematics of flies in response to optogenetic stimulation consisting of varying step durations and pulse frequencies. Flies had their forelegs amputated to prevent confounding contacts between the forelegs and the head or antennae. **(B)** An unsupervised clustering algorithm, DBSCAN, was used to cluster neurons based on the connectivity in one hemisphere of the symmetrized network. These clusters served as a proxy for cell types in the network. It was applied to a similarity matrix of the grooming network (left) restricted to one brain hemisphere and excluding sensory and motor neurons. Clusters are color-coded. Neurons that were not clustered (left-most green cluster) were assigned to their own group. **(C)** Virtual optogenetic stimuli (red) were delivered to JO-F neurons in the connectome-constrained neural network. Readouts from antennal and neck motor neurons were fed into separate decoders, representing the musculoskeletal properties of the neck and antennae. The decoders output one-dimensional joint angles for antennal pitch (right/blue, left/orange) and head pitch (black). The left and right antennal motor neuron activities were fed to the same decoder separately. The mechanosensory inputs JO-C and JO-E receive antennal sensory feedback: a processed and time-delayed copy of antennal joint angles. Model parameters were optimized to match real kinematic measurements from panel A. The loss was evaluated on a held-out test dataset, unseen during training. **(D)** Training was performed for 30 different random seeds and models. **(E)** Trained models were analyzed by applying slightly asymmetric JO-F activation and quantifying the unilaterality of neural responses for each neuron pair. The unilateral selectivity index (USI) metric is defined as the area under the right neuron’s response curve minus that of the left neuron’s response, divided by the sum of both responses. Only positive neural responses were considered in this calculation. For instance, the metric equals one when there is a positive response in the right neuron but no response in the left neuron. The metric is undefined (N.A.) when both neurons do not respond. **(F)** Responses of antennal brain interneurons (aBNs), descending neurons (aDNs), and motor neurons (aMNs) to asymmetric JO-F input (left*>*right) in every trained model. Neural responses were quantified using the USI metric. Grey squares indicate zero neural activity in both neurons (USI = 0/0). Magenta and green squares represent neurons with larger ipsilateral and contralateral responses, respectively. The darkest colors denote cases in which neurons on one side are predominantly active. **(G)** Summary of unilateral selectivity of aBNs, aDNs, and aMNs across models. Among antennal motor neurons, only antennal motor neuron 4 (aMN4) consistently exhibits a contralateral response to asymmetric (left greater than right) JO-F input. Each dot represents a model (a square in panel F). **(D, G)** Box plots show the median (red line) and quartiles. Whiskers extend to the full distribution, excluding outliers beyond 1.5 times the interquartile range (IQR).

Due to imperfections in connectome data acquisition and reconstruction as well as real biological variation, there are differences in connectivity across the left and right brain hemispheres^20,62^ (see **Extended Data Fig. 12** and Discussion). This might introduce spurious, artifactual asymmetries in network simulations. Therefore, to minimize the inductive biases stemming from this asymmetry, we made our network bilaterally symmetric. We symmetrized the adjacency matrix by setting the synaptic values for each connection to the maximum among each bilateral pair of neurons (**Extended Data Fig. 10B**; see **Extended Data Fig. 10C** to compare results using different methods). Consequently, each paired neuron has bilaterally-symmetric pre- and post-synaptic neighbors as well as an equal number of synapses in both hemispheres. Having prepared our network in this way, we next trained it to reproduce measured antennal and head kinematics in response to virtual optogenetic stimulation of JO-F neurons (**Fig. 5C**). The mechanosensory JO-C/E neurons also received a fictive sensory feedback: antennal kinematics with a sensorimotor delay of 40 ms^63,64^. We read out motor neuron activities from five pairs of antennal and four pairs of neck pitch motor neurons in the brain^65^ but excluded neck roll motor neurons because they have not been identified in the brain connectome. Motor neuron activities were then fed into two separate decoders, encapsulating antennal and neck musculoskeletal systems, which output fictive antennal and head pitch joint angles (**Fig. 5C**). As in previous connectome-constrained modeling work^51^, the edges of this network and each neuron’s neurotransmitter identity were fixed as they are in the brain connectome^19–21,66^. However, neuronal parameters including the membrane time constants, resting potentials, synaptic strengths, and decoder parameters were optimized via backpropagation through time (BPTT)^67^ to match our training dataset. We performed training across thirty random seeds and confirmed convergence in all cases to small loss values (**Fig. 5D**). We next analyzed neural dynamics in our trained models. To focus on the winner-take-all aspect of unilateral grooming, we presented slightly asymmetric JO-F input (left antenna input slightly exceeding the right) and examined which neurons are driven to purely right or left activation. We devised a metric, the unilateral selectivity index (USI), for each bilateral neuron or cluster pair by measuring the area under the response curves for the left and right hemispheres, and then computing their difference as a fraction of the total area (**Fig. 5E**). Thus, a USI of one indicates fully right-dominant activity (contralateral to the more stimulated left antenna), while a USI of negative one indicates fully left-dominant activity (ipsilateral to the more stimulated left antenna). Intuitively, a USI of one is analogous to unilateral left grooming in which the right (non-targeted) antenna lifts in response to stimulation of the left (targeted) antenna to avoid collisions with the forelegs (**Fig. 1E, left**). We applied this metric to key antennal grooming neurons (e.g., aBNs, aDNs, and aMNs) with asymmetric JO-F input (left *>* right) and observed consensus across thirty models on the responses of each neuron class (**Fig. 5F**). Among the five motor neurons, only aMN4 consistently exhibited a contralateral response (**Fig. 5G; Extended Data Fig. 10D** for an exemplary model), suggesting that aMN4 may drive upward antennal pitch of the non-targeted antenna during unilateral grooming.

### Coupled circuit motifs enable robust unilateral coordination

Having generated a connectome-derived model of the antennal grooming network, we next set out to identify circuit motifs that may underlie body part coordination during unilateral grooming. We focused our analysis on antennal pitch coordination: upward pitch of the contralateral, non-targeted antenna and quiescence of the targeted antenna. We studied antennal movements for several reasons. First, upward pitch of the non-targeted antenna is a hallmark of unilateral grooming that is synchronous with head roll and lateral foreleg movements (**Fig. 1E**). Second, unlike neck and leg motor neurons, antennal motor neurons are located exclusively in the fully mapped brain. Third, these motor neurons are a compact and tractable system for analysis: only five motor neurons control four muscles in each antenna^68^, compared with the numerous neurons and muscles controlling the neck^65^ and forelegs^24^.

The precise roles of individual antennal motor neurons have not been fully established. There-fore, because in six models aMN4 neurons consistently exhibited contralateral responses to asymmetric JO-F input (models 10, 11, 13, 16, 22, and 23; dark green rectangles in the aMN4 row in Fig. **5**F), we used the activities of aMN4 motor neurons in these models as a readout. In combination with neural perturbations, this readout allowed us to identify neurons and circuits that encourage exclusively upward pitch of the non-targeted, contralateral antenna in response to bilaterally asymmetric JO-F input. Indeed, aMN4 activity closely reflected this action selection process; when systematically testing a range of left-right JO-F input current pairs, we found that even slight input asymmetries nevertheless result in fully unilateral aMN4 responses (**Fig. 6A**). To investigate the neural mechanisms underlying this winner-take-all response, we focused on three models with the most biologically relevant characteristics: fully unilateral aMN4 activity during slightly asymmetric JO-F input as well as no aMN4 activity during bilaterally symmetric JO-F input (akin to no antennal pitch during bilateral grooming) (**Fig. 6B–C;** see **Extended Data Fig. 11A-B** for the other three models).

**Fig 6.**
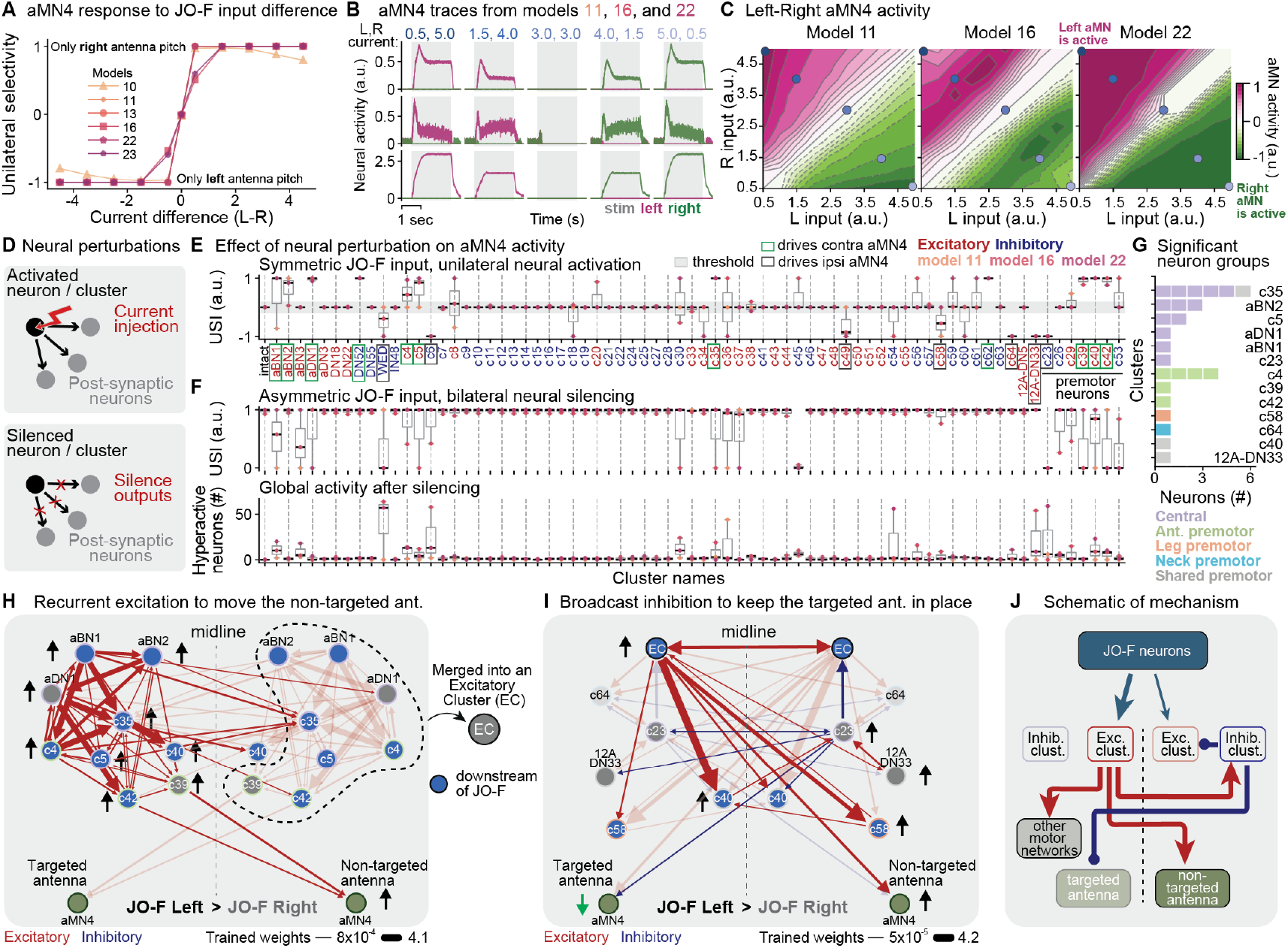
Simulated neural activation uncovers neurons and circuit motifs driving unilateral coordination. **(A)** aMN4 responses quantified by the USI metric for different JO-F input current pairs (5.0, 0.5; 4.5, 1.0; 4.0, 1.5; 3.5, 2.0; 3.0, 3.0; as well as their mirrored values). Six trained models are shown. USI values of 0 indicate no bias or equal response, whereas -1 and 1 correspond to fully left and fully right aMN4 responses, respectively. **(B)** Left (magenta) and right (green) aMN4 neural activity traces in response to JO-F input pairs marked in panel C, across models 11, 16, and 22. JO-F stimulation period is indicated (gray shaded region). During asymmetric JO-F input, the aMN4 contralateral to the stronger input responds, whereas the other aMN4 does not. Note that voltage traces are processed through a ReLU activation function, thus there may be subthreshold, negative responses. **(C)** Activities of aMN4 on the left or right side of the brain. These are shown as a function of the input current magnitudes to left and right JO-F in the intact network. Values represent the difference between the area under the curve for the left and right aMN responses (magenta for left aMN-dominant, green for right aMN-dominant). Solid contours mark positive value intervals, and dashed contours mark negative intervals, in increments of 0.1. Neither antennal motor neuron dominates along or near the diagonal (white). Circles indicate the five pairs of current values shown in panel B, using the same color code. **(D)** Neural perturbations used to assess the contribution of neurons/cluster to driving unilateral coordination: (i) bilaterally symmetric JO-F input during unilateral activation of left-hemisphere neurons/clusters (top) and (ii) slightly asymmetric JO-F input (left *>* right) during bilateral silencing of neurons/clusters (bottom). Perturbations were systematically applied to each cluster/neuron in the network. **(E)** Effects of unilateral neural activation on aMN4 responses (USI metric). Bilaterally symmetric JO-F input drives equal left and right aMN4 responses in the unperturbed, intact network (far-left column). Neurons whose unilateral activation transform this into contralateral right aMN4 responses are outlined in green; those driving ipsilateral left aMN4 responses are outlined in gray. Each dot represents a model. The median thresholds of 0.1 (contralateral) and -0.1 (ipsilateral) are highlighted (gray horizontal bar). Red and blue labels indicate excitatory and inhibitory neurons/clusters, respectively. **(F)** Effects of bilateral neural silencing on aMN4 responses (top) and global network activity (bottom). USI was calculated for responses during slightly asymmetric JO-F input (left *>* right). The intact, unperturbed response (far-left column) is fully right-dominant (USI = 1). Global activity quantifies the number of neurons in the perturbed network with activity five times greater than their mean activity in the intact, unperturbed network. Each dot represents a model. **(E, F)** Box plots show the median and quartiles. Whiskers extend to the full distribution, excluding outliers beyond 1.5 times the interquartile range (IQR). **(G)** The number and type of neurons for each significant cluster shown in panels H and I. Neuron types are color-coded. **(H)** Diagram illustrating the recurrent excitation motif driving contralateral aMN4 activation in panel E (green boxed neurons/clusters). This recurrent excitatory motif was then merged into a single cluster (excitatory cluster or ‘EC’). **(I)** Diagram illustrating connections between the EC (self connections of EC are not shown) and the neurons eliciting ipsilateral aMN4 activation in panel E (gray boxed neurons/clusters). In this broadcast inhibition motif, the inhibitory neuron c23 (right) prevents upward pitch of its contralateral antenna (left) by suppressing its contralateral aMN4. Cluster c64 is dimmed because it is inactive in the intact network. **(H, I)** Neurons or clusters with higher activity compared to their contralateral counterparts are marked with upward black arrows, while those with lower activity are indicated with downward green arrows. Connections from neurons with lower activity are made transparent for visualization purposes. Neurons/clusters directly downstream of JO-F are shown in blue, and edge colors correspond to neuron groups as in panel G. Red and blue lines denote excitatory and inhibitory connections, respectively, with line thicknesses proportional to normalized weights after the training of model 22. **(J)** Schematic representation of the mechanism underlying unilateral coordination via aMN4 activation. JO-F neurons activate excitatory clusters on the targeted antenna’s side (thicker arrow from JO-F), which activates aMN4 pitch motor neurons of the non-targeted antenna and other network modules. Simultaneously, inhibitory neurons on the non-targeted antenna’s hemisphere suppress excitation of the targeted antenna’s motor neurons and its excitatory clusters, preventing its upward pitch. Red and blue lines indicate excitation and inhibition, respectively. Less active elements are dimmed.

We reasoned that central circuits promoting unilateral pitch might be identified by their ability to drive asymmetric network activity in the presence of equal JO-F sensory input to both antennae. Therefore, in a first neural activation screen, we provided bilaterally symmetric JO-F input and simultaneously activated individual neurons/clusters in the left hemisphere (**Fig. 6D, top**). In a second, complementary neural silencing screen, we presented asymmetric JO-F input (left *>* right) to drive unilateral aMN4 responses. Simultaneously we systematically silenced bilateral pairs of neurons/clusters to identify those necessary for driving the selection of unilateral antennal pitch (i.e., unilateral aMN4 responses) **(Fig. 6D, bottom**).

Our neural activation screen uncovered eighteen neurons/clusters whose unilateral activation could drive aMN4 activity, during bilaterally symmetric JO-F stimulation. These produced either higher contralateral (**Fig. 6E, green outlines**: aBN1,2, aDN1, DN52, c4, c5, c35, c62, c39, c40, c42), or ipsilateral (**Fig. 6E, gray outlines**: WED, c6, c49, c58, 12A-DN33, c23) responses. Notably, when perturbed in the neural silencing screen, not all of these neurons/clusters had an impact (**Fig. 6F**). This may be due to redundancy in the network or inactivity during JO-F stimulation in the unperturbed network. It is also worth noting that, across all seeds, silencing the inhibitory neurons/clusters WED and c6 also more globally amplified network activity **(Supplementary Video 12)** (**Fig. 6F, bottom**).

We observed that, although our primary focus was on antennal motor control, activation screen hits were not exclusively antennal premotor neurons/clusters (see **Extended Data Fig. 11C** for all aMN4 premotor neurons) and showed similar neural responses to JO-F stimuli across seeds (**Extended Data Fig. 11D**). We found numerous hits that could be categorized as central, shared premotor, neck premotor, and leg premotor (**Fig. 6G**). Thus, we hypothesized that hits may contribute to circuits performing motor coordination more broadly. To test this hypothesis, we bundled two groups of activation hits based on whether they tipped the balance towards driving contralateral or ipsilateral aMN4 activity. Remarkably, this simple bundling yielded several well-connected circuit motifs (**Fig. 6H-I**).

The first motif consists of a circuit dominated by recurrent excitation between aBN1, aBN2, aDN1, c4, c5, c35, c39, c40, and c42 (**Fig. 6H**). The majority of these neurons/clusters are directly downstream of JO-F neurons (**Fig. 6H**, nodes in blue). Thus, we envision that this circuit may amplify small biases in JO-F input. As well, models predict that activating any neuron/cluster within this circuit may robustly recruit the majority of the network, drive the activity of premotor neurons c39 and c42, and elicit a fully unilateral response from contralateral aMN4s (**Extended Data Fig. 11E**) to drive upward pitch of the non-targeted antenna. This group also includes two inhibitory neurons: DN52 and c62 (**Fig. 6E**). We did not include them in the motif because they are normally inactive during JO-F stimulation (**Supplementary Video 13**). However, they may be uncovered in the activation screen as encouraging contralateral aMN4 activation because they suppress their contralateral excitatory cluster, c40, which is involved in ipsilateral aMN4 activation. Therefore, rather than directly exciting the contralateral aMN4 and pitching the non-targeted antenna, DN52 and c62 indirectly inhibit the ipsilateral aMN4 and movements of the targeted antenna (**Extended Data Fig. 11F**).

Next we focused on neurons/clusters in the unilateral activation screen which drove responses in the ipsilateral aMN4. Among the seven neurons/clusters identified, two inhibitory clusters, WED and c6, reduce ipsilateral JO-F activity via presynaptic inhibition (**Extended Data Fig. 11G**). Their activation creates input asymmetry by suppressing sensory input from ipsilateral JO-F neurons (**Extended Data Fig. 11H**). However, the remaining five neurons/clusters appear to engage the inhibitory neuron, c23 (‘asteriod’^69^). c23 inhibits its ipsilateral recurrent excitatory circuit (‘EC’) and c40. The latter cluster acts as a bridge between the two motifs in that it receives strong excitatory input from EC and, in turn, excites the contralateral c23 (**Fig. 6I**). Additionally, c23 directly inhibits the targeted antenna’s aMN4. Interestingly, c23 neurons across the brain also reciprocally inhibit one another, a competitive inhibition motif commonly associated with decision-making and action selection^70–77^. Finally, other neurons in this motif include the leg premotor cluster c58, the neck premotor cluster c64—which is normally inactive during JO-F stimulation (**Supplementary Video 13**)—and the shared premotor cluster 12A-DN33. These neurons receive excitatory input from the contralateral EC and activate their ipsilateral c23 either directly or indirectly (**Fig. 6I**), highlighting the central role of c23-based broadcast inhibition.

Thus, our connectome-derived network neural activation screen has uncovered two interconnected motifs that likely mediate winner-take-all unilateral antennal pitch in response to symmetric or only slightly asymmetric JO-F stimulation: (i) a recurrent excitatory circuit (EC) that encourages and maximizes contralateral aMN4 activity and non-targeted antennal pitch, as well as (ii) EC/c40-based activation of the contralateral broadcast inhibitor c23 which suppresses the contralateral EC and movements of the targeted antenna. Taken together, these findings illustrate how both excitatory and inhibitory motifs can be combined to more robustly drive network activity into one of two discrete unilateral grooming states (**Fig. 6J**).

## Discussion

Here, we have combined behavioral quantification and perturbations, biomechanical simulations, connectome analysis, and connectome-derived artificial neural network simulations to investigate how the adult fly, *Drosophila melanogaster*, synchronizes head, antennae, and foreleg movements during antennal grooming. We found that this tripartite coordination does not rely on proprioceptive sensory feedback from individual body parts but appears to be driven by a centralized network of interneurons and shared premotor neurons. Embedded within this network, we discovered coupled recurrent excitation and broadcast inhibition circuit motifs which drive the unilateral selection to pitch upward the non-targeted antenna while suppressing similar movements of the targeted/groomed antenna.

### The utility of coordinating multiple body parts during grooming

Why might it be beneficial for the fly to coordinate head, antennae, and forelegs movements while grooming its antenna? Simulated kinematic replay in a biomechanical model suggests that this body part synchronization facilitates unobstructed and more forceful tibial rather than tarsal contact with the antennae. In line with this, we have observed that flies often retract their antennae towards their head during bilateral grooming, possibly to increase the stiffness of the scape-pedicel joint and stabilize the antenna. In addition, we speculate that brushing one antenna with both forelegs may be more effective in removing debris because it contacts areas that the ipsilateral leg alone cannot reach. It also allows for greater forces to be applied to the antenna. Complex hair-like structures on the tibial segments may also act as a brush to improve debris cleaning and, thus, improve olfactory sensing ^78,79^. Finally, because the neuromuscular system controlling the tibia is more complex than the tarsal control system^24,80^, this strategy maximizing tibia-antenna contact may benefit from more precise leg positioning.

### Proprioceptive sensory feedback is not required for body part coordination

A longstanding question in motor control has been the extent to which body part coordination arises from sensory feedback versus feedforward centralized control^1,5^. In some cases, movements are primarily driven by sensory feedback^81,82^, while in others centrally generated motor patterns remain intact even without input from leg mechanosensors^83,84^. In walking flies, mechanosensory feedback does not contribute strongly to interleg coordination but is important for precise foot placement^85,86^. However, unlike locomotion in which the legs are mechanically coupled to one another through the substrate, there is no such mechanical coupling between the head, antennae, and legs during grooming. This might suggest that accurate grooming must rely on ongoing proprioceptive feedback to precisely position the body parts with respect to one another. Surprisingly, we found that *Drosophila* antennal grooming does not require proprioceptive feedback to initiate body part coordination. This is consistent with previous studies of head grooming in other insects^87^. We speculate that the unimportance of proprioceptive feedback during grooming may be acceptable because imperfect coordination does not pose an existential threat. As a result, a simpler centralized control strategy may eliminate the computational and energy costs associated with continuously processing sensory feedback.

### Centralized networks may enable flexible coordination

Within the open-loop grooming control framework, we observed that motor modules are primarily interconnected by central interneurons, rather than by inputs (i.e., JO sensory neurons) or outputs (i.e., premotor neurons). This configuration may best balance the needs for robust yet flexible coordination. We speculate that if motor modules were all directly targeted by JO sensory inputs they might be able to generate fast and reliable coordinated movements—something that would be desired for an escape response. However, this configuration would impede the independent and flexible control of individual body parts because the control signal stems from a single shared source. Furthermore, any input noise or perturbation would directly propagate to downstream motor networks. Similarly, if motor modules were connected near the output layer we might observe slower but similarly inflexible coordination: the movements of multiple body parts would be inextricably yoked together. Therefore, coupling motor modules at a central layer (i) offers multiple entry points to drive grooming (e.g., JO or bristle stimulation), (ii) allows behaviors to be more readily gated by internal state, and (iii) still enables the independent control of constituent body parts for different purposes (e.g., head pitch for gaze stabilization). We speculate that in this way centralized coordination may simplify the evolution of new behaviors through the coupling or uncoupling of motor modules.

This centralized coordination mechanism may be conserved across species in different contexts. Rodents also self-groom using similar kinematics including cyclical forelimb movements and downward head pitch^29^. In rats, the brainstem is both necessary and sufficient to execute a complete sequence of self-grooming ^88,89^. Most of our antennal grooming network is located in the fly’s gnathal ganglia, a brain region that has been compared to the vertebrate brainstem^90^.

### Inductive bias in the brain connectome and idiosyncratic behavior

Previous studies have shown that fruit flies exhibit individual preferences in walking handedness^91,92^ as well as olfactory^93^ and phototactic^94^ decision-making. Recent modeling work has suggested that slight variations in synaptic connectivity might account for these idiosyncratic behaviors^95,96^. Notably, structural asymmetries exist even among fully reconstructed and proofread neurons in the fly brain connectome^19,20^. We hypothesize that these asymmetries might explain why we observe some flies consistently initiating unilateral grooming of the same antenna across trials, even during bilaterally symmetric optogenetic stimulation.

To investigate whether structural asymmetries in the connectome could drive grooming preferences in response to bilaterally symmetric JO-F input, we trained the original, non-symmetrized network. Indeed, the original network exhibited a strong and consistent bias toward unilaterally activating key antennal grooming neurons despite bilaterally symmetric JO-F stimulation **(Extended Data Fig. 12A, left)**. For example, among the antennal motor neurons, aMN5 consistently showed rightward selectivity, whereas aMN1, aMN2, and aMN4 exhibited a bias to the left **(Extended Data Fig. 12B)**. This bias disappeared when the network was symmetrized **(Extended Data Fig. 12A, right)**. Although experimental variability could partially be attributed to genetic factors, such as differences in CsChrimson expression levels, our findings support the possibility that asymmetries in brain connectivity may contribute to idiosyncratic behavioral preferences.

## Methods

### Data acquisition

#### Fly husbandry and stocks

All experiments were performed on female adult *Drosophila melanogaster* raised at 25 ^°^C and 50% humidity on a 12 hr light-dark cycle. Two days (36-40h) before optogenetic experiments, we transferred experimental flies to a vial containing food covered with 40 µl of all-trans-retinal (ATR) solution (100 mM ATR in 100% ethanol; Sigma Aldrich R2500, Merck, Germany) and wrapped in aluminium foil to limit light exposure. Experiments were performed on flies 3-5 days post-eclosion (dpe) between Zeitgeber Times (ZT) 4-10. Genotype used and sources are indicated in Table 1.

**Table 1.**
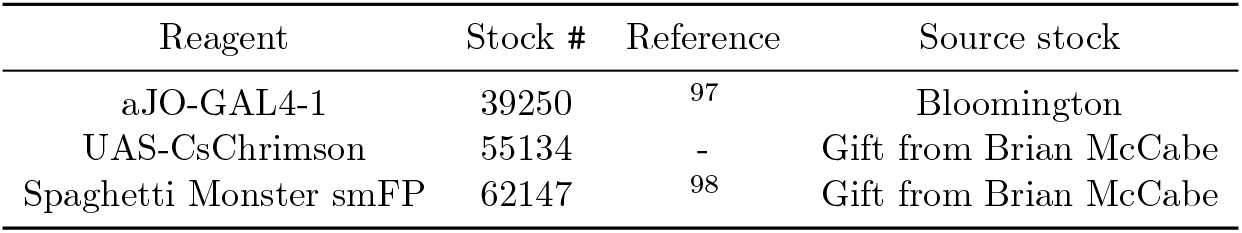
Fly strains used in this study.

#### Behavior recording system

Tethered fly behaviors were recorded using a previously described 7-camera (Basler, acA1920-150 *µ*m, Germany) system^46^ with the exception that rear right and rear left cameras were ignored, which does not capture anterior grooming behaviors. Animals were illuminated with an infrared (850 nm) ring light (CSS, LDR2-74IR2-850-LA, Japan). To track the positions of each leg joint, five cameras were equipped with 94 mm focal length 1.00×InfiniStix lenses (Infinity, 94 mm 1.00×, USA). Cameras recorded data at 100 fps and were synchronized with a hardware trigger. The full field-of-view (FOV) of each cameras is 1920x1200 pixels with a pixel size of 4.8x4.8 *µ*m. To reduce the size of the captured images and to increase acquisition rate, we set the ROI of each camera to 960x480 pixels. Flies were tethered to a wire, but otherwise freely behaving upon an air-supported (0.8 L/min) spherical treadmill. Foam balls were manually fabricated to be 10 mm in diameter (Last-A-Foam FR-7106, General Plastics, Burlington Way, WA USA, density: 96.11 kgm^−3^).

#### Confocal imaging

We dissected the brain and VNC from 3-6 dpe female flies as described in ref^61^. Primary and secondary antibodies were applied for 24 hrs and sample was rinsed 2–3 times after each step (for details, see ref^99^). Antibodies and concentrations used for staining are indicated in Table 2. Samples were imaged using a Carl Zeiss LSM 700 Laser Scanning Confocal Microscope. Standard deviation z-projections of imaging volumes were made using Fiji^100^. We rotated, translated, and modified the brightness and contrast of images to enhance their clarity (**Extended Data Fig. 1A**).

**Table 2.** List of antibodies used for immunofluorescence tissue staining in Extended Data Fig. 1A.

| Type | AB name | Dilution | Company | AB ID |
| --- | --- | --- | --- | --- |
| 1° | Anti-Bruchpilot (mouse) nc82 | 1:20 | Dev. Studies Hybridoma Bank | AB2314866 |
| 1° | GFP Tag Rabbit | 1:500 | ThermoFisher | G10362 |
| 2° | Goat anti-Mouse Alexa 633 | 1:500 | ThermoFisher | A21052 |
| 2° | Goat anti-Rabbit Alexa 488 | 1:500 | ThermoFisher | A11008 |

#### Tethering for behavioral measurements

For optogenetics experiments, we used a stick-tether method. First, we cooled a copper apparatus housing a fly-sized ‘coffin’ on a cold plate for 10 minutes. This is used to keep the fly anesthetized during tethering. The fly was then gently placed inside this coffin using forceps, and its position was adjusted using a brush. If the thorax was misaligned, the coffin was rotated (using a knob) to reposition the fly upright.

A silver wire, 0.2032 mm in diameter (A-M Systems, Silver 0.008” 25 feet wire), was then glued (UV-curable glue, Bondic, Aurora, ON Canada) to the fly’s scutoscutellar suture and cured with UV light. This wire was connected to a female contact (Distrilec, female contact size 20 7.5 A 14458991) that was then inserted into a corresponding male contact on the experimental setup, securing the fly onto an air-suspended ball or spherical treadmill. Each experiment began at least ∼ 30 minutes after tethering to allow the fly to acclimate to its environment. Experiments were performed at 25 ^°^C and 50% humidity, in the dark.

#### Optogenetic stimulation

For optogenetic stimulation, we used a 617 nm LED (ThorLabs, M617L3) mounted behind a lens (Thorlabs, LA1951–N–BK7 Plano-Convex Lens) to deliver ∼ 6.0 mW/mm^2^ intensity light to the fly from the right-anterior side (as shown in Fig. **1**). The entire anterior body of the fly was thus illuminated. For flies used in Fig. **1** and Fig. **3**, we delivered step pulses of 2-3 s duration, with at least 30 s intervals. In Fig. **5**, stimulation patterns included both step pulses and flickering of varying periods and frequencies. All experiments were conducted in the dark.

#### Air puff stimulation

To elicit antennal grooming, humidified, non-odorized air was delivered at the fly’s antennae, deflecting them towards the head. Mass flow controllers (MFCs, Bronkhorst High-Tech B.V., Netherlands) supplied regulated air flow at 70 mL/min. Airflow was diverted using six solenoid valves (SMC, S070C-6AG-32, Japan) controlled by an Arduino UNO (Arduino, Italy). This air was delivered to the fly’s antennae by way of a glass capillary held by a probe holder (MXB, Siskiyou Corporation, USA) linked to a post (ThorLabs, MS3R) and positioned facing the fly’s head. To better target air puffs, the glass capillary was pulled to thin its edge (P-1000, Sutter Instrument, USA; parameters: Pull: 0; Velocity: 10; Heat: 502; Pressure: 500).

To compare air puff- and optogenetically-induced antennal grooming, we stimulated the same individual flies in alternation using a custom Arduino script to switch between the two stimulus sources.

#### Morphological perturbation experiments

To investigate the role of sensory feedback in body part coordination, we first recorded optogenetically-elicited antennal grooming in intact flies over 5 trials (each with 2 stimulation periods and ∼ 40 s intervals). We then used cold anesthesia to surgically remove sensory feedback via leg amputation, antennal amputation, head fixation, or multiple combinations of two of these perturbations.

To amputate the legs, flies were first cold anesthetized. We then extended a foreleg using forceps and amputated it near the thorax-coxa joint with clipper scissors (FST, Clipper Neuro Scissors, no. 15300-00, Fine Science Tools GmbH). This, rather than pulling, ensured that the VNC would not be damaged. To prevent desiccation and movement of the remaining leg piece, we sealed the stump with UV-curable glue. To amputate the antennae, we used two precision forceps to gently remove the pedicel and funiculus by pulling the antenna away from the head. To fix the head at its resting position, we applied a small drop of UV curable glue between the dorsal head and anterior thorax, avoiding head bristle deflection. After each surgery, we ensured that the flies could still actively move their other body parts; those that could not were discarded. Experimental flies were then placed on a spherical treadmill for 5 trials after a 20-minute acclimation period. For two-body part perturbation experiments, we repeated the same procedure and conducted an additional 5 trials to ensure comparisons were made using the same flies across conditions.

### Data processing

#### 2D & 3D pose estimation

To quantify foreleg and head kinematics during antennal grooming, we used DeepLabCut^53^ and Anipose^47^. We annotated 10, 11, and 9 key points on the forelegs, head, and thorax, respectively. The first two sets of key points were used to calculate joint angles, while the thorax key points were used to help align the fly’s 3D pose within a common coordinate system. Three neural networks were trained for distinct sets of cameras: the front camera (camera 3), the front-right and front-left cameras (cameras 2 and 4), and the side-right and side-left cameras (cameras 1 and 5), as shown in Fig. **1**. We used DeepLabCut v2.2.1 to annotate camera images and train ResNet50 models. Each network was trained on ∼ 650 manually annotated frames for 500,000 epochs with batch size of 8, using a 95-5% train/test ratio. The dataset included primarily anterior grooming behaviors, including those with variations in head and foreleg configurations across different conditions such as leg or antennal amputation. Several iterations of training were conducted to correct outlier frames, culminating in a final comprehensive training phase where all annotated frames were merged and networks were retrained from scratch.

For 3D pose reconstruction, we used Anipose v0.9.0 and calibrated five cameras with a ChArUco board. The pattern was designed using OpenCV v4.5.5 with the board marker dictionary number 250 (`aruco.DICT_4×4_250`) and 4-bit markers^101^. The board contains 7x6 squares, with each marker measuring 0.225 mm and each square 0.300 mm. The pattern size is 2.1×1.8 mm, and the board size is 2.4×2.1 mm (± 0.1 mm). The board, printed on Opal with Blue chrome etching by Applied Image (Rochester, NY), features an etching dye that minimizes light reflection from the infrared (IR) ring-light, reducing interference with the cameras. We attached the board to a pin, allowing smooth movement while maintaining stability when inserted into the fly holder. The calibration video was captured at 40 FPS with full FOV (1920x1200 pixels) for 2 min, ensuring that the board remained visible and in focus in at least two cameras simultaneously. We used Anipose for marker detection and manually verified the accuracy frame-by-frame. The video acquisition and calibration process were repeated until the intrinsic and extrinsic camera matrices aligned with expected values. For instance, we verified that the camera locations from the calibration process matched those in our behavioral recording setup for the extrinsic values.

We performed 3D pose reconstruction on filtered 2D pose tracking data using the Viterbi filter provided in Anipose. We chose this filter for its simplicity and effectiveness ^47^. The filter window length was set to 25 frames (250 ms), which preserved rapid behavioral movements while mitigating most outliers. Since camera calibration is a one-time process, the quality of pose reconstruction can degrade due to environmental factors such as changes in lighting or slight shifts in cameras’ positions. To improve the robustness of our reconstructions, we enabled animal calibration and extended the number of iterations while tightening the tolerance in the bundle adjustment algorithm, which increased the processing time (these adjustments are available in the repository: https://github.com/gizemozd/anipose/tree/master). We disabled Ransac triangulation and activated spatiotemporal regularization.

The y and z axes of the 3D coordinate system were aligned with the vector from the right to left dorsal humeral, and from the left ventral to dorsal humeral bristles, respectively. The x axis was defined as the cross product of the y and z axes. The thorax “mid-point” was designated as the origin of the coordinate system, as these key points were minimally occluded in the recordings, providing sufficient stability. For more details, refer to the code, which includes all configuration files and a page explaining parameter choices (https://github.com/NeLy-EPFL/kinematics3d).

#### 3D pose alignment & inverse kinematics

To calculate joint angles, we first align the experimentally acquired 3D poses to a template biomechanical fly model (NeuroMechFly v2^49^), using a process also called *scaling* ^102,103^. This alignment is performed in two stages. First, we calculate the distances between key body landmarks to derive scaling factors that adjust the experimental 3D data to match the body segment proportions of the biomechanical model. These landmarks include the Thorax-Coxa and Claw (or Tibia-Tarsus joint) for each foreleg, base and tip of each antenna, and mid-wing hinge to mid-antennae for the head (when the fly is stationary). This process yields five scaling factors—two for the forelegs, two for the antennae, and one for the head. We then multiply each scaling factor with the corresponding limb, allowing us to match the task space of the real animal with that of the biomechanical model. In the second stage, we translate the positions of “fixed” joints (e.g., Thorax-Coxa, Base Antenna joints) to their respective locations on the biomechanical model. This two-step process aims to (i) reduce noise from jittery fixed key points and (ii) minimize leg size variations caused by triangulation or false positives in pose tracking.

Conventional optimization-based inverse kinematics methods aim to match the end effector position closely but often disregard the positions of preceding joints, leading to unrealistic movements of kinematic chains. To track each joint position closely, we developed a sequential inverse kinematics method, constrained by the fly’s exoskeleton^54^, also known as “body movement optimization”^104,105^. Our approach begins with the proximal-most leg segment to calculate the degrees of freedom (DOF) angles for the next joint. It then sequentially extends the kinematic chain by adding one segment at a time, repeating this process until it reaches the chain’s tip. This method is performed in four steps as follows:

- **Stage 1:** The kinematic chain includes only the coxa, used to calculate Thorax-Coxa yaw (rotation around the anteroposterior axis) and pitch (rotation around the mediolateral axis) by following the coxa tip as the end-effector.
- **Stage 2:** The chain extends to the coxa and the trochanter + femur (fused), calculating Thorax-Coxa roll (rotation around the dorsoventral axis) and Coxa-Trochanter pitch, using the femur tip as the end-effector.
- **Stage 3:** The chain includes the coxa, trochanter + femur, and tibia, used to calculate Trochanter-Femur roll and Femur-Tibia pitch by following the tibia tip as the end-effector.
- **Stage 4:** The full leg is included to calculate Tibia-Tarsus pitch, using the claw as the end-effector.

Our pipeline builds on the open-source inverse kinematics library IKPy^106^, which uses SciPy’s least squares optimizer^107^ to minimize the Euler distance between the original end-effector pose and the forward kinematics derived from the calculated joint angles.

Since the head has two moving parts (left and right antennae) parented by the main neck joint, the kinematic chain method can introduce errors by favoring one antenna over the other.

To avoid this, we calculated neck and antennal joint angles using the cosine angle formula between two vectors. The vectors for the head joint angles are defined as follows:

- **Head roll:** The angle between the vector from the right antenna base to the right antenna tip and the global mediolateral axis in the transverse plane.
- **Head pitch:** The angle between the vector from the neck to the mid-antennae base and the global anteroposterior axis in the sagittal plane.We subtracted the resting head pitch angle from the calculated joint angles to obtain joint angles relative to the resting position.
- **Head yaw:** The angle between the vector from the right antenna base to the right antenna tip and the global anteroposterior axis in the dorsal plane.
- **Antennal pitch:** The angle between the vector from the neck to the antenna base and the vector from the antenna base to the antenna tip in the sagittal plane.
- **Antennal yaw:** The angle between the vector from the right antenna base to the left antenna base and the vector from the antenna base to the antenna tip in the transverse plane.

Note that, when head rotation reaches 90°, the antennal pitch and yaw calculations switch roles, leading to inaccuracies. To avoid this, we first calculate the head joint angles, then derotate the head key points by the head rotation to compute the antennal joint angles.

Performance-wise, the entire pipeline takes 36 s to run inverse kinematics for six legs on 100 frames, using a MacBook Pro with a 2.3 GHz Quad-Core Intel Core i7, when parallelized. Our method is publicly accessible at^54^:

https://nely-epfl.github.io/sequential-inverse-kinematics.

#### Classification of behaviors

To investigate the kinematics of different antennal grooming subtypes, we labeled the recordings based on behavior. Seven distinct labels were used to annotate the videos. Five of these groups represent some variations of antennal grooming, while the remaining two correspond to other behaviors unrelated to antennal grooming. Each antennal grooming subtype is characterized by a specific coordination between the movements of the forelegs, head, and antennae. Using DeepEthogram v0.1.4 GUI^108^, we labeled each video frame for 33 trials across 10 flies (see **Supplementary Video 3**). The subtypes can be summarized as follows:

- **Bilateral grooming**
  - Both antennae are cleaned simultaneously.
  - The forelegs move synchronously, typically at the same height.
  - Frequent observation of head pitch, with occasional slight head roll (∼10°).
- **Unilateral antennal grooming (right or left)**
  - Grooming is focused on a single antenna.
  - The head rotates towards the groomed antenna, lowering it.
  - The non-groomed antenna actively lifts up.
  - The forelegs target the groomed antenna, shifting their position to one side.
  - The head is slightly pitched downward.
- **Unilateral non-tripartite antennal grooming (right or left)**
  - A single antenna is groomed, but not all conditions described in unilateral antennal grooming are met.
  - A single leg is raised to touch one antenna.
- **Non-classified grooming**
  - The forelegs are not in contact with the head but hover in front of the fly, typically at the level of the maxillary palps.
  - Involves other forms of anterior grooming, such as head grooming.
- **Background**
  - Behaviors outside of the anterior grooming such as foreleg rubbing, resting, or locomotion.

### Data analysis

#### Transitions between behaviors

We computed the transition frequencies between grooming subtypes. Each time point was assigned a behavior label, and we counted the number of transitions from one label to another during each trial, ignoring transitions with the same label.

For visualization purposes, we used the NetworkX^109^ library to create a directed graph, where each node represents a behavior, and the edges indicate the transition frequencies between behaviors. To represent these transitions as a probability matrix, we converted the graph into a matrix using NetworkX and normalized each row by the sum of its values, ensuring that the transition probabilities from one behavior to all others sums to one.

#### Dimensionality reduction using PCA

To reduce the dimensionality of optogenetically-induced antennal grooming kinematics data, we performed Principal Component Analysis (PCA). We first identified kinematic variables showing the most significant changes during antennal grooming. These included joint space variables such as antennal pitch, head pitch and roll, thorax-coxa pitch and roll, coxa-trochanter pitch and roll, as well as the 3D positions of the antennal base and tip, and the foreleg Tibia-Tarsus joints in the transverse plane. In total, we had 28 time series inputs for dimensionality reduction (12 joint angles and 16 joint positions in 3D). Each kinematic variable was standardized to have zero mean and unit variance. We then partitioned the dataset (size *N*_timesteps_, 28) into 10-time-step chunks (*N*_chunks_, 10, 28) using a sliding window of 8. As the sampling rate of the data is 100 fps, this amounts to the data partitions of 100 ms with a 20 ms of overlap. To ensure that each chunk contained continuous time series data, rather than transitions between trials, this process was performed on a trial-by-trial basis.

Note that each chunk is assumed to represent one behavior; however, a chunk might be populated by several behavioral labels. To ensure data chunks predominantly reflected a single behavior, we excluded chunks with fewer than 60% of the labels corresponded to a specific behavior. That is, we removed chunks with fewer than six labels for a given behavior. Additionally, chunks labeled as ‘background’ were excluded, as this category includes a diverse set of behaviors unrelated to antennal grooming. After this filtering, we retained 2,537 chunks as data points. Next, we reshaped our kinematic matrix (*N*_chunks_, 10, 28) into a 2D array (*N*_chunks_, 280) for PCA. Five principal components captured more than 60% of the variance in our data. For visualization, we plotted the first two columns of the weight matrix as it captured 40% of the variance (**Extended Data Fig. 1F**). Each point, representing a chunk, was colored according to the most frequent behavioral label within that chunk.

#### Analysis of perturbation experiments

For each perturbation type, we first divided the kinematics during optogenetic stimulation into chunks of 300 ms with 50 ms overlaps for each fly after denoising single kinematic traces with a Savitzky–Golay filter (window size: 9, degree: 3). For each chunk, we checked if a given epoch of kinematics is free from outliers **(Table 3, outlier check)** and if any of the body parts was moving **(Table 3, behavior check)**. If these conditions were not met, we discarded the chunk. Valid chunks were labeled based on the movements of freely moving body parts **(Table 3, right label condition)**. For head-fixed flies, we labeled chunks based on antennal movements, using the difference between left and right antennal pitch angles. If this difference exceeded a set threshold, we annotated the chunk according to the lifted antenna. The annotated data was then used to plot the distribution of Tibia-Tarsus joint positions in the mediolateral plane (**Fig. 3E**). Similarly, we used antennal pitch angles to annotate leg amputation experiments, but we plotted the head rotation angles this time (**Fig. 3G**). For antennae amputation, we designated the labels based on the head rotation: chunks were annotated if the median head roll angle fell within a certain range; otherwise, they were labeled as either right or left based on the direction of rotation. As with head-fixed experiments, we then plotted the Tibia-Tarsus joint position distribution (**Fig. 3I**). The thresholds used for each kinematic variable are listed in (**Table 3**).

**Table 3.**
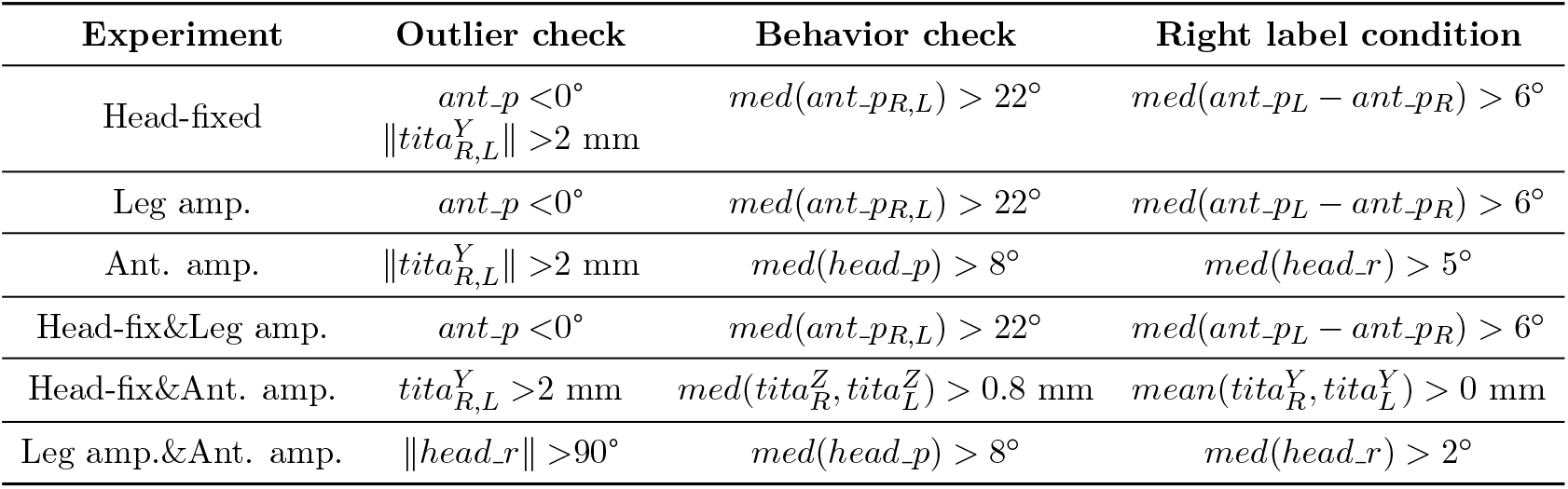
Thresholds used for annotating behavioral chunks in Fig. 3. Variables shown: *ant p*: antennal pitch, *head r*: head roll, *head p*: head pitch, *tita*: tibia-tarsus joint position. Only conditions for labeling a chunk ‘right’ are shown: labeling the ‘left’ is simply the opposite.

For phenotype analysis (**Fig. 3E-I, bottom left**), we calculated the median value for each biological replicate (i.e., each fly) for both phenotypes (left and right) and visualized them using scatter plots. To compare intact and experimental flies, we examined the difference between these left and right label values, using the median across all trials for each fly to measure the variation in joint configurations (**Fig. 3E-I, bottom right**). This approach allowed us to assess the range of movement for a given degree-of-freedom between the left and right behavioral conditions.

From two-body part perturbations, we compared the movement range of the remaining body part to that in intact flies. For head-fixed and foreleg-amputated flies, we first checked for outliers and antennal pitch movement. If a chunk was valid and one antenna was pitching, we proceeded with the chunk corresponding to the upward-pitched antenna. We repeated the same procedure for the head roll in antennae- and leg-amputated flies, and for the lateral position of forelegs in antennae-amputee and head-fixed flies. Each valid chunk was labeled based on the direction of the freely moving body part (i.e., right or left). The distribution was then plotted using all valid chunk data (**Fig. 3K-O, left**).

To compare flies across different conditions, we calculated the difference between the 90^th^ and 10^th^ percentile values of antennal pitch as a proxy for the movement’s maximum and minimum range (**Fig. 3K right**). For the head roll and Tibia-Tarsus joint positions, we took the median of all left-labeled chunks from each fly and subtracted it from the median of the right-labeled chunks (**Fig. 3M,O right**). We kept the fly identities across conditions, indicated by a line between each dot in the scatter plots (**Fig. 3E,G,I,K,M,O**).

### Kinematic replay and antennal contact detection

To infer limb-antennal contacts, we performed kinematic replay using the updated fly biomechanical model from^49^ in the physics engine MuJoCo v2.3.7^110^ and integrated in the FARMS framework^50^.

Simulating antennal grooming in a physics engine poses several challenges. First, because there are numerous contact points between the antennae and foreleg meshes, extensive collision detections are required at every time step. Second, we used mesh-based collision bodies, including complex geometries, further increasing the computational load.

To address these issues and to ensure smooth kinematic replay, we implemented several optimizations. First, we reduced the time step to 10^−4^ s and limited the simulations to short snippets (around 5 s) to increase the stability of integrators and to avoid error accumulation throughout the simulation. We also fine-tuned the physics engine parameters, using the ‘Projected Gauss–Seidel’ solver with an Euler integrator. We increased the number of solver iterations to 10^7^ and lowered the residual threshold to 10^−10^ to improve stability. Additionally, to speed up the simulation, we restricted collision detection to only between the forelegs and head segments.

In total, we actively controlled 16 degrees of freedom: 7 for each foreleg, as described in^48^, and 2 for head roll and pitch. We set the antennal joints as passive (following spring-damper dynamics). We maintained a fixed resting pose because replaying measured antennal joint angles introduced confounding factors due to collisions occurring during these measurements. To better emulate the kinematics of unilateral grooming, we adjusted the antennal joint angles (e.g., pedicel and funiculus pitch) to different values, placing the non-groomed antenna in an upward pitch position. We empirically tuned the joint damping and stiffness parameters to qualitatively mimic passive antennal movements after contact. All passive antennal joint angles are provided in Table 4.

**Table 4.** Resting positions of passive joint angles given in degrees.

|  | Head yaw | Pedicel roll, pitch, yaw |  | Funiculus roll, pitch, yaw |  | Arista roll, pitch, yaw |  |
| --- | --- | --- | --- | --- | --- | --- | --- |
| Behavior | - | Right | Left | Right | Left | Right | Left |
| UniL | 0 | 0, -60, -35 | 0, -40, 35 | 10, -25, 0 | -10, -10, 0 | 0, 0, 35 | 0, 0, -35 |
| UniR | 0 | 0, -40, 35 | 0, -60, -35 | 10, -10, 0 | -10, -25, 0 | 0, 0, 35 | 0, 0, -35 |
| BiLat | 0 | 0, -40, 35 | 0, -40, 35 | -10, -10, 0 | -10, -10, 0 | 0, 0, 35 | 0, 0, -35 |

To visualize collisions between the antennae and leg segments (**Fig. 1H** and **Fig. 2B,E,H**), we binarized the contact force read-outs, converting any non-zero forces between a collision pair to 1, representing “contact on” time points. For articulated parts—pedicel, funiculus, arista for antenna, and tarsi 1-5 for the tarsus—we combined the binarized contact arrays into a composite contact array by taking their union. The resulting binary contact arrays between each antenna and leg segment pair over time were displayed as collision diagrams.

To quantify contact duration (**Fig. 2C,F,I**), we summed each contact array over time for a given collision pair, representing the total contact duration for that trial. We performed kinematic replay at different gains and normalized the data to a 0-1 range based on the maximum and minimum values observed. This normalization allowed us to identify the gain at which maximum contact occurred for each trial. We modulated degree of freedom kinematics by multiplying the original joint angles with a constant factor (attenuating ∈ [0, 1) or amplifying ∈ (1, 1.3]) while keeping the kinematics of other degrees of freedom unchanged. Due to noise and outliers in the 2D and 3D pose estimation, and variations in animal morphology, the mapped kinematics might not always yield an accurate kinematic replay of the behavior. To mitigate this effect, we ignored simulations where the detected contacts lasted less than 10 ms, amounting to, on average, ∼ 2% of a simulation trial.

To quantify contact forces (**Extended Data Fig. 4C**), we calculated the median of all nonzero contact force values for each replay experiment. Specifically, we measured contact forces between the tibia and its ipsilateral antenna, and between the tarsus and its ipsilateral antenna. Each data point in the distribution represents the median contact force for a single trial. This process was repeated across multiple animals/trials and gain values.

### Connectome analysis

#### Loading connectomics data

As for brain connectivity analysis, we used the female adult fly brain (FAFB^20,19^) connectomics dataset from Codex (FlyWire materialization snapshot 783; https://codex.flywire.ai/api/download) to generate figures Fig. **4**, Extended Data Fig. 7B-F and Extended Data Fig. 8. We also used the male adult nerve cord (MANC, version 1.2.1^111,25^) dataset using NeuPrint Python API to generate figures Extended Data Fig. 7G-H.

#### Constructing the antennal grooming network

We constructed a comprehensive antennal grooming network in two stages, starting with a smaller foundational network and then expanding it by exploring its neighboring connections.

To build the foundational network, we first identified key antennal grooming-related neurons in the brain connectome, including JO-C/E/F, antennal bristles, aBN1-3, and aDN1-3^37,40^. JO-C/E/F and antennal bristle mechanosensory neurons were selected because they are known to trigger antennal grooming^39,40^. We also included antennal and neck motor neurons in this foundational network, as they act as the output layer of this system.

We used FlyWire Community labels^19,20,55^ to identify neck motor neurons, but similar labels were not available for antennal motor neurons. To address this, we examined motor neurons passing through the antennal nerve, focusing on their branching patterns. Among the seven pairs of motor neurons we found, two primarily received inputs from visual neurons and were likely retinal motor neurons^112^. Therefore, to narrow down the search space of network, we excluded those two motor neurons. Among the remaining five, several motor neurons received inputs from JO and aBN neurons, suggesting a role in antennal motor control.

Having established the foundational network, we next expanded it by mapping out all monosynaptically connected neurons using the connectivity diagram from the FAFB connectome (**Extended Data Fig. 7A**). In particular, for each of these monosynaptically connected neurons, we calculated the percentage of synapses incoming from and outgoing to foundational network neurons. Neurons with connectivity percentages below a predefined threshold were pruned from the network. Depending on the neuron type (e.g., descending neurons sometimes lacking axon terminals in the brain and sensory neurons lacking dendrites), we applied specific rules based on “super class” annotations in FAFB to guide the pruning process, described below:

- **Sensory neurons:** outgoing synapse percentage *>* threshold
- **Interneurons:** incoming syn. perc. *>* threshold *and* outgoing syn. perc. *>* threshold
- **Descending neurons:** incoming synapse percentage *>* threshold

We excluded ascending neurons and other sensory neurons from our network, as their role in antennal grooming is not yet well characterized. To ensure that we capture only neurons with significant information exchange, we applied a threshold of 5%, accounting for only those neurons that contribute to at least 5% of the input/output interactions within our predefined network. This threshold is ten times lower than the threshold after which the algorithm does not find any new neurons (**Extended Data Fig. 7B**), suggesting that 5% is sufficiently high to discover new neurons. Furthermore, this choice of threshold is consistent with recent findings indicating that connections providing more than 1.1% of target neuron’s inputs are 90% more likely to be preserved across brains^20^. This process introduced approximately 240 new neurons to the network (**Extended Data Fig. 7C**). Because all leg motor neurons are located downstream of the brain within the VNC, descending neurons with more than 10 synapses upon foreleg-controlling (T1) leg motor neurons were defined as ‘leg premotor neurons’. These were limited to descending neurons that have been matched across brain^19–21^ and VNC^111,25^ connectome datasets^113^. We did not include foreleg motor neurons as a separate group, because they are part of the VNC dataset. Neurons that did not fit into any predefined categories were left unassigned.

Most of these neurons had contralateral counterparts, but due to differences in synapses between the left and right hemispheres, our network construction algorithm was not always able to find these pairs automatically. Therefore, to identify missing contralateral pairs, we used two approaches. First, we calculated dissimilarity scores between a neuron and all of its contralateral candidates. The dissimilarity score **D**_*ij*_ between neuron *i* and neuron *j* is given by

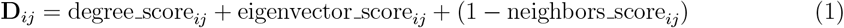

where

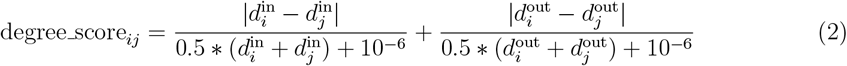

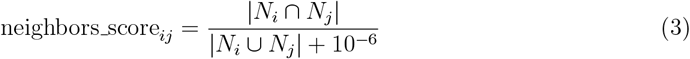

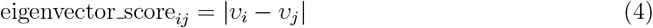

where *d*^in^, *d*^out^ in- and out-degrees, *υ* eigenvector scores, and *N* sets of neighbors of nodes. We obtained degree and eigenvector scores using the built-in NetworkX^109^ functions `degree_centrality` and `eigenvector_centrality`. Note that, for identical neurons, the dissimilarity metric becomes zero. We verified that dissimilarity scores for the same cell types were lower than those for different cell types. We used this approach to match sensory neurons. Specifically, we first divided sensory neurons into high-level classes, that is, JO-C/E/F and antennal bristles based on the Fly-Wire neuron classification (cell type attribute). Within each class, we then computed dissimilarity scores for each neuron pair, resulting in a global list of,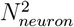 entries storing dissimilarity scores. We then assigned neurons by sequentially pairing those with the highest dissimilarity scores.

The remaining neurons were matched through a combination of community label matching, anatomical and biological comparisons (e.g., hemilineage), and the similarity of their neighbors. In cases where FlyWire had already identified a “mirror twin” neuron, we adopted its match. For neuron populations where individuals were indistinguishable, we assigned pairs randomly.

#### Connectivity analysis

For our antennal grooming network, we identified pre- and post-synaptic neurons, along with all the connections between them, including neurotransmitter types. We used the publicly available FlyWire brain connectome dataset to conduct these analyses. For each synaptic connection, we assigned the neurotransmitter with the highest prediction probability. To maintain consistency, we assigned the most commonly predicted neurotransmitter to each neuron and applied it uniformly to all of its connections. This rule, however, did not apply to neurons lacking axon terminals in the brain. To analyze connectivity in the VNC, we used the `neuprint.fetch_adjacencies` function to retrieve synaptic adjacency data.

#### Assigning neurons to groups

To investigate the coordination between antennal, foreleg, and neck motor neurons, we analyzed common connectivity motifs within our network. To simplify this task, we categorized the network into eight main groups: sensory neurons (JO-F), central neurons, antennal premotor neurons, neck premotor neurons, foreleg premotor neurons, shared premotor neurons, antennal motor neurons, and neck motor neurons. Since leg motor neurons are part of the VNC dataset and there is no one-to-one mapping between descending or ascending neurons in the brain and those in the VNC, we excluded them from our analysis. Neurons that did not fit into these categories were classified as “other.”

Motor neurons and JO-F sensory neurons were already annotated in the brain connectome. We defined premotor neurons as those having more than 5% of their total output directed toward motor neurons that control the same appendage. For example, an antennal premotor neuron projected more than 5% of its output to antennal motor neurons, but may have had less than 5% output to other types of motor neurons. Shared premotor neurons were those with more than 5% of their output projecting onto more than one type of motor neuron.

Because leg motor neurons are not in the brain, we also examined descending neurons (DNs) projections in the VNC to identify those with direct connections with T1 leg motor neurons (annotated as *MNfl* in MANC^80,25^). Since a completely proof-read “full CNS” connectome is not yet available, we relied on a recent study^113^ that bridged brain descending neurons with their VNC counterparts through light-level descriptions of their full morphology. Among the 188 DNs in our network, 75 were matched with their VNC extensions. However, a bijection could not be achieved for some DN populations because it was challenging to distinguish individual neurons whose axons travel together in bundles. Therefore, if any neuron in a population projected onto a leg motor neuron, we classified the entire population as ‘leg premotor’. We set a threshold of 10 synapses in the VNC to qualify as a premotor neuron. We employed the same approach for VNC neck motor neurons (annotated as *MNnm* in MANC^80,25^). Since the DNs presynaptic to VNC neck motor neurons belonged to the same DN population presynaptic to leg premotor neurons, we excluded neck premotor neurons in the VNC to avoid overestimating the number of shared premotor neurons.

After defining the premotor neuron classes, we proceeded to identify central neurons. Central neurons were defined as those located between the input (JO-F neurons) and output layers (premotor and motor neurons) of the network. To identify these neurons, we used the `networkx.all_simple_paths`^109^ function to generate all simple paths between source (JO-F) and target (premotor and motor neurons) neurons, with a maximum of four hops. We set the limit to four layers, as this has been shown to be sufficient to reach a majority of neurons in the fly nervous system^56,80,114^. To further refine the network and eliminate neurons with minimal synaptic contribution, we computed the average synaptic percentage, defined as the number of synapses between a pre- and post-synaptic neuron divided by the total synapses the presynaptic neuron makes. We discarded paths with an average synaptic percentage below 5%.

To create randomized networks (**Extended Data Fig. 9A,B**), we reassigned existing connections, along with their synaptic counts, to random pairs of presynaptic neurons (excluding motor neurons) and postsynaptic neurons (all neurons). The identities of JO-F sensory neurons, motor neurons, and leg premotor neurons were preserved. Using these predefined modules, we constructed other neuron groups (e.g., premotor and central) following the same procedure.

#### Graph visualizations

We used the NetworkX package^109^ to visualize network connectivity in Fig. **4**B,C,J; Fig. **6**H,I; Extended Data Fig. 8 and Extended Data Fig. 11C,F,G. From the connectivity table, we generated a directed graph where the source node represented the pre-synaptic neuron and the target node represented the post-synaptic neuron. The edge widths were proportional to synaptic counts. We used dark blue, light blue, and dark red to denote GABA, glutamate, and acetylcholine neurotransmitters, respectively, and gray for the remaining neurotransmitters (e.g., dopamine and serotonin). For graph visualizations in Fig. **4**J, Fig. **6**H,I and Extended Data Fig. 11F,G we exported NetworkX graphs in .gephx format and imported them into Gephi v0.10.1 to modify node locations and graph aesthetics.

### Connectome-constrained neural network modeling

#### Preparing the training dataset

The training dataset consisted of head kinematics (i.e., antennal pitch and head pitch joint angles) from leg-amputated flies. We optogenetically elicited antennal grooming (as described in Extended Data Fig. 1) following a recovery and habituation period of approximately 20 min. Data was collected from 10 flies; around 22 trials were conducted per fly. Each trial involved two types of stimuli: step inputs of varying duration (0.5, 1, 2, 3 s) and pulsatile inputs of varying frequencies (5, 10, 20 Hz) delivered over a 2 s period.

Measured antennal and head pitch angles were then used as an output dataset, and optogenetic stimulation patterns served as an input dataset. Stimulation values were coded 0 for off periods and 1 for on periods. Joint angles were scaled with 99^th^ percentile corresponding to 1 and 1^st^ percentile corresponding to 0. This effectively mapped the joint angle range to a 0-1 scale. Baseline subtraction was performed to ensure that the resting pose corresponds to 0. Since the model was simulated at a 1 ms resolution, we interpolated the data captured at 100 FPS to match this sampling rate.

To create fictive sensory feedback, we only used antennal pitch angles. Neck proprioceptive neurons are not yet fully characterized. By contrast, the antennal JO is well-studied and contains distinct populations of mechanosensory neurons (i.e., JO-C and JO-E), which are tuned to upward and downward movements of the antenna, respectively ^37,115^. From antennal joint angle traces, we first standardized the antennal joint angles such that the resting position of the antenna would correspond to zero. We then identified movements above and below the baseline, corresponding to upward and downward antennal movements. Since these signals would be provided as inputs to their corresponding sensory neurons, we converted negative values to positive. Additionally, we introduced a 40 ms time delay to emulate the sensorimotor delay between the creation of motor commands and their reception by mechanosensory neurons.

Each input-output pair represented a single stimulation period and had a fixed length of 3.8 s, allowing us to run the network with no input for a certain duration (with the longest stimulus being 3 s). To ensure that the loss was calculated only during optogenetic stimulation, we constructed a “mask” to indicate the start and end of stimulation, and only calculating the loss (and hence the gradient) during the stimulus periods. In total, we obtained 412 trials, resulting in an input dataset of (*N*_trials_ = 412, *N*_time_ = 3800, *N*_neurons_ = 852) and an output dataset of (*N*_trials_ = 412, *N*_time_ = 3800, *N*_joints_ = 3). A sample is shown in Extended Data Fig. 10A.

#### Adjacency matrix preparation

While constructing our antennal grooming network, we ensured that each neuron had a contralateral pair. However, we observed differences in synaptic connectivity between the left and right hemispheres, likely due to biological variability and imperfections in the connectome dataset. To eliminate the impact of this asymmetry on our results, we tried three different ways to make connections across both hemispheres symmetrical:

- **Maximum:** Set synaptic counts to the maximum observed for a neuron pair.
- **Minimum:** Set synaptic counts to the minimum observed for a neuron pair.
- **Average:** Set the synaptic counts to the average of observed for a neuron pair.

This resulted in three different adjacency matrices: 7,772 connections for the maximum and average methods, 2,148 connections for the minimum method, and 4,961 connections in the original network. The minimum method is likely to eliminate important connections and the average method might bias synaptic counts if a connection was missing on one side. Therefore, we used the maximum model.

Next, in our maximum adjacency matrix, we grouped neurons—except for sensory and motor neurons—based on the similarity between their pre- and post-synaptic connections. We calculated the Pearson correlation coefficient between two neurons’ upstream and downstream connections and summed them. Therefore, in this similarity matrix, two identical neurons will have an entry value of 2 whereas two highly different neurons will have a score of −2. From this matrix, we calculated a distance matrix, calculated my 0.5 * (2 − *A*_*sim*_) where *A*_*sim*_ is the similarity matrix.

Then, we applied the unsupervised clustering algorithm DBSCAN ^116^ to the distance matrix to cluster neurons. We performed a parameter search to optimize the algorithm’s parameters— epsilon (set to 0.5) and the minimum number of samples (set to 1)—and to ensure that the resulting clusters are biologically meaningful (i.e., each cluster is either excitatory or inhibitory). Any neuron that DBSCAN left unclustered was assigned as its own individual cluster. We set sensory neurons (i.e., JO-C/E/F and antennal bristle neurons) to their respective cell types. Furthermore, each right-left motor neuron pair was assigned to a different cell type. In total, we obtained 104 clusters, reducing the number of node type 8-fold (from 852).

We additionally trained models based on the three different adjacency matrices (maximum, minimum, and average), a shuffled, and the original (i.e., unprocessed) version for multiple seeds (**Extended Data Fig. 10C**). For the shuffled matrix, we started with the symmetric adjacency matrix from the maximum count approach, then randomly rearranged the post-synaptic connections of each neuron on one hemisphere while preserving neurotransmitter identity. This shuffled matrix was then mirrored across hemispheres to maintain symmetry. For all types of adjacency matrices, we used the same cell types obtained from the previously mentioned clustering process.

The training results revealed that the network with the minimum number of connections, and thus the fewest open parameters, performed the worst (**Extended Data Fig. 10C**). By contrast, the shuffled and maximum connection networks, which had the highest number of open parameters, achieved the smallest test errors. These findings highlight the trade-off between model complexity and computational efficiency. Notably, the original connectome network’s performance was between the maximum and minimum connection networks (**Extended Data Fig. 10C**).

#### Model parameters and training

We adapted the open-source Python package for connectome-constrained model training^51^ (https://github.com/TuragaLab/flyvis, commit 056e4aa) to the grooming network and dataset. Specifically, each neuron 0 ≤ *j* ≤ *N* is modeled as a leaky integrator neuron, where *N* is the number of neurons in the network, whose voltage dynamics *v*_*j*_(*t*) are governed by the following equations (trainable parameters are shown in red),

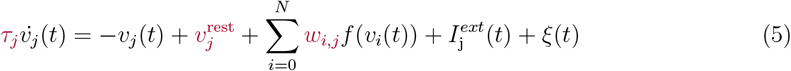

where *τ*_*j*_ is membrane time constant and,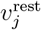 is resting potential, shared among neurons of the same cell-type (previously defined by an unsupervised clustering algorithm). Mechanosensory neurons JO-C/E/F receive an external input,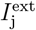 (i.e., sensory feedback and optogenetic stimuli) as described above while for other neurons in the network external input is zero. Each neuron also has an intrinsic noise *ξ* ∼𝒩 (0, 0.01) and receives input from its pre-synaptic neurons *i*. We used rectified-linear unit (ReLU) to model neurotransmitter release *f* (*x*) = *max*(0, *x*). Transformed membrane potential of each pre-synaptic neuron *i* to post-synaptic neuron *j* is then weighted by the synaptic connection between the two *w*_*i*,*j*_ as in reference^51^,

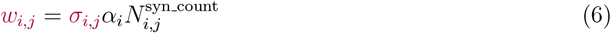

where,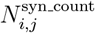 is the natural logarithm of the average synaptic count between cell types *i* to *j*, and *α*_*i*_ is the neurotransmitter sign that is − 1 if the detected neurotransmitter type is inhibitory and +1 otherwise. We assigned both GABA and Glutamate to be inhibitory^117^. We assigned Acetylcholine and Dopamine to be excitatory. Synaptic strength *σ*_*i*,*j*_ is an non-negative parameter and modulates the connection strength between neurons, initialized as

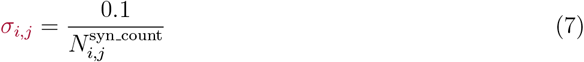

Neuron parameters were initialized within physiologically plausible ranges. Resting potential was drawn from a normal distribution,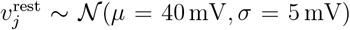, clamped to a minimum of 0 mV to prevent excessive positive bias in neurons. The membrane time constant was uniformly initialized to 30 ms for all neurons and constrained to remain within the range [0, 150 ms].

To transform motor neuron activity into joint kinematics, we designed two Multi Layer Perceptrons (MLPs) (**Table 5**) by using built-in PyTorch functions. Decoders are used to emulate the nonlinearities arising from the musculoskeletal properties of antennal and neck pitch joints. We choose a feedforward network, rather than a recurrent neural network, to limit the capacity of decoder. In particular, there are 5 pairs of antennal motor neurons and 4 pairs of neck pitch motor neurons in the brain. left and right antennal MN activities were passed separately through the same antenna decoder, assuming that the left and right antennal muscles have identical properties. For the neck pitch motor neurons, both left and right activities were passed through a single neck decoder. In total, we had 3 output traces, obtained as follows:

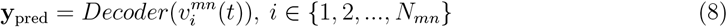

where,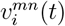 denotes motor neuron voltage values, and *N*_*mn*_ is the number of motor neurons. The model parameters are optimized through Backpropagation Through Time (BPTT)^67^ to minimize the loss between the decoder output and measured kinematics for every output trace, as described below:

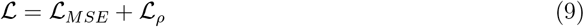

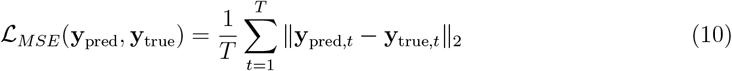

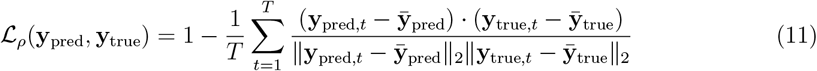

**Table 5.**
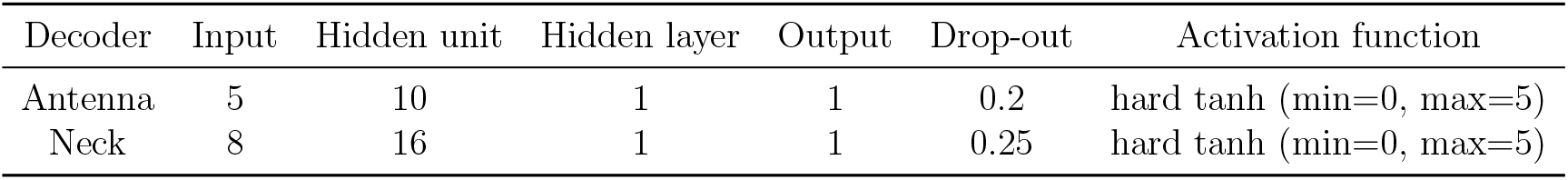
Antennal and neck motor decoder parameters.

where *T* is the number of time points,,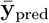 and,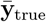 are the mean of **y**_pred_ and **y**_true_, respectively. ℒ_*MSE*_ and ℒ_*ρ*_ denote mean-squared-error and correlation losses.

Our connectome-constrained brain model had *N*_*cell type*_ ×*N*_*neuron param*_ = 104 × 2 = 208 open parameters for neuronal dynamics and 624 (number of unique connections between cell types) open parameters for weights, giving a total of 832 parameters. We used the optimizer AMSGrad^118^, a variance of Adam optimizer with a learning rate of 10^−4^ and batch size of 16 over 5,000 iterations. The models were trained using NVIDIA Hardware (GeForce RTX 2080, GeForce RTX4080, and V100). Each model took about 3-6 days to complete training.

#### Computational neural activation and silencing experiments

In our activation screen, we simultaneously delivered bilaterally symmetric input (left: 1.5, right: 1.5) to the JO-F neurons and unilateral input (left: 10, right: 0) to each neuron in the network for 2 s. The goal of this experiment was to identify neurons whose activation was sufficient to convert bilateral grooming into unilateral grooming.

In our silencing experiment, we bilaterally silenced neurons by setting all of their post-synaptic connections to zero. An bilaterally asymmetric step input (left: 3, right: 5) was then given to JO-F neurons. Here the objective was to identify neurons whose silencing disrupts unilateral aMN4 responses.

In both experiments, the aMN4 response was quantified using our response metric **(Fig. 5E)** after denoising single neural traces with a Savitzky–Golay filter (window size: 11, degree: 3).

Additionally, global network activity was evaluated after silencing neurons by comparing intact network activity to post-silencing activity. Neurons with mean activity more than five times the intact network activity during stimulation were labeled as ‘highly active.’ The total number of highly active neurons was then counted. Note that, this metric did not account for neurons that decreased global network activity after silencing.

All analyses were performed at the cluster (population) level by averaging the activity of all neurons within each cluster. Neural activity was passed through a ReLU activation function before analysis.

#### Identifying network motifs

Neurons were identified as important in our activation screen if their median USI metric (across three models) was either greater than 0.1 or less than -0.1. These neurons were categorized based on their effect of driving either ipsilateral aMN4 or contralateral aMN4.

To compute cluster weights from the single-neuron adjacency matrix, we summed the presynaptic and postsynaptic connections of all neurons within each cluster. Since each cluster is exclusively inhibitory or excitatory, the sign of the summed weights was preserved during this operation.

When visualizing network motifs, we included neurons that had both incoming and outgoing synapses within the motif, except for motor neurons. To simplify the network diagram, connection weights below the 1^st^ percentile of all weights in the network were omitted from the visualization.

### Statistical analysis

All statistical analyses were conducted using Python v3.10 with SciPy v1.10.1^107^. Unless otherwise specified, we employed the Mann-Whitney U test, a non-parametric method that does not assume any particular underlying probability distribution of the samples.

In Fig. **2**C,F,I and Extended Data Fig. 4C, we compared force or contact read-outs across different gain values (modified versus intact) and between two leg segments (tibia versus tarsus) using a two-sided Mann-Whitney U test. Specifically, each trial is summarized with a single value using a metric, and distributions of these trial values were used for comparison. Each gain value was compared to the natural behavior (gain = 1 or gain = 60°), using the natural distribution repeatedly. To account for multiple comparisons, we applied the Simes–Hochberg false discovery rate correction, with a significance threshold of *α* = 0.05^119^.

In Fig. **3**E,G,I,K,M,O, we performed within-fly comparisons between two phenotypes (uniR versus uniL kinematics; **Fig. 3E,G,I** bottom left) and between experimental conditions (intact versus experimental animals; **Fig. 3E,G,I,K,M,O** bottom right). We summarized data from each fly by taking the median of all trials. For phenotype comparisons, we used a one-sided Mann-Whitney U test, with the alternative hypothesis selected based on the specific kinematic variable. Comparisons between intact and experimental animals were made using a two-sided Mann-Whitney U test.

In all figures showing statistical tests, significance levels are indicated as follows: ***: *P <* 0.001, **: *P <* 0.01, *: *P <* 0.05 and not significant (NS): *P* ≥ 0.05. Sample sizes and P values are described in the respective figure legends.

## Data availability

Data are available at:

https://dataverse.harvard.edu/dataverse/ozdil_2024_antennal_grooming

This repository includes behavioral recordings used in Fig. **1**, Fig. **3**, Extended Data Fig. 1, Extended Data Fig. 2, Extended Data Fig. 3, Extended Data Fig. 6; trained DeepLabCut networks to perform 2D pose estimation; a table representing the antennal grooming network used in Fig. **4**. Raw behavioral videos are available upon request from the authors and are omitted here due to storage limitations.

## Code availability

Code is available at:https://github.com/NeLy-EPFL/antennal-grooming

## Extended data

**Extended Data Fig. 1.**
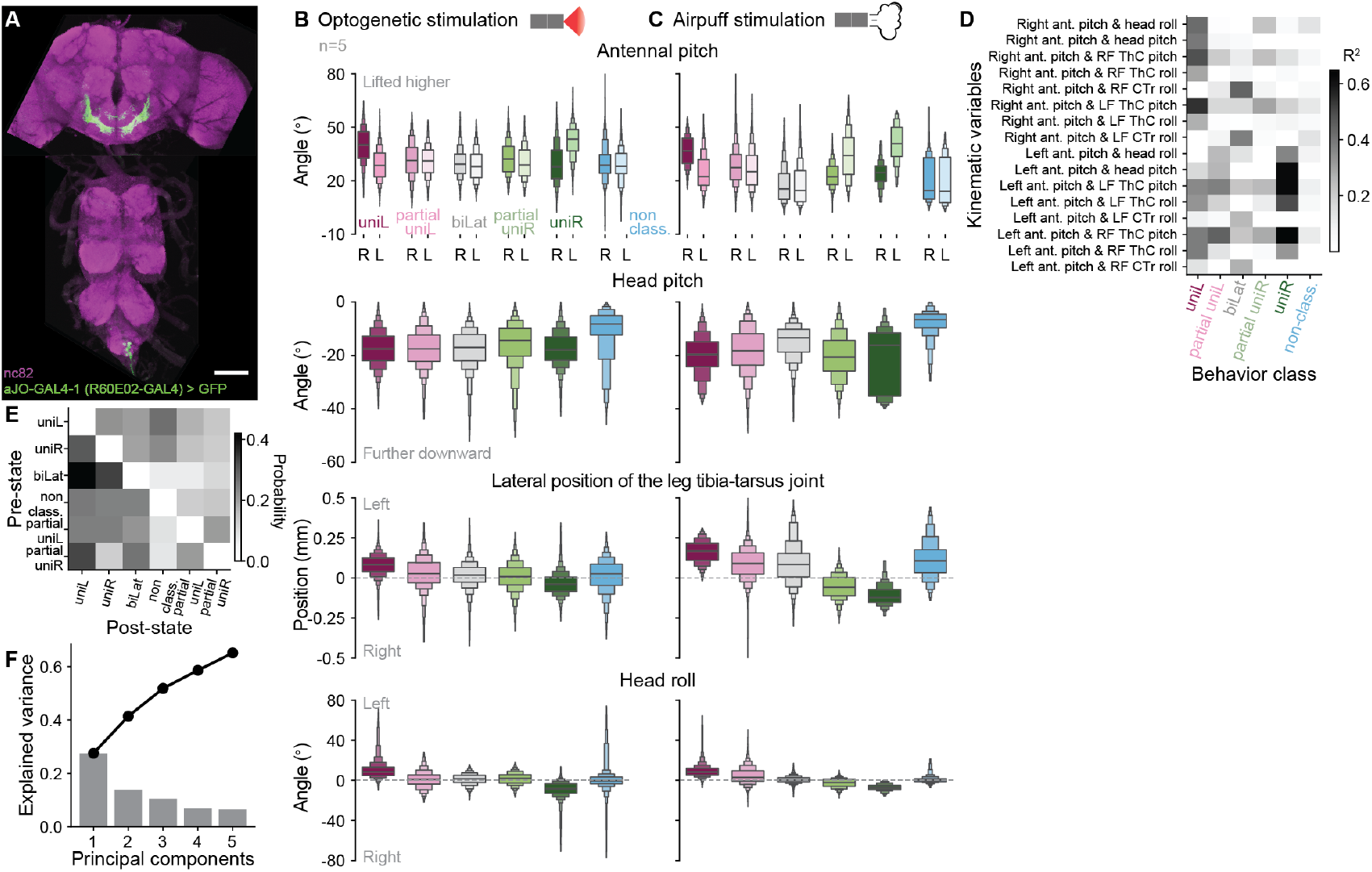
Characterization of antennal grooming. **(A)** Confocal image showing nervous system expression for the *aJO-GAL4-1* ^40^ driver line used to optogenetically-elicit antennal grooming. GFP (green) and nc82 (purple) are stained. Scale bar is 100 µm. **(B-C)** Boxen plots showing the distribution of kinematic variables during **(B)** optogenetic-or **(C)** air puff-elicited antennal grooming. These include antennal pitch **(first row)**, head pitch **(second row)**, tibia-tarsus joint position **(third row)**, and head roll **(bottom row)**. Data are color-coded by grooming class. In **(B)**, light and dark shades represent the left and right antennae, respectively. For all boxen plots, the center line represents the median, and each successive box denotes a halved quantile range of the data. Data are taken from n=5 flies. **(D)** Squared Pearson’s correlation (*ρ*^2^) between joint angles (rows) as a function of antennal grooming class (columns). Darker boxes indicate higher correlation. **(E)** Transition matrix between antennal grooming subtypes. Self-transitions are excluded and row values are normalized to sum to one. **(F)** Explained variance for the first five principal components. Bar graph shows the individual contribution of each principal component to the total variance. Line plot shows the cumulative explained variance. Data are combined from panels **(B-C)** (n=5 flies), and **(D-E)** (n=10 flies).

**Extended Data Fig. 2.**
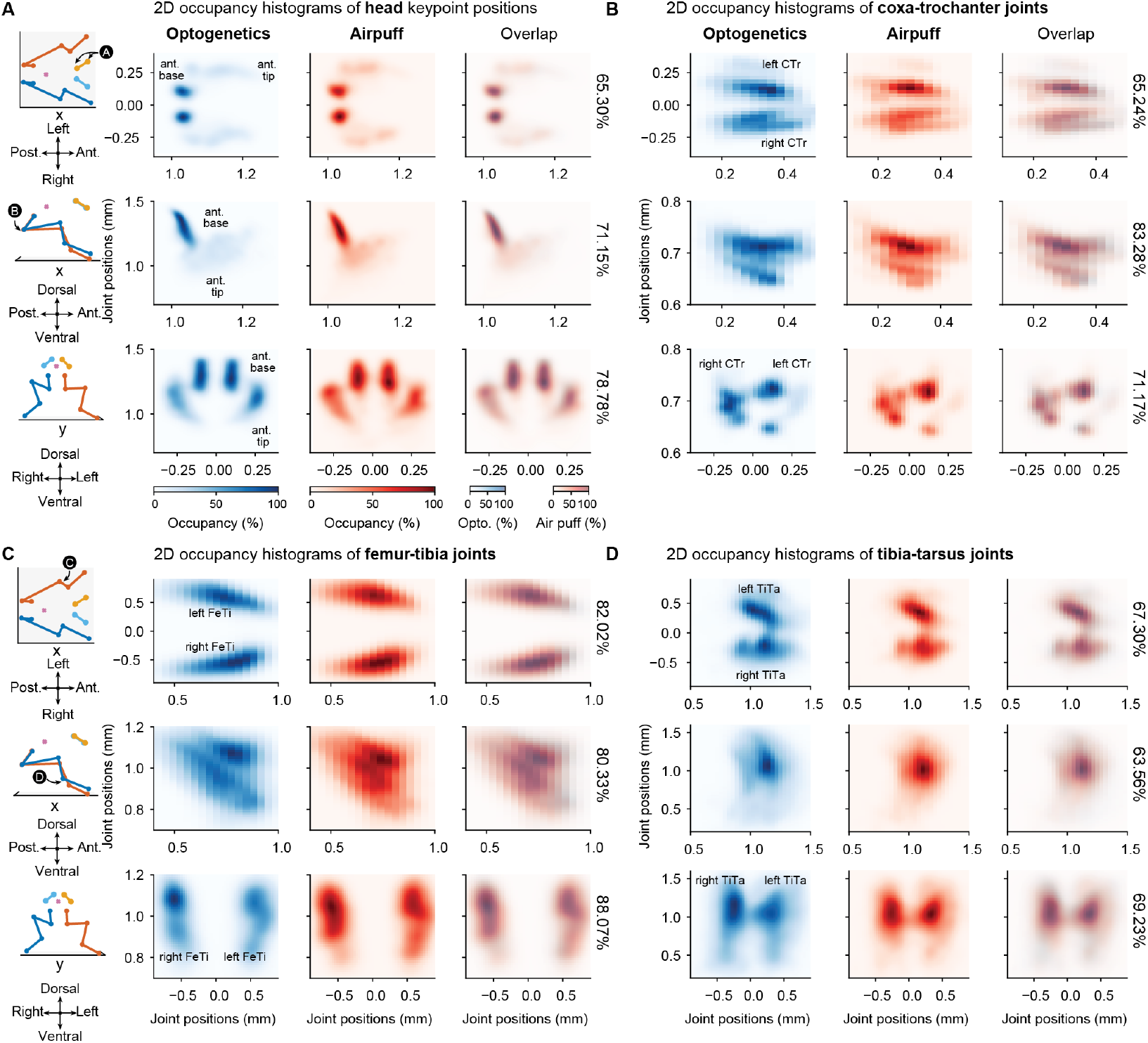
Comparison of optogenetic-versus air puff-elicited antennal grooming kinematics. **(A**,**C, left)** 3D visualizations illustrating the fly’s orientation in neighboring 2D histograms. Key points shown in panel A-D occupancy histograms are indicated (black circles). Histograms display the position occupancy of the left and right: **(A)** antennal bases and tips, **(B)** the coxa-trochanter joints, **(C)** femur-tibia joints, and **(D)** tibia-tarsus joints. **(A-D)** From top to bottom, 2D occupancy histograms show body segment positions in the x-y (top view), x-z (side view), and y-z (front view) planes. From left to right, the histograms illustrate optogenetic-(blue) or air puff-elicited (red) antennal grooming kinematics, as well as the overlap between the two. Each 2D histogram represents the frequency of a body part’s presence in each spatial location. Darker colors indicate higher occupancy. Overlaps illustrate an intersection between occupied areas. Indicated is the precise percent of overlap (right). Data are taken from n=5 flies.

**Extended Data Fig. 3.**
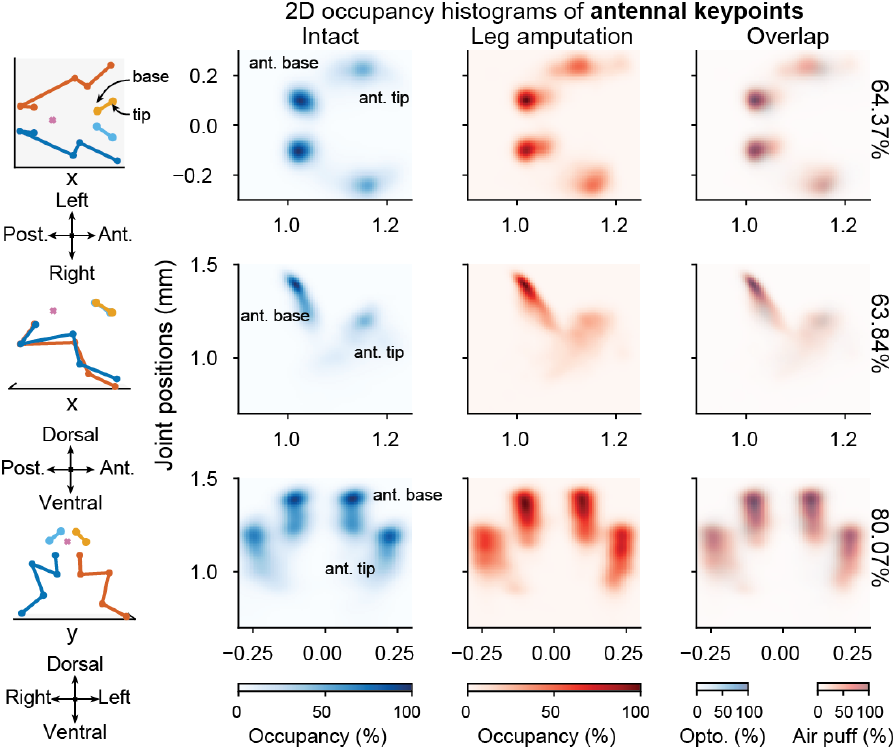
Comparison of optogenetically-elicited antennal grooming kinematics in intact versus foreleg amputee animals. **(left)** 3D visualizations illustrate the fly’s orientation in neighboring 2D histograms. Key points shown in occupancy histograms are indicated (arrows).**(right)** Histograms display the positional occupancy of the left and right antennal base and tip key points. From top to bottom, 2D occupancy histograms show body segment positions in the x-y (top view), x-z (side view), and y-z (front view) planes. From left to right, the histograms illustrate intact (blue) or leg amputee (red) antennal grooming kinematics, as well as the overlap between the two. Each 2D histogram represents the frequency of a body part’s presence in each spatial location. Darker colors indicate higher occupancy. Overlaps illustrate an intersection between occupied areas. Indicated is the precise percent of overlap (right). Data are taken from n=7 flies.

**Extended Data Fig. 4.**
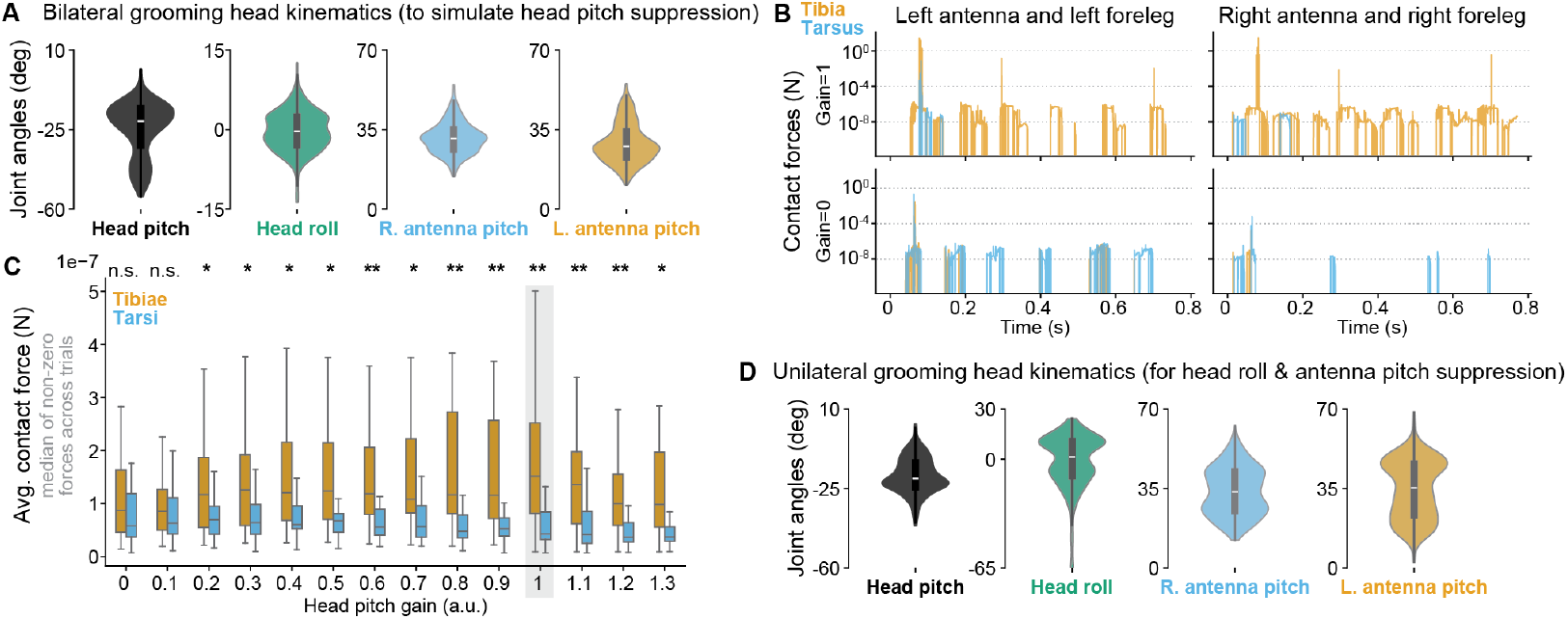
Dataset used for the kinematic replay and quantification of antennal grooming contact forces. **(A)** Violin plots showing the distribution of head and antennal movements during bilateral grooming. These data were used to investigate the effect of downward head pitch in kinematic replay experiments. **(B)** Time series illustrating the contact forces exerted by the left and right foreleg tibial (orange) or tarsal (blue) segments with their ipsilateral antennae at head pitch gains of 1 (top) or 0 (bottom). **(C)** Box plots summarizing the distribution of contact forces between tibial and tarsal leg segments and the antennae as a function of head pitch gain. Box plots show the median of each trial’s non-zero contact forces. Statistics compare tibial and tarsal contact force distributions at a single gain value using a two-sided Mann-Whitney U test. **(D)** Violin plots showing the distribution of head and antennal movements during unilateral antennal grooming. These data were used to study the impact of head roll and antennal pitch suppression. Significance levels are as follows: ***: *P <* 0.001, **: *P <* 0.01, *: *P <* 0.05 and not significant (NS): *P* ≥ 0.05.

**Extended Data Fig. 5.**
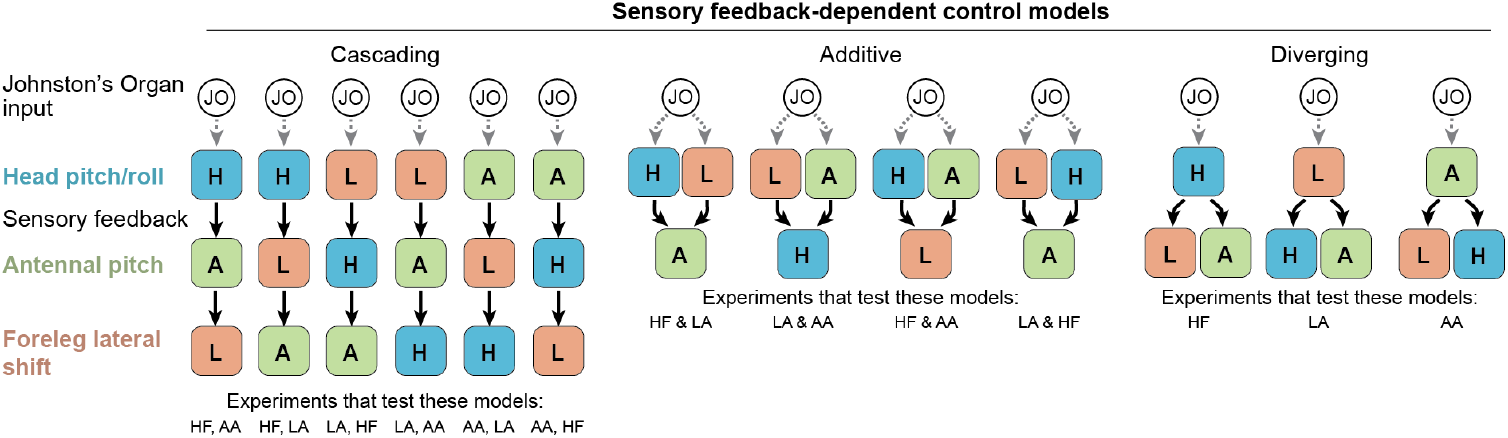
Diagrams of proprioceptive sensory feedback control models. Each colored block represents a motor module consisting of motor neurons and their premotor partners driving a particular body part degree of freedom. For each model, all configurations are shown. In *cascading coordination*, proprioceptive sensory feedback from the first moving body part drives movements of the following body parts. In *additive coordination*, feedback from the first two moving body parts jointly drive movements of the third. In *diverging coordination*, feedback from one body part drives the movements of the other two.

**Extended Data Fig. 6.**
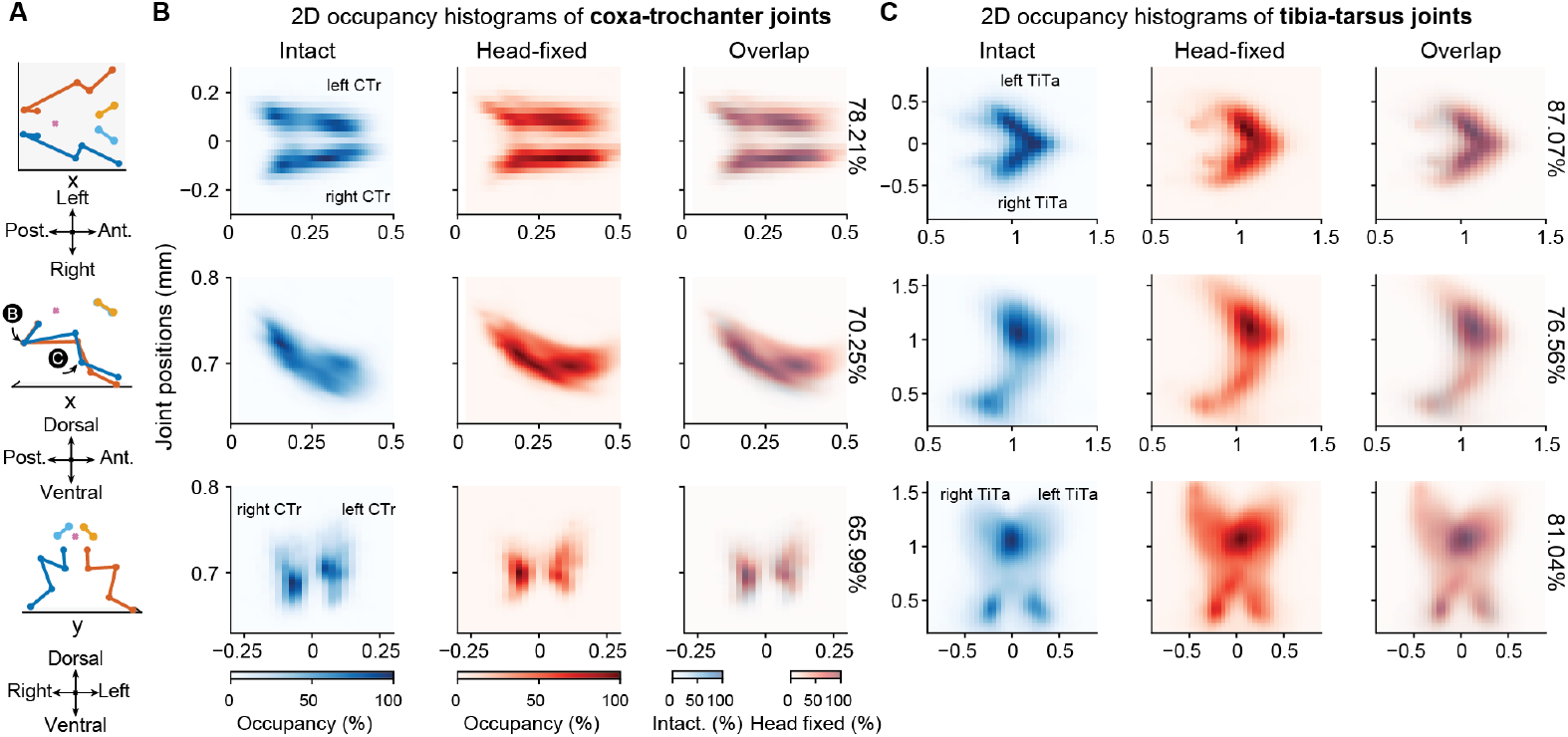
Spatial distribution of foreleg keypoint positions in intact versus head-fixed animals during grooming. **(A)** 3D visualizations illustrate the fly’s orientation in neighboring 2D histograms. Key points shown in panel B-C occupancy histograms are indicated (black circles). Histograms display the positional occupancy of the left and right **(B)** coxa-trochanter joints and **(C)** tibia-tarsus joints. From top to bottom, 2D occupancy histograms show body segment positions in the x-y (top view), x-z (side view), and y-z (front view) planes. From left to right, the histograms illustrate intact (blue) or head-fixed (red) antennal grooming kinematics, as well as the overlap between the two. Each 2D histogram represents the frequency of a body part’s presence in each spatial location. Darker colors indicate higher occupancy. Overlaps illustrate the intersection between occupied areas. Indicated is the percent of overlap (right). Data are from n=9 flies.

**Extended Data Fig. 7.**
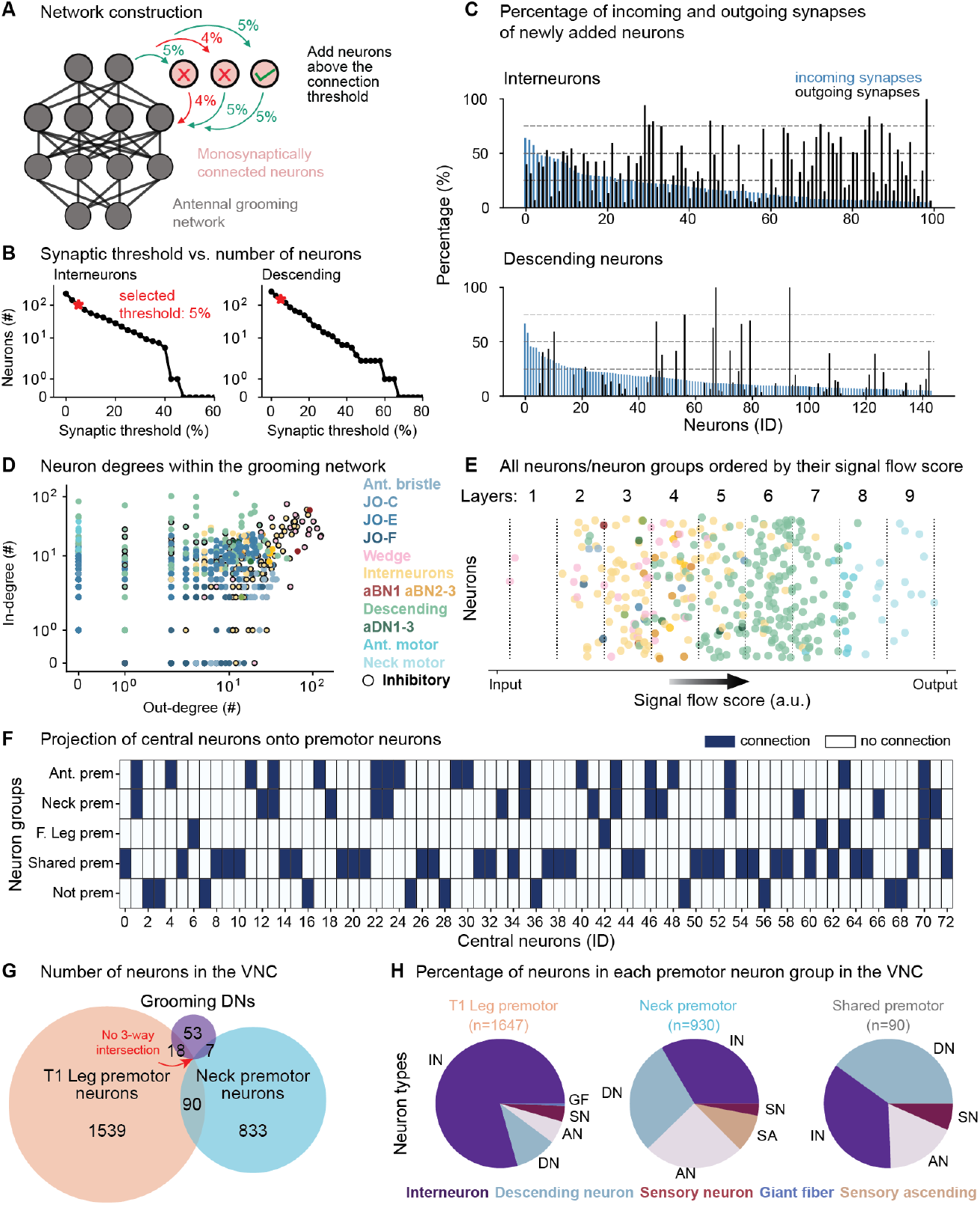
Construction and characterization of the connectome-derived antennal grooming network in the brain and VNC. **(A)** To construct our antennal grooming network, we identified all neurons that are monosynaptically connected (light pink circles) to previously identified antennal grooming neurons (dark gray circles) in the brain. Neurons with both presynaptic and postsynaptic connections exceeding a threshold (green connections) were included in this new network. **(B)** The number of interneurons and descending neurons included as a function of the synaptic percentage threshold. The selected thresholds for network construction are indicated (red asterisks). **(C)** Percentage of incoming (blue) and outgoing (black) synapses for newly included interneurons (top) and descending neurons (bottom) using the selected threshold. No outgoing synapse threshold was applied to descending neurons because they often lack substantial outputs (i.e., axon terminals) in the brain. **(D)** Inversus out-degree for neurons in our constructed antennal grooming network. Neurons are color-coded by type. Inhibitory neurons are indicated (encircled in black). Note the logarithmic scales. **(E)** All neurons ordered by their signal flow score. The signal flow axis is divided into nine equal intervals representing layers from input (left) to output (right). Each circle represents an individual neuron except for sensory neurons (JO-C/E/F and ant. bristles), which are grouped based on their cell type. **(F)** Heatmap illustrating the projections of central neurons onto various premotor types, including antennal, neck, foreleg, and shared premotor neurons. Dark blue squares indicate that the central neuron is a presynaptic partner to the premotor neuron in the corresponding row. **(G)** Venn diagram illustrating the classification of descending neurons in the brain antennal grooming network (purple) as being also either VNC foreleg premotor (orange), or VNC neck premotor (blue). Note that no descending neurons are classified as both foreleg and neck premotor. **(H)** Pie charts showing the percentage of neuron types in each premotor neuron group in the VNC for T1 leg premotor (left), neck premotor (center), and shared premotor neurons (right). The proportion of descending neurons is highest among shared premotor neurons.

**Extended Data Fig. 8.**
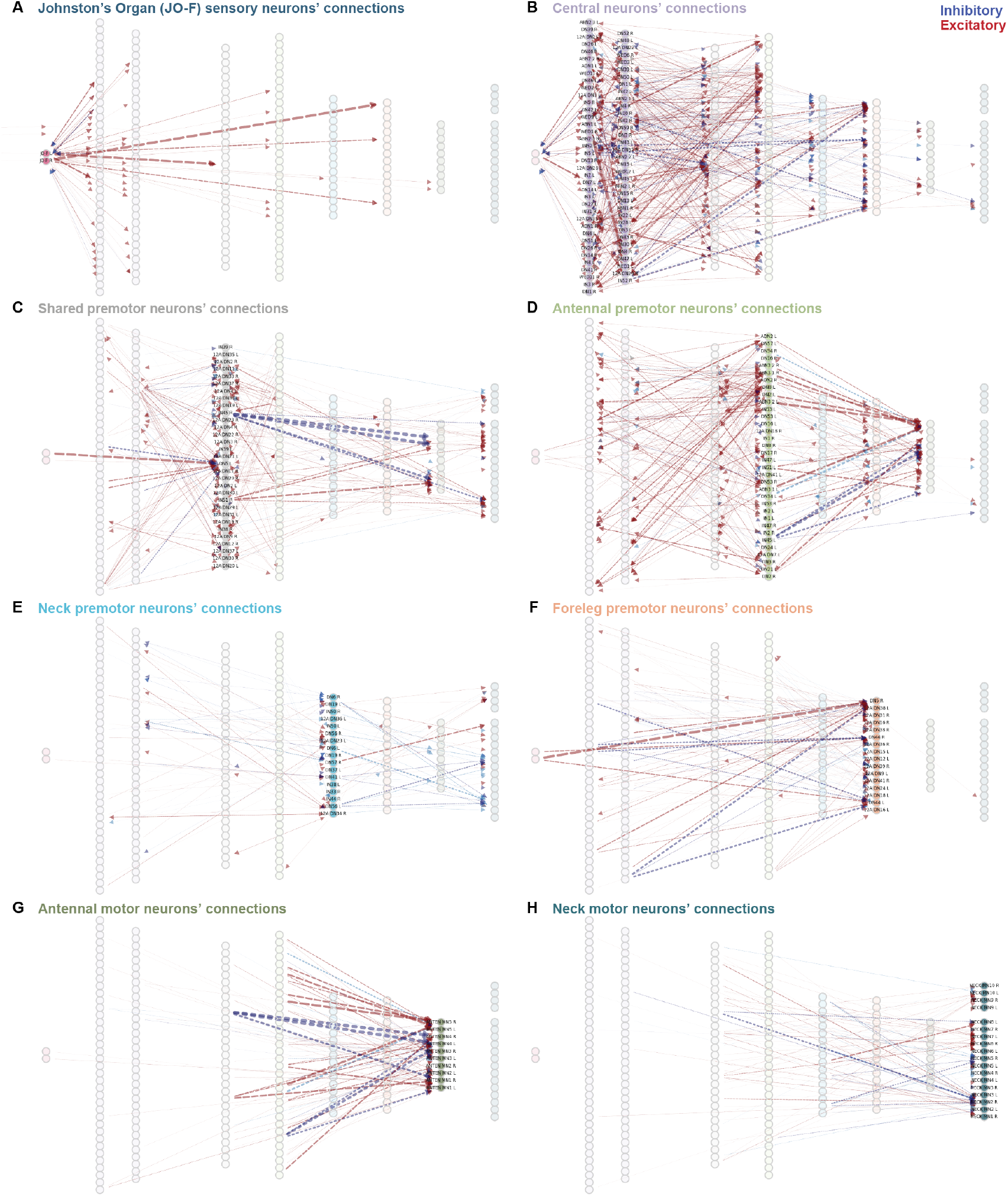
Connectivity of different neuron types in the antennal grooming network. Illustrated are connections to other network neurons by **(A)** Johnston’s Organ sensory inputs, **(B)** central neurons, **(C)** shared premotor neurons, **(D)** antennal premotor neurons, **(E)** neck premotor neurons, **(F)** leg premotor neurons, **(G)** antennal motor neurons, and **(H)** neck motor neurons. High-lighted in each panel are the neurons of interest. Neurotransmitter types are color coded: inhibitory (blue; GABAergic or glutamatergic), excitatory (red; cholinergic), and other neurotransmitter (grey; e.g., dopaminergic). Line widths are proportional to synaptic count.

**Extended Data Fig. 9.**
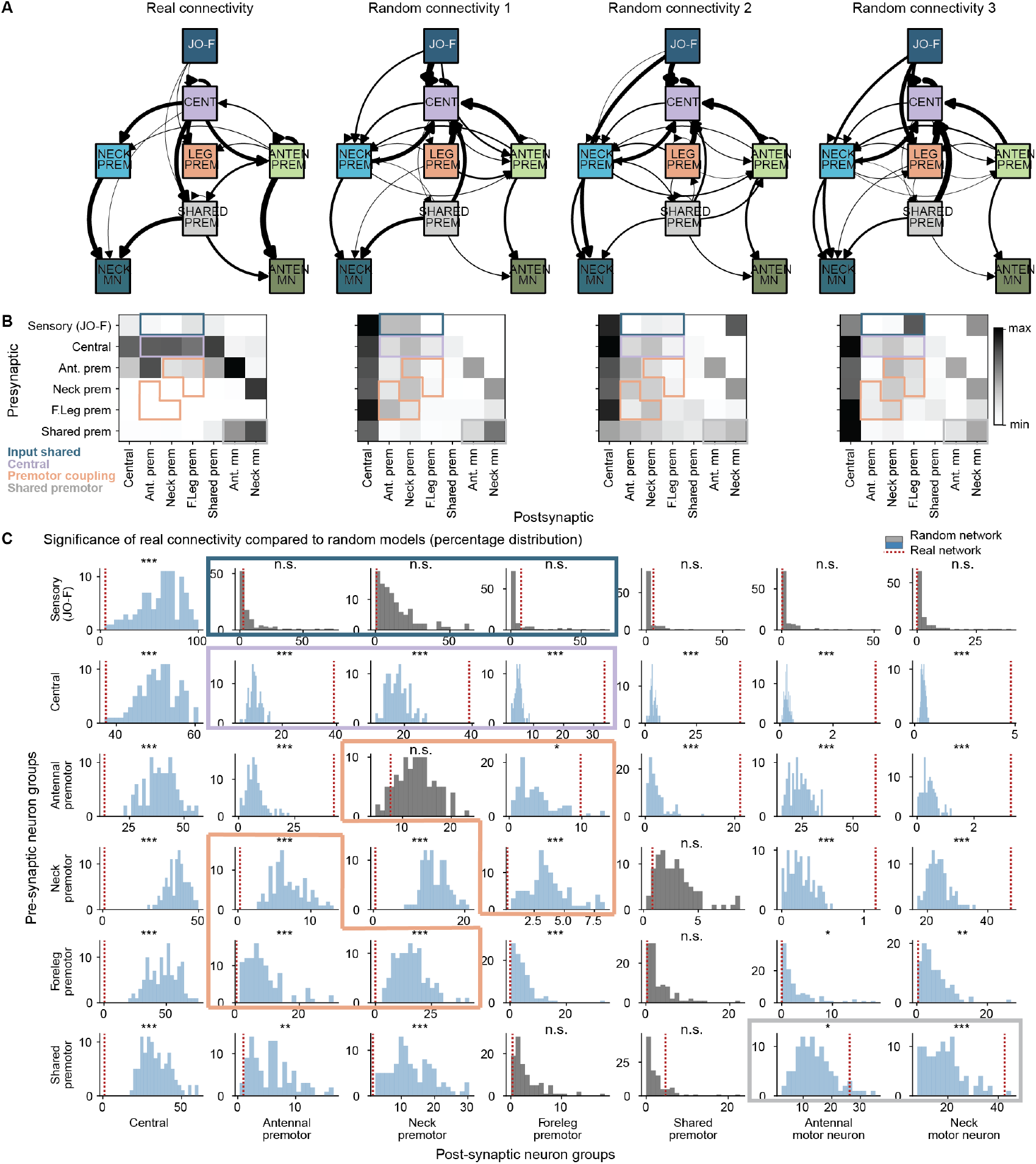
Connectivity between neuron groups in real and randomly shuffled grooming networks. **(A)** Graph representations of connectivity between neuron groups in real and randomized networks. Line widths indicate the percentage of connectivity between groups, with connections below 5% of the maximum strength omitted. The far left shows the real connectivity. The remaining three are examples of randomly shuffled networks. **(B)** Heatmap displaying input contributions between neuron groups. The color scale is normalized within each heatmap. Heatmaps are each taken from the corresponding graph representation (above) in panel A. **(C)** Percentage of connections in real (red dashed line) and randomized (histogram) networks. Randomized network distributions that are significantly different from the real network are colored light blue; non-significant ones are colored gray. Significance levels are as follows: ***: percentile 1 or 99; **: percentile 2.5 or 97.5; *: percentile 5 or 95; and not significant (NS): otherwise. **(B**,**C)** Anticipated connections for each hypothetical model are outlined by colored boxes.

**Extended Data Fig. 10.**
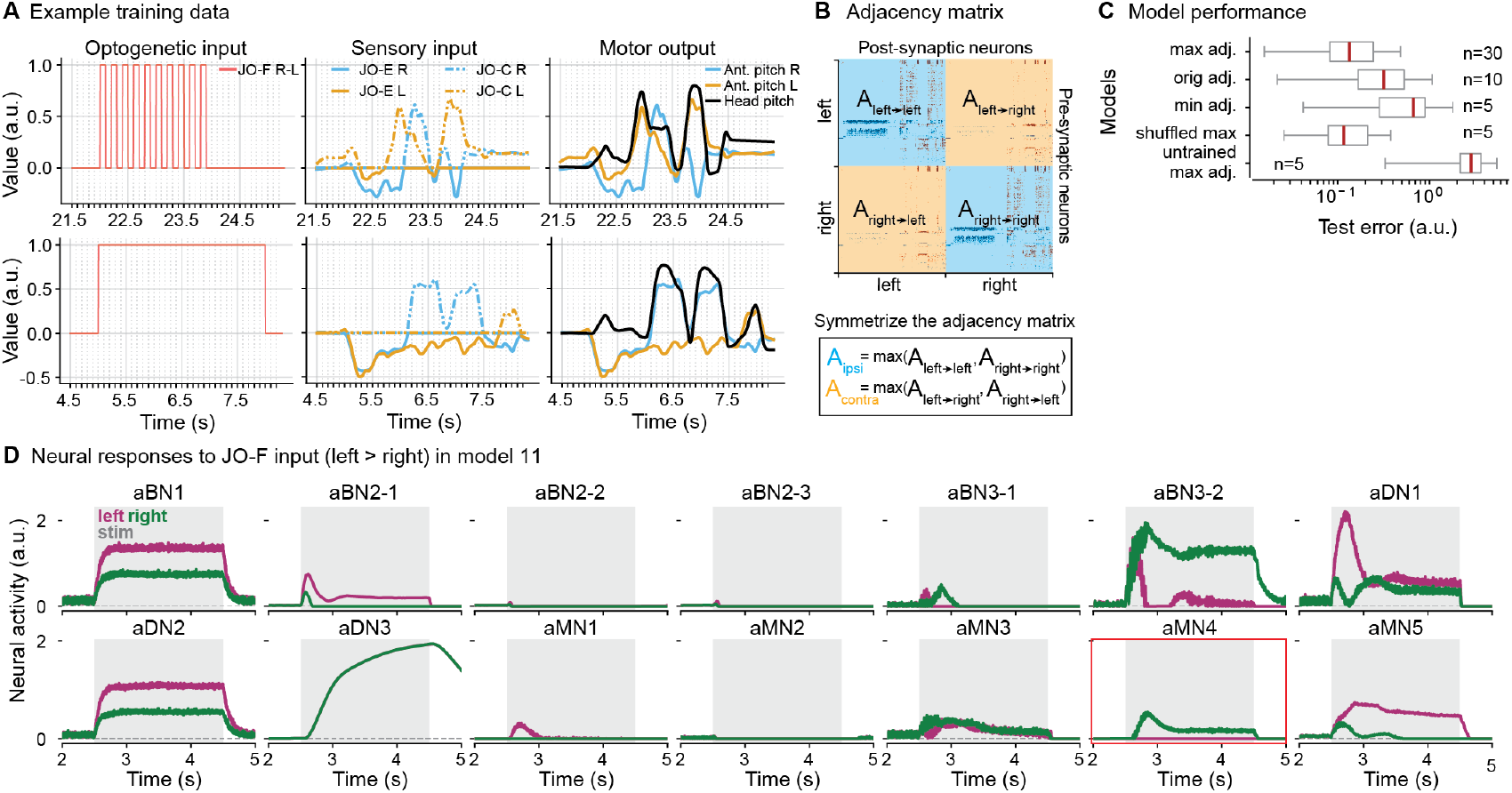
Training and analysis of connectome-derived artificial neural networks. **(A)** Example experimental trials showing an input-output pair from the training dataset with a 5 Hz pulsatile input (top) and a 3 s step input (bottom). Optogenetic stimuli delivered to JO-F neurons mirror those used in real experiments. In addition to this optogenetic input, we also provided JO-CE neurons with fictive sensory feedback. To do this, we processed the antennal pitch motor output to separate upward (JO-C) and downward (JO-E) antennal movements, and then added a 40 ms sensorimotor delay. The unprocessed motor output served as the decoder’s output. **(B)** We symmetrized the network’s adjacency matrix by setting the connections between the two hemispheres to the maximum observed value. This was applied separately for ipsilateral (blue) and contralateral (orange) connections. **(C)** Test errors for connectome-derived neural network models trained using various adjacency matrices to evaluate the effects of different connectivity structures on network performance. *Max adjacency* represents fully symmetrized networks, where connections between ipsilateral and contralateral neuron pairs were set to their maximum observed values (as shown in panel B). *Original adjacency* refers to the non-symmetrized, original connectivity matrix. *Min adjacency* denotes fully symmetrized networks, but with connections set to their minimum observed values. *Shuffled max adjacency* represents symmetrized networks where neuronal connections were randomized to disrupt anatomical specificity while preserving neurotransmitter identity. *Untrained max adjacency* refers to symmetrized networks with maximum connectivity values but without training. The number of models trained for each condition is indicated next to each box plot. Box plots show the median, quartiles, and whiskers extending up to 1.5 times the interquartile range (IQR). Lower test errors indicate better performance, with symmetrized networks generally outperforming original or sparsified counterparts. **(D)** Activities of antennal brain interneurons (aBNs), descending neurons (aDNs), and motor neurons (aMNs) (left/magenta, right/green) from model 11 when the left JO-F input is slightly higher than the right one. aMN4 is indicated (red outline). Gray areas indicate the JO-F stimulation period. Voltage traces are processed through an activation function (rectified linear unit or ReLU).

**Extended Data Fig. 11.**
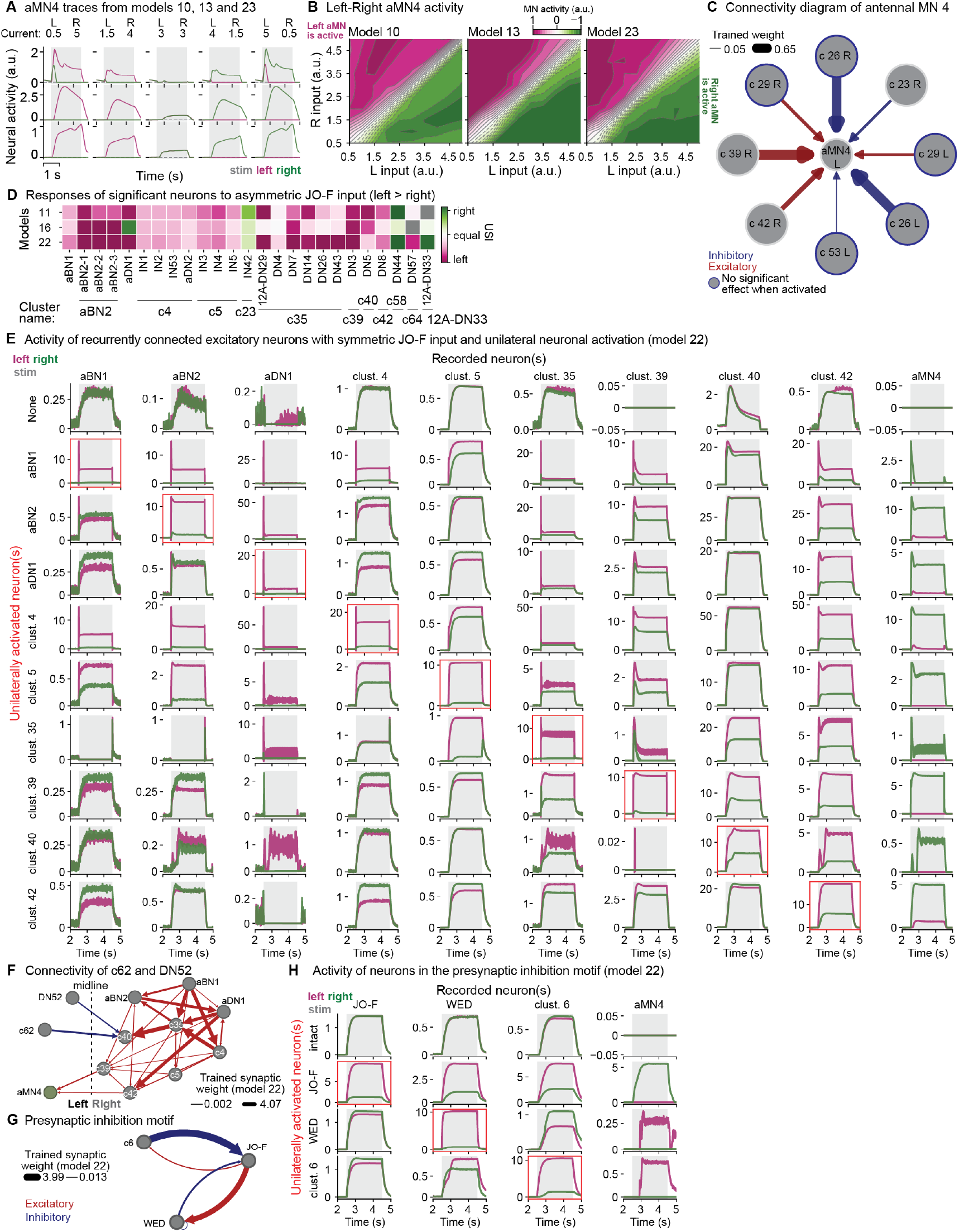
The connectivity of antennal MN4, activity dynamics of neuron clusters in the intact network, and activity dynamics in the presynaptic inhibition motif. **(A)** Left (magenta) and right (green) aMN4 activity traces for different JO-F input pairs across models 10, 13, and 23. The JO-F stimulation period is shaded in gray. During asymmetric JO-F input, the contralateral aMN4 responds, while the ipsilateral aMN4 shows diminished or subthreshold activity (i.e., below zero). **(B)** Activities of aMN4 on the left or right side of the brain. These are shown as a function of the input current magnitudes to the left and right JO-F in the intact network. Values represent the difference between the area under the curve of left and that of right motor neuron activity (magenta for left MN-dominant, green for right MN-dominant). Solid lines mark positive intervals, and dashed lines mark negative intervals, in increments of 0.1. Neither motor neuron dominates along and around the diagonal (white). **(C)** Connectivity of aMN4. Neurons outlined in dark blue did not significantly affect aMN4 activity when unilaterally activated. **(D)** Responses of neurons within motifs (**Fig. 6H-I**) to asymmetric JO-F input (left*>*right) for models 11, 16, and 22. Each column represents one neuron, grouped by cluster (horizontal lines and cluster names). Neural responses were quantified using the USI response metric. Grey squares indicate no activity in both neurons (USI = 0/0). Magenta and green squares denote ipsilateral and contralateral responses, respectively, with darker shades indicating fully unilateral activity. **(E)** Simulated neural dynamics of neurons/clusters in the recurrent excitation network motif (**Fig. 6H**). Each row corresponds to a unilaterally (left) activated neuron (boxed in red) during bilaterally symmetric JO-F input. Each column shows activity of neurons (left/magenta; right/green). JO-F and neuron stimulation periods are shaded in gray. **(F)** Diagram illustrating connections between the recurrent excitation motif and the inhibitory neurons DN52 and c62. Dashed vertical line separates the left and right hemispheres. **(G)** Diagram illustrating the presynaptic inhibition motif between inhibitory clusters and JO-F neurons. Neurons from only one hemisphere are shown. **(C**,**F**,**G)** Red and blue lines represent excitatory and inhibitory connections, respectively, with line thickness proportional to the trained weights from model 22. **(H)** Simulated neural dynamics of neurons/clusters in the presynaptic inhibition network motif (panel G). Each row corresponds to a unilaterally (left) activated neuron (boxed in red) during bilaterally symmetric JO-F input. Each column shows activity of neurons (left/magenta; right/green). JO-F and neuron stimulation periods are shaded in gray. **(A**,**E**,**H)** Voltage traces are processed through an activation function (rectified linear unit or ReLU). For clusters containing multiple neuron pairs, average neural activity is shown.

**Extended Data Fig. 12.**
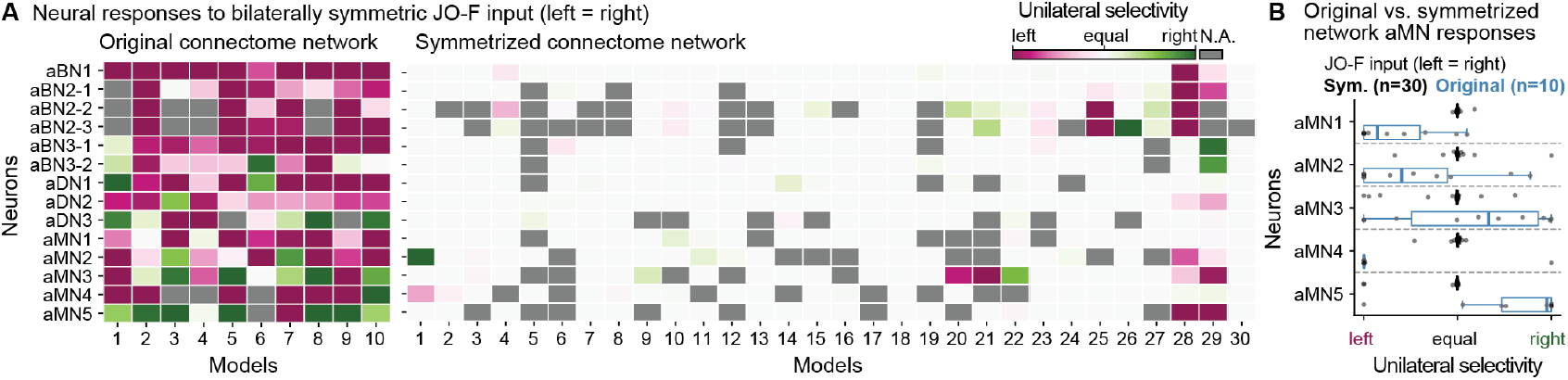
Neural responses of models trained with the original (non-symmetrized) adjacency matrix. **(A)** Responses of antennal brain interneurons (aBNs), descending neurons (aDNs), and motor neurons (aMNs) to bilaterally symmetric JO-F input (left = right) in trained models using **(left)** the original, non-symmetrized connectome network (*n* = 10) or **(right)** the symmetrized connectome network (*n* = 30). Neural responses were quantified using the USI response metric. Grey squares indicate zero neural activity in both neurons (USI = 0/0). Magenta and green squares represent neurons responding more to stimulation of their ipsilateral or contralateral JO-F, respectively. Darker colors indicate that only one neuron is active. **(B)** Response types of motor neurons for bilaterally symmetric JO-F input across models using the original non-symmetric connectome network (blue) or the symmetrized connectome network (black) as adjacency matrices. Each dot represents a model (corresponding to a square in panel A). Box plots display medians and quartiles, while whiskers extend to the full distribution, excluding outliers beyond 1.5 times the interquartile range (IQR).

## Supplementary Information Files

**Supplementary Information File:** Exact p-values for statistical tests performed in this study (Ref. Fig. **2**, Fig. **3**, and **Extended Data Fig**. 4). Excel file containing FlyWire IDs, names, modules. Clusters of neurons that constitute the antennal grooming network (Ref. Fig. **4**.) Link to Supporting Information File

### Supplementary Videos

**Supplementary Video 1: Behavioral recordings, 3D pose estimation, inverse kinematics, and joint angles during antennal grooming**. Experimental recordings (left, top and middle rows) were used to estimate the fly’s 3D pose (left, bottom row). Inverse kinematics calculations allowed us to derive joint angles for the head and antennae (right, top row) as well as for the forelegs (right, second and third rows). Indicated are the onset and offset of optogenetic stimulation (red circles in camera images, red vertical lines on plots). Here and elsewhere, the fly genotype is *20xUAS-CsChrimson; +* ; *GMR60E02-GAL4*. Video and data are shown at 0.5x real-time.

Link to Supplementary Video 1

**Supplementary Video 2: Behavioral recordings, 3D pose estimation, inverse kinematics, and kinematic replay in a biomechanical model during antennal grooming**. Shown is the original video (left), 3D pose estimation (middle, solid lines), inverse kinematics (middle, dashed lines), and kinematic replay in NeuroMechFly, a biomechanical fly simulation (right). Video and data are shown at 0.25x real-time.

Link to Supplementary Video 2

**Supplementary Video 3: Behavioral classification of optogenetically-elicited antennal grooming**. Videos of four optogenetic stimulation trials for six flies. Overlaid are seven behavior classification labels: ‘bilateral’, unilateral tripartite (‘unilateral t right, or left’), partial unilateral non-tripartite (‘unilateral nt right, or left’), non-classified (‘nc’), and ‘background’. Indicated are the onset and offset of optogenetic stimulation (red circles in camera images). Video and data are shown at 0.5x real-time.

Link to Supplementary Video 3

**Supplementary Video 4: Comparison of air puff-versus optogenetic stimulation-elicited antennal grooming**. Videos of two air puff (top) and optogenetic stimulation (bottom) trials for three individual flies. Each fly is numbered. Glass capillary for air puff stimulation is on the left. Indicated are the onset and offset of the air puff (blue circles in camera images) or optogenetic stimulus (red circles in camera images). Video and data are shown at 0.5x real-time.

Link to Supplementary Video 4

**Supplementary Video 5: Kinematic replay of intact versus computationally perturbed antennal grooming**. Biomechanical simulation kinematic replay in NeuroMechFly of intact (top, ‘Gain=1’), or perturbed (bottom, ‘Gain=0’) inverse kinematics-derived antennal grooming. Grooming subtypes are bilateral (left) or unilateral (middle and right). In each column one degree of freedom is perturbed: head pitch (left), head roll (middle), or antennal pitch (right). For head pitch and head roll, Gain=1 indicates the original joint angles, while Gain=0 indicates no movement. For antennal pitch, Gain=1 indicates 60°upward pitch and Gain=0 indicates a resting pose (no pitch) at 10°. Data are replayed at 0.1× real-time.

Link to Supplementary Video 5

**Supplementary Video 6: Behavioral recordings of optogenetically-elicited antennal grooming in flies before versus after head fixation**. Videos of two optogenetic stimulation trials for three individual flies either before (top) or after (bottom) head fixation. Each fly is numbered. Indicated are the onset and offset of optogenetic stimulation (red circles in camera images). Video and data are shown at 0.5x real-time.

Link to Supplementary Video 6

**Supplementary Video 7: Behavioral recordings of optogenetically-elicited antennal grooming in flies before versus after foreleg amputation**. Videos of two optogenetic stimulation trials for three individual flies either before (top) or after (bottom) leg amputation. Each fly is numbered. Indicated are the onset and offset of optogenetic stimulation (red circles in camera images). Video and data are shown at 0.5x real-time.

Link to Supplementary Video 7

**Supplementary Video 8: Behavioral recordings of optogenetically-elicited antennal grooming in flies before versus after amputation of their antennae**. Videos of two optogenetic stimulation trials for three individual flies either before (top) or after (bottom) amputation of their antennae. Each fly is numbered. Indicated are the onset and offset of optogenetic stimulation (red circles in camera images). Video and data are shown at 0.5x real-time.

Link to Supplementary Video 8

**Supplementary Video 9: Behavioral recordings of optogenetically-elicited antennal grooming in flies before perturbation, after amputation of their forelegs, and then also following head immobilization**. Videos of two optogenetic stimulation trials for three individual flies either before any perturbation (top), after amputation of their forelegs (middle), and after head immobilization as well (bottom). Each fly is numbered. Indicated are the onset and offset of optogenetic stimulation (red circles in camera images). Video and data are shown at 0.5x real-time.

Link to Supplementary Video 9

**Supplementary Video 10: Behavioral recordings of optogenetically-elicited antennal grooming in flies before perturbation, after amputation of their antennae, and then also following foreleg amputation**. Videos of two optogenetic stimulation trials for three individual flies either before any perturbation (top), after amputation of their antennae (middle), and after amputation of their forelegs as well (bottom). Each fly is numbered. Indicated are the onset and offset of optogenetic stimulation (red circles in camera images). Video and data are shown at 0.5x real-time.

Link to Supplementary Video 10

**Supplementary Video 11: Behavioral recordings of optogenetically-elicited antennal grooming in flies before perturbation, after amputation of their antennae, and then also following head immobilization**. Videos of two optogenetic stimulation trials for three individual flies either before any perturbation (top), after amputation of their antennae (middle), and after head immobilization as well (bottom). Each fly is numbered. Indicated are the onset and offset of optogenetic stimulation (red circles in camera images). Video and data are shown at 0.5x real-time.

Link to Supplementary Video 11

**Supplementary Video 12: Animation of network dynamics in intact, WED-, and c6-silenced networks for models 11, 16, and 22**. Network dynamics in (left) intact, (middle) WED-silenced, and (right) c6-silenced networks in response to bilaterally symmetric JO-F input. Shown are networks from models (top) 11, (center) 16, and (bottom) 22 are shown. JO-F stimulation begins at 400 ms and ends at 2400 ms. Circles represent clusters. Indicated are inhibitory clusters (black outline). Node colors are proportional to non-normalized neural activity: red indicates depolarization, blue indicates hyperpolarization, and white indicates neurons at rest. For clusters containing multiple neurons, the average activity is displayed. The left and right halves of each panel correspond to the left and right hemispheres of the network. The title shows time points.

Link to Supplementary Video 12

**Supplementary Video 13: Animation of network dynamics in intact, unperturbed models 11, 16, and 22**. Intact network dynamics in response to (left) bilaterally symmetric, or (right) asymmetric JO-F (right *>* left) input. Shown are dynamics in models (top) 11, (center) 16, and (bottom) 22 are shown. JO-F stimulation begins at 400 ms and ends at 2400 ms. Circles represent clusters. Indicated are inhibitory clusters (black outline). Node colors are proportional to non-normalized neural activity: red indicates depolarization, blue indicates hyperpolarization, and white indicates neurons at rest. For clusters containing multiple neurons, the average activity is displayed. The left and right halves of each panel correspond to the left and right hemispheres of the network. The title shows time points.

Link to Supplementary Video 13

## Acknowledgments

We thank Stefanie Hampel and Andrew Seeds for helpful discussions and sharing the identification of antennal grooming neurons in FAFB; Janne Lappalainen for assistance in using the Flyvis package; Melissa Faggella and Olivia Caroline Ruggaber for 2D pose estimation annotations; Stefanie Boy-Röttger and Maite Azcorra for fly dissections and confocal imaging; Jasper Phelps for help with EM datasets; Kathi Eichler and Gregory Jefferis for sharing comprehensive proofreading and annotation of DNs in the FAFB and FANC datasets before publication; Marie Suver for helpful discussions on the identification of antennal motor neurons; Stephen Huston for sharing neck motor neuron identification data from FANC and MANC before publication. We thank members of the Biorobotics and Neuroengineering Laboratory for helpful discussions and invaluable feedback on the manuscript. We thank Brian McCabe (EPFL, Lausanne, Switzerland) for transgenic *Drosophila* strains. Stocks obtained from the Bloomington Drosophila Stock Center (NIH P40OD018537) were used in this study. PR acknowledges support from an SNSF Project Grant (175667) and an SNSF Eccellenza Grant (181239). JA acknowledges support from a European Research Council Synergy grant (951477). PGÖ acknowledges support from a Swiss Government Excellence Scholarship for Doctoral Studies and a Google PhD Fellowship.

## Author Contributions

P.G.O. - Conceptualization, Methodology, Software, Investigation, Formal Analysis, Data Acquisition, Data Curation, Visualization, Writing – Original Draft Preparation, Writing - Review & Editing.

J.A. - Methodology, Software, Investigation, Visualization, Writing - Review & Editing.

C.S. - Methodology, Investigation, Formal Analysis, Data Curation, Writing - Review & Editing.

A.J.I. - Conceptualization, Methodology, Investigation, Resources, Writing - Review & Editing, Supervision, Project Administration, Funding Acquisition.

P.R. - Conceptualization, Methodology, Resources, Writing – Original Draft Preparation, Writing - Review & Editing, Supervision, Project Administration, Funding Acquisition.

## Ethical compliance

All experiments were performed in compliance with relevant national (Switzerland) and institutional (EPFL) ethical regulations.

## References

[1] Bidaye, S. S., Bockemühl, T. & Büschges, A. Six-legged walking in insects: how CPGs, peripheral feedback, and descending signals generate coordinated and adaptive motor rhythms. Journal of Neurophysiology 119, 459–475 (2018).

[2] Dickinson, M. H. et al. How Animals Move: An Integrative View. Science 288, 100–106 (2000).

[3] Ruder, L. & Arber, S. Brainstem circuits controlling action diversification. Annual review of neuroscience 42, 485–504 (2019).

[4] Skinner, F. K. & Mulloney, B. Intersegmental coordination in invertebrates and vertebrates. Current Opinion in Neurobiology 8, 725–732 (1998).

[5] Ijspeert, A. J. & Daley, M. A. Integration of feedforward and feedback control in the neuromechanics of vertebrate locomotion: a review of experimental, simulation and robotic studies. Journal of Experimental Biology 226 (2023).

[6] Kiehn, O. Decoding the organization of spinal circuits that control locomotion. Nature Reviews Neuroscience 17, 224–238 (2016).

[7] Grillner, S. & El Manira, A. Current Principles of Motor Control, with Special Reference to Vertebrate Locomotion. Physiological Reviews 100, 271–320 (2020).

[8] Lanuza, G. M., Gosgnach, S., Pierani, A., Jessell, T. M. & Goulding, M. Genetic identification of spinal interneurons that coordinate left-right locomotor activity necessary for walking movements. Neuron 42, 375–386 (2004).

[9] Gabriel, J. P. et al. Principles governing recruitment of motoneurons during swimming in zebrafish. Nature neuroscience 14, 93–99 (2011).

[10] Talpalar, A. E. et al. Dual-mode operation of neuronal networks involved in left–right alternation. Nature 500, 85–88 (2013).

[11] Zelenin, P. V. et al. Differential contribution of v0 interneurons to execution of rhythmic and nonrhythmic motor behaviors. Journal of Neuroscience 41, 3432–3445 (2021).

[12] Arshavsky, Y. I., Orlovsky, G., Panchin, Y. V., Roberts, A. & Soffe, S. Neuronal control of swimming locomotion: analysis of the pteropod mollusc clione and embryos of the amphibian xenopus. Trends in neurosciences 16, 227–233 (1993).

[13] Wilson, A. C. & Sweeney, L. B. Spinal cords: Symphonies of interneurons across species. Frontiers in Neural Circuits 17, 1146449 (2023).

[14] Ryczko, D., Simon, A. & Ijspeert, A. J. Walking with salamanders: from molecules to biorobotics. Trends in neurosciences 43, 916–930 (2020).

[15] Büschges, A., Akay, T., Gabriel, J. P. & Schmidt, J. Organizing network action for locomotion: Insights from studying insect walking. Brain Research Reviews 57, 162–171 (2008).

[16] Pick, S. & Strauss, R. Goal-driven behavioral adaptations in gap-climbing drosophila. Current Biology 15, 1473–1478 (2005).

[17] Muijres, F. T., Elzinga, M. J., Melis, J. M. & Dickinson, M. H. Flies evade looming targets by executing rapid visually directed banked turns. Science 344, 172–177 (2014).

[18] Seeds, A. M. et al. A suppression hierarchy among competing motor programs drives sequential grooming in Drosophila. eLife 3, e02951 (2014).

[19] Dorkenwald, S. et al. Neuronal wiring diagram of an adult brain. Nature 634, 124–138 (2024).

[20] Schlegel, P. et al. Whole-brain annotation and multi-connectome cell typing of drosophila. Nature 634, 139–152 (2024).

[21] Zheng, Z. et al. A complete electron microscopy volume of the brain of adult drosophila melanogaster. Cell 174, 730–743 (2018).

[22] Scheffer, L. K. et al. A connectome and analysis of the adult Drosophila central brain. eLife 9, e57443 (2020).

[23] Phelps, J. S. et al. Reconstruction of motor control circuits in adult Drosophila using automated transmission electron microscopy. Cell 184, 759–774.e18 (2021).

[24] Azevedo, A. et al. Connectomic reconstruction of a female drosophila ventral nerve cord. Nature p1–9 (2024).

[25] Takemura, S. et al. A connectome of the male drosophila ventral nerve cord. eLife (2024).

[26] Meissner, G. W. et al. A searchable image resource of Drosophila GAL4 driver expression patterns with single neuron resolution. eLife 12, e80660 (2023).

[27] Simpson, J. H. & Looger, L. L. Functional Imaging and Optogenetics in Drosophila. Genetics 208, 1291–1309 (2018).

[28] Li, J. et al. A defensive kicking behavior in response to mechanical stimuli mediated by drosophila wing margin bristles. Journal of Neuroscience 36, 11275–11282 (2016).

[29] Kalueff, A. V. et al. Neurobiology of rodent self-grooming and its value for translational neuroscience. Nature Reviews Neuroscience 17, 45–59 (2016).

[30] Sachs, B. D. The Development of Grooming and Its Expression in Adult Animals. Annals of the New York Academy of Sciences 525, 1–17 (1988).

[31] Zhukovskaya, M., Yanagawa, A. & Forschler, B. T. Grooming behavior as a mechanism of insect disease defense. Insects 4, 609–630 (2013).

[32] Böröczky, K., Wada-Katsumata, A., Batchelor, D., Zhukovskaya, M. & Schal, C. Insects groom their antennae to enhance olfactory acuity. Proceedings of the National Academy of Sciences of the United States of America 110, 3615–3620 (2013).

[33] Wada-Katsumata, A. & Schal, C. Antennal grooming facilitates courtship performance in a group-living insect, the German cockroach Blattella germanica. Scientific Reports 9, 2942 (2019).

[34] Hampel, S., McKellar, C. E., Simpson, J. H. & Seeds, A. M. Simultaneous activation of parallel sensory pathways promotes a grooming sequence in Drosophila. eLife 6, e28804 (2017).

[35] Mueller, J. M., Zhang, N., Carlson, J. M. & Simpson, J. H. Variation and variability in drosophila grooming behavior. Frontiers in behavioral neuroscience 15, 769372 (2022).

[36] Mueller, J. M., Ravbar, P., Simpson, J. H. & Carlson, J. M. Drosophila melanogaster grooming possesses syntax with distinct rules at different temporal scales. PLOS Computational Biology 15, e1007105 (2019).

[37] Hampel, S. et al. Distinct subpopulations of mechanosensory chordotonal organ neurons elicit grooming of the fruit fly antennae. eLife 9, e59976 (2020).

[38] Zhang, N., Guo, L. & Simpson, J. H. Spatial Comparisons of Mechanosensory Information Govern the Grooming Sequence in Drosophila. Current Biology 30, 988–1001.e4 (2020).

[39] Eichler, K. et al. Somatotopic organization among parallel sensory pathways that promote a grooming sequence in drosophila. eLife (2024).

[40] Hampel, S., Franconville, R., Simpson, J. H. & Seeds, A. M. A neural command circuit for grooming movement control. eLife 4, e08758 (2015).

[41] Guo, L., Zhang, N. & Simpson, J. H. Descending neurons coordinate anterior grooming behavior in Drosophila. Current Biology 32, 823–833.e4 (2022).

[42] Zhang, N. & Simpson, J. H. A pair of commissural command neurons induces drosophila wing grooming. IScience 25 (2022).

[43] Syed, D. S., Ravbar, P. & Simpson, J. H. Inhibitory circuits coordinate leg movements during drosophila grooming. bioRxiv 2024–06 (2024).

[44] Yoshikawa, S., Tang, P. & Simpson, J. H. Mechanosensory and command contributions to the Drosophila grooming sequence. Current Biology 0 (2024).

[45] Ravbar, P., Zhang, N. & Simpson, J. H. Behavioral evidence for nested central pattern generator control of Drosophila grooming. eLife 10, e71508 (2021).

[46] Günel, S. et al. DeepFly3D, a deep learning-based approach for 3D limb and appendage tracking in tethered, adult Drosophila. eLife 8, e48571 (2019).

[47] Karashchuk, P. et al. Anipose: A toolkit for robust markerless 3D pose estimation. Cell Reports 36, 109730 (2021).

[48] Lobato-Rios, V. et al. NeuroMechFly, a neuromechanical model of adult Drosophila melanogaster. Nature Methods 19, 620–627 (2022).

[49] Wang-Chen, S. et al. Neuromechfly v2: simulating embodied sensorimotor control in adult drosophila. Nature Methods 1–10 (2024).

[50] Arreguit, J., Ramalingasetty, S. T. & Ijspeert, A. Farms: Framework for animal and robot modeling and simulation. bioRxiv (2023).

[51] Lappalainen, J. K. et al. Connectome-constrained networks predict neural activity across the fly visual system. Nature 1–9 (2024).

[52] Shiu, P. K. et al. A drosophila computational brain model reveals sensorimotor processing. Nature 634, 210–219 (2024).

[53] Mathis, A. et al. DeepLabCut: markerless pose estimation of user-defined body parts with deep learning. Nature Neuroscience 21, 1281–1289 (2018).

[54] Ozdil, P. G., Ijspeert, A. & Ramdya, P. sequential-inverse-kinematics: v1.0.0 (2024). URL 10.5281/zenodo.12601317.

[55] Dorkenwald, S. et al. Flywire: online community for whole-brain connectomics. Nature methods 19, 119–128 (2022).

[56] Lin, A. et al. Network statistics of the whole-brain connectome of drosophila. Nature 634, 153–165 (2024).

[57] Winding, M. et al. The connectome of an insect brain. Science 379, eadd9330 (2023).

[58] Dallmann, C. J. et al. Presynaptic inhibition selectively suppresses leg proprioception in behaving drosophila. bioRxiv (2023).

[59] Scharstein, H. Input-output relationship of the Leaky-Integrator Neuron Model. Journal of Mathematical Biology 8, 403–420 (1979).

[60] Chen, C. et al. Functional architecture of neural circuits for leg proprioception in Drosophila. Current Biology 31, 5163–5175.e7 (2021).

[61] Chen, C. et al. Ascending neurons convey behavioral state to integrative sensory and action selection brain regions. Nature neuroscience 26, 682–695 (2023).

[62] Pascual, A., Huang, K.-L., Neveu, J. & Préat, T. Brain asymmetry and long-term memory. Nature 427, 605–606 (2004).

[63] Tuthill, J. C. & Wilson, R. I. Mechanosensation and Adaptive Motor Control in Insects. Current Biology 26, R1022–R1038 (2016).

[64] Azevedo, A. W. et al. A size principle for recruitment of Drosophila leg motor neurons. eLife 9, e56754 (2020).

[65] Gorko, B. et al. Motor neurons generate pose-targeted movements via proprioceptive sculpting. Nature 1–8 (2024).

[66] Eckstein, N. et al. Neurotransmitter classification from electron microscopy images at synaptic sites in drosophila melanogaster. Cell 187, 2574–2594 (2024).

[67] Werbos, P. Backpropagation through time: what it does and how to do it. Proceedings of the IEEE 78, 1550–1560 (1990).

[68] Suver, M. P., Medina, A. M. & Nagel, K. I. Active antennal movements in Drosophila can tune wind encoding. Current Biology 33, 780–789.e4 (2023).

[69] Sterne, G. R., Otsuna, H., Dickson, B. J. & Scott, K. Classification and genetic targeting of cell types in the primary taste and premotor center of the adult Drosophila brain. eLife 10, e71679 (2021).

[70] Erginkaya, M. et al. A competitive disinhibitory network for robust optic flow processing in drosophila. bioRxiv 2023–08 (2023).

[71] Feng, K. et al. A central steering circuit in drosophila. bioRxiv (2024).

[72] Machens, C. K., Romo, R. & Brody, C. D. Flexible control of mutual inhibition: a neural model of two-interval discrimination. Science 307, 1121–1124 (2005).

[73] Koyama, M. & Pujala, A. Mutual inhibition of lateral inhibition: a network motif for an elementary computation in the brain. Current opinion in neurobiology 49, 69–74 (2018).

[74] Jovanic, T. et al. Competitive Disinhibition Mediates Behavioral Choice and Sequences in Drosophila. Cell 167, 858–870.e19 (2016).

[75] Mysore, S. P. & Knudsen, E. I. Reciprocal inhibition of inhibition: a circuit motif for flexible categorization in stimulus selection. Neuron 73, 193–205 (2012).

[76] Jing, J. & Gillette, R. Neuronal elements that mediate escape swimming and suppress feeding behavior in the predatory sea slug pleurobranchaea. Journal of neurophysiology 74, 1900–1910 (1995).

[77] Berkowitz, A., Roberts, A. & Soffe, S. R. Roles for multifunctional and specialized spinal interneurons during motor pattern generation in tadpoles, zebrafish larvae, and turtles. Frontiers in behavioral neuroscience 4, 1810 (2010).

[78] Hlavac, T. F. Grooming Systems of Insects: Structure, Mechanics1. Annals of the Entomological Society of America 68, 823–826 (1975).

[79] Rebora, M., Salerno, G., Piersanti, S., Michels, J. & Gorb, S. Structure and biomechanics of the antennal grooming mechanism in the southern green stink bug nezara viridula. Journal of insect physiology 112, 57–67 (2019).

[80] Cheong, H. S. et al. Transforming descending input into behavior: The organization of premotor circuits in the drosophila male adult nerve cord connectome. eLife (2024).

[81] Büschges, A., Schmitz, J. & Bässler, U. Rhythmic patterns in the thoracic nerve cord of the stick insect induced by pilocarpine. Journal of Experimental Biology 198, 435–456 (1995).

[82] Pearson, K. G. Proprioceptive regulation of locomotion. Current Opinion in Neurobiology 5, 786–791 (1995).

[83] Fuchs. Intersegmental coordination of cockroach locomotion: adaptive control of centrally coupled pattern generator circuits. Frontiers in Neural Circuits (2010).

[84] Ayali, A. et al. Sensory feedback in cockroach locomotion: current knowledge and open questions. Journal of Comparative Physiology A 201, 841–850 (2015).

[85] Mendes, C. S., Bartos, I., Akay, T., Márka, S. & Mann, R. S. Quantification of gait parameters in freely walking wild type and sensory deprived Drosophila melanogaster. eLife 2, e00231 (2013).

[86] Chockley, A. S. et al. Subsets of leg proprioceptors influence leg kinematics but not interleg coordination in drosophila melanogaster walking. Journal of Experimental Biology 225 (2022).

[87] Zack, S. The effects of foreleg amputation on head grooming behaviour in the praying mantis,Sphodromantis lineola. Journal of comparative physiology 125, 253–258 (1978).

[88] Berridge, K. C. Progressive degradation of serial grooming chains by descending decerebration. Behavioural brain research 33, 241–253 (1989).

[89] Berntson, G. G., Jang, J. F. & Ronca, A. E. Brainstem systems and grooming behaviors a. Annals of the New York Academy of Sciences 525, 350–362 (1988).

[90] Hsu, C. T. & Bhandawat, V. Organization of descending neurons in drosophila melanogaster. Scientific reports 6, 20259 (2016).

[91] de Bivort, B. et al. Precise quantification of behavioral individuality from 80 million decisions across 183,000 flies. Frontiers in Behavioral Neuroscience 16, 836626 (2022).

[92] Buchanan, S. M., Kain, J. S. & De Bivort, B. L. Neuronal control of locomotor handedness in drosophila. Proceedings of the National Academy of Sciences 112, 6700–6705 (2015).

[93] Honegger, K. S., Smith, M. A.-Y., Churgin, M. A., Turner, G. C. & de Bivort, B. L. Idiosyncratic neural coding and neuromodulation of olfactory individuality in drosophila. Proceedings of the National Academy of Sciences 117, 23292–23297 (2020).

[94] Kain, J. S. et al. Variability in thermal and phototactic preferences in drosophila may reflect an adaptive bet-hedging strategy. Evolution 69, 3171–3185 (2015).

[95] Skutt-Kakaria, K., Reimers, P., Currier, T. A., Werkhoven, Z. & de Bivort, B. L. A neural circuit basis for context-modulation of individual locomotor behavior. BioRxiv (2019).

[96] Churgin, M. A. et al. Neural correlates of individual odor preference in drosophila. eLife (2023).

[97] Jenett, A. et al. A gal4-driver line resource for drosophila neurobiology. Cell reports 2, 991–1001 (2012).

[98] Nern, A., Pfeiffer, B. D. & Rubin, G. M. Optimized tools for multicolor stochastic labeling reveal diverse stereotyped cell arrangements in the fly visual system. Proceedings of the National Academy of Sciences 112 (2015).

[99] Braun, J., Hurtak, F., Wang-Chen, S. & Ramdya, P. Descending networks transform command signals into population motor control. Nature 1–9 (2024).

[100] Schindelin, J. et al. Fiji: an open-source platform for biological-image analysis. Nature methods 9, 676–682 (2012).

[101] Bradski, G. The OpenCV Library. Dr. Dobb’s Journal: Software Tools for the Professional Programmer 25, 120–123 (2000).

[102] Pagnon, D., Domalain, M. & Reveret, L. Pose2sim: An open-source python package for multiview markerless kinematics. Journal of Open Source Software 7, 4362 (2022).

[103] Werling, K. et al. Addbiomechanics: Automating model scaling, inverse kinematics, and inverse dynamics from human motion data through sequential optimization. Plos one 18, e0295152 (2023).

[104] Begon, M., Andersen, M. S. & Dumas, R. Multibody Kinematics Optimization for the Estimation of Upper and Lower Limb Human Joint Kinematics: A Systematized Methodological Review. Journal of Biomechanical Engineering 140 (2018).

[105] Kim, C., Kim, D. & Oh, Y. Solving an inverse kinematics problem for a humanoid robot’s imitation of human motions using optimization. In International conference on informatics in control, automation and robotics (2005).

[106] Manceron, P. IKPy (2022). URL https://zenodo.org/record/6551158.

[107] Virtanen, P. et al. SciPy 1.0: Fundamental Algorithms for Scientific Computing in Python. Nature Methods 17, 261–272 (2020).

[108] Bohnslav, J. P. et al. DeepEthogram, a machine learning pipeline for supervised behavior classification from raw pixels. eLife 10 (2021).

[109] Hagberg, A., Swart, P. J. & Schult, D. A. Exploring network structure, dynamics, and function using networkx. Tech. Rep., Los Alamos National Laboratory (LANL), Los Alamos, NM, United States (2008).

[110] Todorov, E., Erez, T. & Tassa, Y. MuJoCo: A physics engine for model-based control. In 2012 IEEE/RSJ International Conference on Intelligent Robots and Systems, 5026–5033 (2012). ISSN: 2153-0866.

[111] Marin, E. C. et al. Systematic annotation of a complete adult male drosophila nerve cord connectome reveals principles of functional organisation. eLife (2024).

[112] Fenk, L. M. et al. Muscles that move the retina augment compound eye vision in drosophila. Nature 612, 116–122 (2022).

[113] Stürner, T. et al. Comparative connectomics of the descending and ascending neurons of the drosophila nervous system: stereotypy and sexual dimorphism. bioRxiv (2024).

[114] Schlegel, P. et al. Information flow, cell types and stereotypy in a full olfactory connectome. eLife 10, e66018 (2021).

[115] Yorozu, S. et al. Distinct sensory representations of wind and near-field sound in the Drosophila brain. Nature 458, 201–205 (2009).

[116] Ester, M., Kriegel, H.-P., Sander, J., Xu, X. et al. A density-based algorithm for discovering clusters in large spatial databases with noise. In kdd, vol. 96, 226–231 (1996).

[117] Liu, W. W. & Wilson, R. I. Glutamate is an inhibitory neurotransmitter in the drosophila olfactory system. Proceedings of the National Academy of Sciences 110, 10294–10299 (2013).

[118] Reddi, S. J., Kale, S. & Kumar, S. On the convergence of adam and beyond. arXiv preprint 1904.09237 (2019).

[119] Simes, R. J. An improved bonferroni procedure for multiple tests of significance. Biometrika 73, 751–754 (1986).

